# Molecular fingerprints of a convergent mechanism orchestrating diverse ligand recognition and species-specific pharmacology at the complement anaphylatoxin receptors

**DOI:** 10.1101/2025.05.26.656101

**Authors:** Sudha Mishra, Manish K. Yadav, Annu Dalal, Manisankar Ganguly, Ravi Yadav, Kazuhiro Sawada, Divyanshu Tiwari, Nabarun Roy, Nilanjana Banerjee, Jenny N. Fung, Jianina Marallag, Cedric S. Cui, Xaria X. Li, John D. Lee, Calvin Aaron Dsouza, Shirsha Saha, Parishmita Sarma, Ganita Rawat, Houming Zhu, Htet A. Khant, Richard J. Clark, Fumiya K. Sano, Ramanuj Banerjee, Trent M. Woodruff, Osamu Nureki, Cornelius Gati, Arun K. Shukla

**Author notes:** Equal contribution (Sudha Mishra, Manish K. Yadav, Annu Dalal, Manisankar Ganguly, Ravi Yadav, Kazuhiro Sawada).

## Abstract

Complement anaphylatoxin receptors (C3aR and C5aR1) are prototypical G protein-coupled receptors (GPCRs) playing crucial physiological roles in innate immunity by combating pathogenic infections and orchestrating inflammatory responses. They continue to be important therapeutic targets for multiple disorders including autoimmune diseases, acute and chronic inflammation, and allergy-related conditions. Recent structural coverage has provided important insights into their activation and signaling, however, confounding observations in the literature related to ligand efficacy and functional responses, especially in different model systems, present a major challenge for drug discovery efforts. Here, we systematically and comprehensively profile a broad set of natural and synthetic ligands at C3aR and C5aR1 and discover a previously unanticipated level of functional specialization in terms of species-specific pharmacology and receptor activation. Taking a lead from this, we determine seventeen cryo-EM structures of different ligand-receptor-G-protein complexes and uncover distinct orientation of agonists between the human and mouse receptors despite an overlapping positioning in the orthosteric binding pocket. Combined with extensive mutagenesis and functional assays, these structural snapshots allow us to decode and validate a convergent molecular mechanism involving a “Five-Point-Switch” in these receptors that orchestrates the recognition and efficacy of diverse agonists. We also identify species-specific differences at the level of phosphorylation patterns encoded in the carboxyl-terminus of these receptors and directly visualize their impact on βarr binding and activation using cryo-EM structures. Interestingly, we observe that βarrs engage with the mouse C5aR1 using a variation of previously discovered P-X-P-P phosphorylation motif via a “Sliding-Mechanism” and also exhibit distinct oligomeric state for the human vs. mouse receptors. Taken together, this study elucidates functional specialization at the complement anaphylatoxin receptors and underlying molecular mechanisms, offering a previously lacking framework with direct and immediate implications for the development of novel therapeutics.

## Introduction

The complement system is an integral part of the innate immune response mechanism that is critical for protecting the host against pathogenic infections including bacteria, viruses and fungi^1, 2^. It involves the activation of a multi-step cascade resulting in the generation of a series of peptides in blood plasma with specialized functions coordinating to neutralize pathogens^3^. Complement proteins mediate and contribute towards the process of opsonization, formation of the membrane attack complex (MAC), and recruitment of immune cells to the site of infection triggering inflammation^4^. The terminal products of complement cascade include complement anaphylatoxins, namely C3a, C4a, and C5a, which are produced by the proteolytic cleavage of C3, C4, and C5 complement proteins^1, 5^. Of these, C3a and C5a exert their cellular and physiological functions such as chemotaxis, degranulation and cytokine production via distinct complement anaphylatoxin receptors namely C3aR, C5aR1, and C5aR2^6–9^. These are integral membrane proteins that are classified as the members of the G protein-coupled receptor (GPCR) superfamily, although C5aR2 lacks functional G-protein-coupling upon activation^10–13^. These receptors are expressed at the surface of a variety of immune cells including mast cells, neutrophils, and macrophages, and their downstream signaling via heterotrimeric G-proteins and β-arrestins (βarrs) is crucial for the physiological responses exerted by the anaphylatoxins^14–16^. C3aR and C5aR1 continue to be important therapeutic targets for acute anaphylaxis, sepsis and lung injury, autoimmune disorders such as vasculitis, rheumatoid arthritis, lupus, and multiple sclerosis, as well as chronic inflammatory conditions such as inflammatory bowel disease and neurodegenerative disease^17–20^.

Physiologically, C3a and C5a are rapidly proteolyzed by carboxypeptidases to remove the terminal arginine residue leading to the generation of C3a^-d-Arg^ and C5a^-d-Arg^, respectively^21, 22^. Such a mechanism is generally believed to help dampen the inflammatory response exerted by C3a and C5a, as C3a^-d-Arg^/C5a^-d-Arg^ are typically believed to lose their ability to bind and activate their cognate receptors^21–26^. However, a few reports have suggested that a complete loss of receptor-binding and activation abilities of C3a^-d-Arg^/C5a^-d-Arg^, as originally conceived, may not be accurate and requires systematic exploration^27^. More recently, a naturally-occurring peptide derived from VGF, the large neuropeptide expressed in peripheral and central nervous system, known as TLQP21 has been identified as the second natural agonist of C3aR with a functional contribution in lipolysis, neurodegeneration, and metabolism^28–31^. In particular, TLQP21-induced, β-adrenergic receptor-mediated lipolysis in adipocytes, and increased resting energy expenditure in mouse model, has uncovered untapped therapeutic potential of C3aR in metabolic conditions including obesity^30–33^. Moreover, TLQP21-mediated activation of microglial BV2 cells promotes the clearance of extracellular amyloid-β fibrils. Additionally, the peptide modulates microglial function through C3aR signaling pathways, reducing neuropathology in mouse models^34, 35^. However, an incomplete understanding of species-specific functional responses encoded by the TLQP21-C3aR axis poses a key challenge in therapeutic development^29, 36, 37^.

Although peptides derived and modified from the carboxyl-terminus of C3a and C5a are known to behave as partial agonists at these receptors with potential therapeutic opportunities^38–41^, a systematic study to decipher the intricate details of receptor subtype selectivity, species-specificity, and signaling-bias is still lacking. For example, EP67, a peptide derived from the carboxyl-terminus of human C5a, has shown promising indications not only as a vaccine adjuvant^42–47^ but also to overcome the burden of pathogenic infections, especially in the context of anti-microbial resistance^48, 49^ and biofilm formation^50^. Still however, there are contradictory reports in the literature about its pharmacology and primary receptor that it engages to elicit the functional responses^48, 49^. Finally, a handful of small molecule ligands have been described for C3aR, including the well-characterized C3aR antagonists SB290157 and JR14a^51–55^. However, emerging evidence from multiple studies has hinted at their agonistic properties and species-specific pharmacology^36, 55–58^, which remains primarily unexplored at the molecular and structural levels, although JR14a-bound C3aR structures have been reported recently^58, 59^. Taken together, C3aR and C5aR1 constitute a pair of closely related receptors with a rich tapestry of ligands and unparalleled opportunity to explore the molecular basis of diverse ligand recognition, species-specific pharmacology, and signaling-bias.

In this backdrop, here we present a comprehensive profiling of a diverse set of ligands at C3aR and C5aR1 to uncover previously unknown species-specific pharmacology of the natural and synthetic agonists as well as naturally-encoded signaling-bias. We discover that contrary to the generally believed notion, C5a^-d-Arg^ maintains G-protein-coupling while displays an attenuated βarr recruitment, which is further reflected in differential responses in terms of cytokine release and neutrophil migration in primary cells and mouse model as presented in the accompanying manuscript. Using cryo-EM structures of distinct agonist-receptor-transducer combinations, we rationalize the agonistic efficacy of diverse ligands including those previously portrayed as antagonists and discover a key switch in the orthosteric binding pocket that is engaged to impart robust agonism. Finally, we also uncover distinct modalities of βarr oligomerization and phosphorylation motif engagement for the human vs. mouse receptors, which further expands the framework of species-specific pharmacology to species-specific transducer-coupling. This study provides the most comprehensive analysis of the complement anaphylatoxin receptor pharmacology and molecular mechanisms to date, with direct implications for novel drug discovery targeting these receptors.

## Results

### Species-specific pharmacology and signaling-bias at C3aR and C5aR1

We first set out to comprehensively profile a broad set of ligands at C3aR and C5aR1 including the natural agonists C3a and C5a, C3a^-d-Arg^ and C5a^-d-Arg^, and TLQP21, synthetic peptide agonist EP54 and EP67, and small molecule ligands SB290157 and JR14a (**Figure 1A-D**). We tested all possible combinations in terms of ligands of human and mouse origin on human and mouse C3aR and C5aR1 in transducer-coupling assays namely cAMP inhibition, G-protein dissociation and βarr1/2 recruitment using GloSensor and NanoBiT assays **(Figure 1E-T and S1A-N, and S3)**^60–62^. We observed that human and mouse C3a exhibited a modest but significant difference in potency at the human C3aR, with human C3a being more potent, although both were full agonists (**Figure 1E and S1A-B**). On the other hand, both human and mouse C3a were equally potent and efficacious at the mouse C3aR (**Figure 1F and S1A-B**). In contrast, human and mouse C5a displayed similar potency and efficacy at the human and mouse C5aR1 (**Figure 1G-H and S1C-D**).

**Fig. 1:**
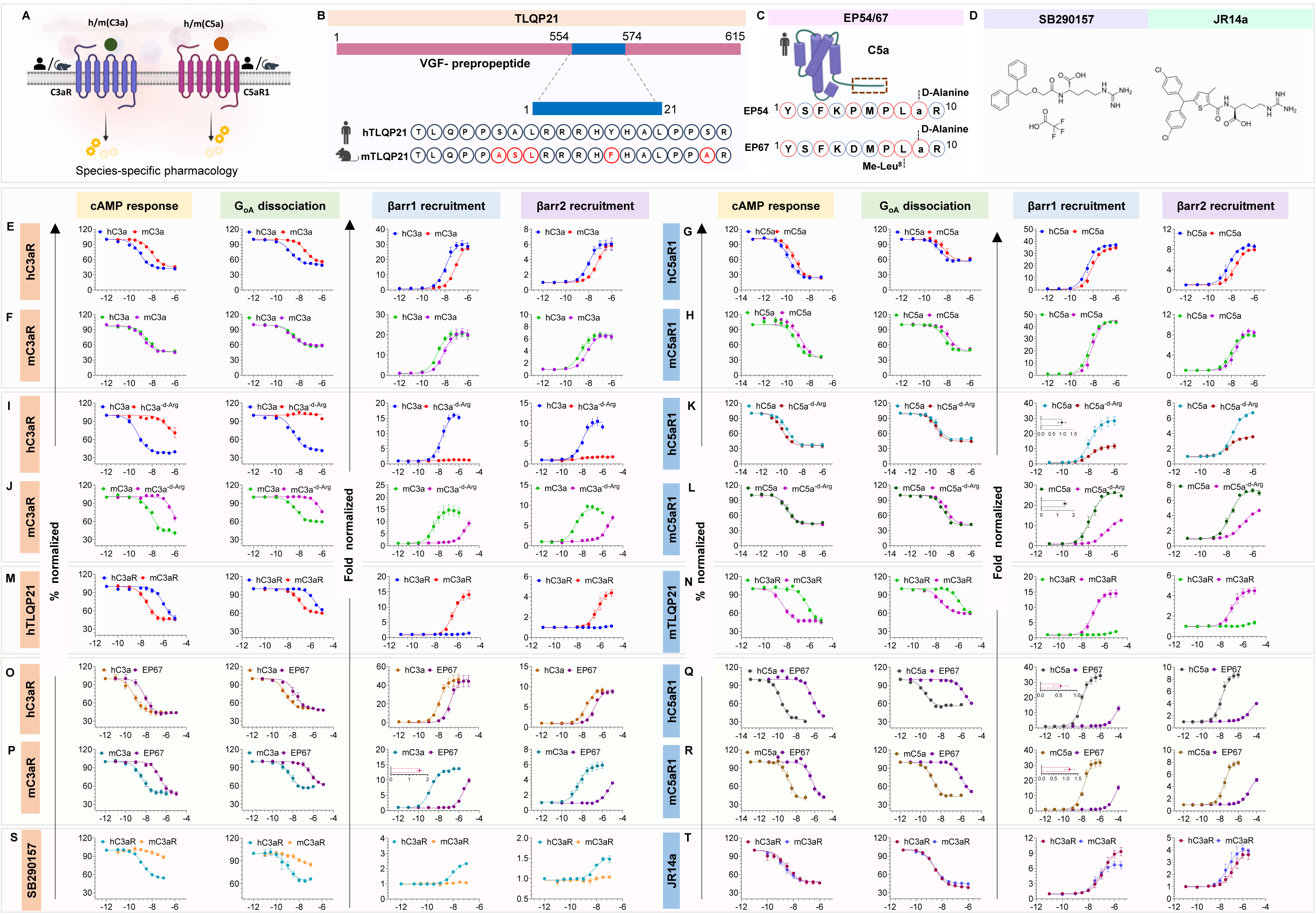
Pharmacological profiling of human and mouse anaphylatoxin receptors C3aR and C5aR1. (A) Schematic illustrating species-specific signaling at C3aR and C5aR1. (B) Overview of the VGF-derived neuropeptide TLQP21, which regulates physiological processes such as lipid metabolism and energy homeostasis. A comparative sequence analysis of human (h) and mouse (m) forms is presented. The red circle indicated the amino acid difference between h and mTLQP-21. (C) Illustration of EP54 and EP67, C5a-derived and modified peptides. The red circle indicates the amino acid difference from hC5a. (D) Chemical structures of SB290157 and JR14a. (E–T) Pharmacological profiling reveals distinct ligand-induced responses at h and mC3aR and C5aR1. Gαi activation was quantified using the GloSensor assay by measuring ligand-mediated inhibition of forskolin-induced cAMP production (mean ± SEM, n = 3). G-protein activation was further validated using the NanoBiT-based GαoA dissociation assay, with responses normalized to the lowest ligand concentration. βarr1/2 recruitment was assessed via a NanoBiT luminescence assay, and ligand-induced responses were normalized with the lowest ligand concentration set to 1. Bias factors for C5a^-d-Arg^ and EP67 were calculated relative to the reference ligand h/mC5a and mC3a using the online bias calculator tool (https://biasedcalculator.shinyapps.io/calc/), based on efficacy and potency data from Gi activation and βarr recruitment assays. (E–H) Comparative pharmacological analysis indicates species-specific differences in C3a-induced signaling at C3aR, while C5aR1-mediated responses remain conserved. (I and J) Pharmacological profiling of C3a^-d-Arg^ showed no detectable response at hC3aR but exhibited a minor effect at mC3aR. (K and L) Pharmacological profiling identifies C5a^-d-Arg^ as a G-protein-biased ligand, maintaining full G-protein response similar to C5a while βarrs recruitment is comparatively lower. (M and N) TLQP21 functional responses downstream of h and mC3aR uncovered its selective activation towards mC3aR. It functions as a biased ligand for mC3aR even at equal receptor surface expression. (O–R) Species-specific pharmacological profiling of EP67 demonstrated higher potency at hC3aR compared to mC3aR and C5aR1 complement receptors. (S and T) SB290157 acts as a species-specific agonist at hC3aR, with no measurable agonistic response observed at mC3aR. JR14a activates both h and mC3aR and functions as a G-protein-biased agonist.

Both C3a and C5a undergo proteolytic cleavage under physiological conditions resulting in the removal of terminal arginine and generation of C3a^-d-Arg^ and C5a^-d-Arg^, respectively, and this is generally believed as a mechanism to dampen the inflammatory response to limit prolonged inflammation^21, 22^. Accordingly, we observed near-complete loss of transducer-coupling for human C3a^-d-Arg^ on the human and mouse C3aR, however, mouse C3a^-d-Arg^ behaved as a weak partial agonist on the mouse C3aR (**Figure 1I-J**). Surprisingly, we observed that both human and mouse C5a^-d-Arg^ maintained equal potency and efficacy on C5aR1 in G-protein coupling and activation but displayed dramatically attenuated responses in βarr recruitment assays (**Figure 1K-L and S1E-H**). In fact, a previous study has suggested that C5a^-d-Arg^ is nearly as potent and efficacious as C5a in activating the rat basophil leukemia (RBL) and human polymorphonuclear cells as assessed using an in-vitro label-free biosensor assay^63^. Moreover, the attenuated βarr recruitment by C5a^-d-Arg^ is also reflected in compromised receptor endocytosis compared to C5a (**Figure S1G-H**), and a differential response in terms of neutrophil migration *in-vitro* and *in-vivo* as presented in the accompanying manuscript. Taken together, these data establish the C5a^-d-Arg^ as a naturally-encoded, G-protein-biased agonist at C5aR1, in contrast to the prevalent perception in the literature that it loses the ability to bind and activate C5aR1.

In addition to C3a as the natural agonist, C3aR also recognizes another naturally-encoded peptide known as TLQP21, which is derived from VGF, a member of the extended granin family with primary expression in the central and peripheral nervous system^28, 29, 64^. Interestingly, TLQP21 is reported to promote energy expenditure, and modulate inflammatory pain and gastric contractility in mouse models^65^, and also inhibit feeding in hamsters^66, 67^, and therefore, the TLQP21-C3aR axis has been gaining traction as a new therapeutic target in obesity and metabolic disorders^68^. Previous studies have established the selectivity of TLQP21 for C3aR over C5aR1^33, 64^, and our functional profiling data presented here revealed that both human and mouse TLQP21 are significantly more potent on the mouse C3aR compared to the human C3aR (**Figure 1M-N**). More interestingly, both human and mouse TLQP21 display a very low level of βarr recruitment on the human C3aR while they are nearly as effective as C3a on the mouse C3aR (**Figure 1M-N and S1I-J)**. These findings underscore the species-specific pharmacology exhibited by TLQP21 and naturally-encoded signaling-bias at C3aR, making it an interesting system for structural analysis.

In addition to the natural-derived peptide agonists of C3aR and C5aR1, several synthetic peptides derived and modified based on the carboxyl-terminus of C3a and C5a have also been explored in terms of their agonism and signaling-bias and receptor selectivity^38, 69–73^. Of these, EP54, a decapeptide designed based on the carboxyl-terminal sequence of human C5a, and EP67, an improved version of EP54, have also been characterized as an effective vaccine adjuvant^42–47^, to facilitate immune response during pathogenic infections such as drug-resistant strains of *Staphylococcus aureus*^48, 49^. However, there are confounding reports in the literature at the level of specific complement anaphylatoxin receptor that it engages to elicit the functional responses^74^. Therefore, we first profiled EP54 and EP67 on the human and mouse C3aR and C5aR1 in the transducer-coupling assays. We observed that EP67 was almost as potent and efficacious as C3a on the human C3aR in G-protein and βarr assays (**Figure 1O**). On the other hand, it displayed a weaker potency but nearly full efficacy in G-protein assays at mouse C3aR while displayed a weak partial agonism in βarr recruitment (**Figure 1P**). More strikingly, EP67 displays a significantly weaker potency at the human and mouse C5aR1 in the G-protein assay although its efficacy is similar to C5a, however, it exhibits a substantially attenuated βarr recruitment on both human and mouse C5aR1 (**Figure 1Q-R and S1K-L**). This observation potentially underscores the importance of engaging the receptor N-terminus for imparting high-affinity and full agonism at C5aR1, while being relatively less crucial for C3aR. On the other hand, EP54 exhibited weaker potency but nearly-full efficacy at C3aR and C5aR1 in G-protein and βarr assays although it was slightly less efficacious at mouse C3aR (**Figure S3A-I)**.

Finally, a series of small molecule ligands portrayed as antagonists have been developed and characterized at C3aR^51, 52^. However, the most well studied compounds, namely SB290157 and JR14a, have also been reported to exhibit agonistic activity in various cellular assays^56–58^. Therefore, we profiled both molecules in our transducer-coupling assays on the human and mouse C3aR, and observed that both elicited agonistic efficacies, with SB290157 being more potent than JR14a but less efficacious (**Figure 1S-T and S1M-N**). Strikingly, SB290157 exhibited a selective activation of the human C3aR but not the mouse C3aR, while JR14a was nearly identical at both human and mouse C3aR (**Figure 1S-T**). In comparison to C3a, SB290157 was equally potent but much less efficacious, while JR14a was less potent but equally efficacious at human C3aR (**Figure S1M**). These data suggest that similar to naturally-occurring peptide agonists i.e., TLQP21, the small molecule compound SB290157 also encodes species-specific pharmacology at C3aR. The receptor expression as measured using cell surface ELISA in each of the functional assays are presented in **Figure S2 and S3**.

Taken together, comprehensive profiling of ligands on C3aR and C5aR1 uncovered modest but significant species-specific differences in anaphylatoxin pharmacology, and near-complete loss of function of C3a^-d-Arg^ but unexpectedly, strong G-protein-bias of C5a^-d-Arg^. We also revealed a striking pattern of selectivity and G-protein-bias of the human and mouse TLQP21 at C3aR, preferential activation of human C3aR by C5a-derived EP67 compared to C5aR1, and robust agonism of SB290157 and JR14a at C3aR with striking selectivity of SB290157 for hC3aR vs. mC3aR.

### Structural coverage of ligand receptor complexes

In order to understand the molecular basis of the intriguing functional responses exhibited by these diverse set of ligands, we next determined seventeen structures of the human and mouse C3aR and C5aR1 in complex with various agonists and heterotrimeric G-proteins using cryo-EM (**Figure 2A-Q**). The experimental details of complex reconstitution, cryo-EM data collection and processing, and overall regions of different components resolved in the structures are summarized in **Figure S4-5 and S7-10 and Table S1 and S2**. As anticipated, the overall structures of the complexes of respective receptors with different ligands are similar in terms of receptor conformation with an RMSD of <1Å. The unambiguous density corresponding to the cryo-EM map allowed us to model all the components of the complex including the majority of side chain residues. In the mouse C3a-bound mouse C3aR structure, the density corresponding to the core domain of the mouse C3a was poorly resolved in the final map, most probably due to flexibility and lack of stable interaction with the receptor, still however, the distal carboxyl-terminus was well resolved to occupy the orthosteric ligand binding pocket in the receptor (**Figure 2A and S11A**). In addition, the density corresponding to the proximal stretch of the N-terminus of C3aR, and majority of the exceptionally long 2^nd^ extracellular loop (ECL2) i.e., from Ser^178^ to His^321^ were also not observed. For the human and mouse TLQP21 in complex with the mouse C3aR, the proximal stretch of the ligand was not resolved, however, the distal segment ranging from R13 to R21 occupying the orthosteric binding pocket was resolved well (**Figure 2B-C, S11B-C)**. The density for the small molecule ligands SB290157 and JR14a in the orthosteric binding pocket of C3aR was clearly observed allowing the modelling of the ligands and clear identification of the differences between the two ligands in terms of their interaction with the receptor (**Figure 2F-H, S13A-C)**. Finally, the densities of mouse C5a on h/mC5aR1, human and mouse C5a^-d-Arg^ on the mouse C5aR1 were clearly resolved and allowed the visualization of receptor N-terminus engagement in addition to the orthosteric binding pocket (**Figure 2J-M and S14A-C, S15A)**. The interaction of EP67 on both human and mouse C3aR and C5aR1, as well as that of EP54 on mC3aR and hC5aR1, were clearly visible with discernible interaction with the receptors (**Figure 2D-E, N-O, S12A-B and S15B-C)**, thereby allowing a direct comparison across these receptors. Taken together with previously reported structures of the C3aR and C5aR1^75–77^, these structures offer a rich tapestry for extracting structure-guided insights into receptor pharmacology and signaling-bias.

**Figure 2.**
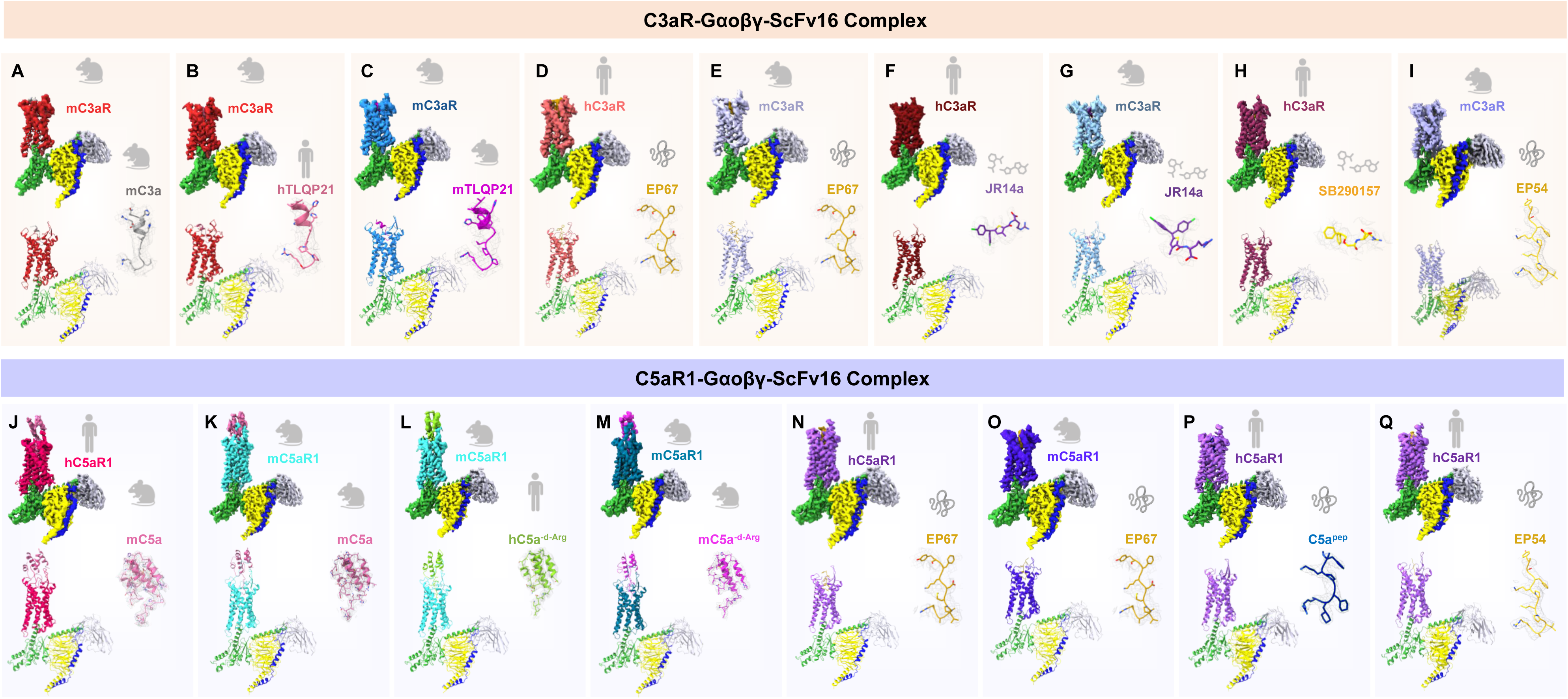
Cryo-EM structures of C3aR-Gαo and C5aR1-Gαo complexes. (A-Q) EM maps and models corresponding to the mC3a-mC3aR, hTLQP21-mC3aR, mTLQP21-mC3aR, EP67-hC3aR, EP67-mC3aR, JR14a-hC3aR, JR14a-mC3aR, SB290157-hC3aR, EP54-mC3aR, mC5a-hC5aR1, mC5a-mC5aR1, hC5a^-d-Arg^-mC5aR1, mC5a^-d-Arg^-mC5aR1, EP67-hC5aR1, EP67-mC5aR1, C5a^pep^-hC5aR1, and EP54-hC5aR1 complexes respectively. Cryo-EM density maps of respective ligands have been shown next to the receptor density. Lime green, Gαo; yellow, Gβ1; dark-blue, Gγ2; grey, ScFv16.)

### A convergent mechanism of diverse ligand recognition and receptor activation

Binding of complement anaphylatoxins C3a and C5a to C3aR and C5aR1, respectively, has been conceived to utilize a two-site mechanism involving the N-terminus of the receptor referred to as “site 1”, and the extracellular side of the transmembrane bundle referred to as “site 2” or orthosteric binding pocket^73^. While C5a binding to C5aR1 has been directly visualized to involve an interface involving the N-terminus, direct involvement of the N-terminus of C3aR in C3a binding has not been observed in the cryo-EM structures^75, 76, 78^. Structural comparison of C3aR and C5aR1 structures determined here reveals that the distal carboxyl-terminus of the human and mouse C3a/C5a penetrate in the orthosteric binding pocket, and the carboxyl-terminal residues make multiple contacts in the orthosteric binding pocket including Ser^2.60^, Ile^3.32^, Arg^5.42^, Tyr^6.51^, and Asp^7.35^ in C3aR, and Leu^2.60^, Leu^3.32^, Arg^4.64^, Tyr^6.51^, Asp/Asn^7.35^ in C5aR1 (**Figure 3A-T S17, S19 and Table S3-S4**). In particular, the terminal arginine (R77/R78 in h/mC3a and R74 in C5a) is involved in strong ionic or cation-π interaction with at least two of these five residues (**Figure S17A-F**). Interestingly, the carboxyl-terminus of C5a^-d-Arg^ slides in the orthosteric binding pocket in a manner that Q71, L72, and G73 interact with the same five residues mentioned above and thereby compensate for the missing R74 (**Figure 3M-O and S17G-I**). Moreover, the carboxyl-terminal stretch of TLQP21 and EP67 also occupy a similar and nearly-overlapping binding pocket on the corresponding receptors, and their terminal arginine residue also engages a similar set of residues (**Figure 3B-E, and 3P-Q, Figure S17K-L and S18A-B**). Interestingly, a closer inspection of SB290157-C3aR and JR14a-C3aR structures reveal that these ligands also occupy nearly the same orthosteric pocket on C3aR and exhibit similar interactions as other agonists (**Figure 3F-H and S18C-I**). For example, the arginine moiety of SB290157 and JR14a occupy a similar position as that of R77 in C3a with the guanidino group engaging Asp^7.35^ and Tyr^6.51^ through hydrogen bond and cation-π interaction (**Figure 3F-H and S18D-I**). In addition, the terminal oxygen of these ligands interacts with Tyr174^ECL2^ through a non-bonded contact (**Figure S18D-I**), which is also corroborated by the recently reported JR14a-hC3aR structures^58, 59^.

**Figure 3.**
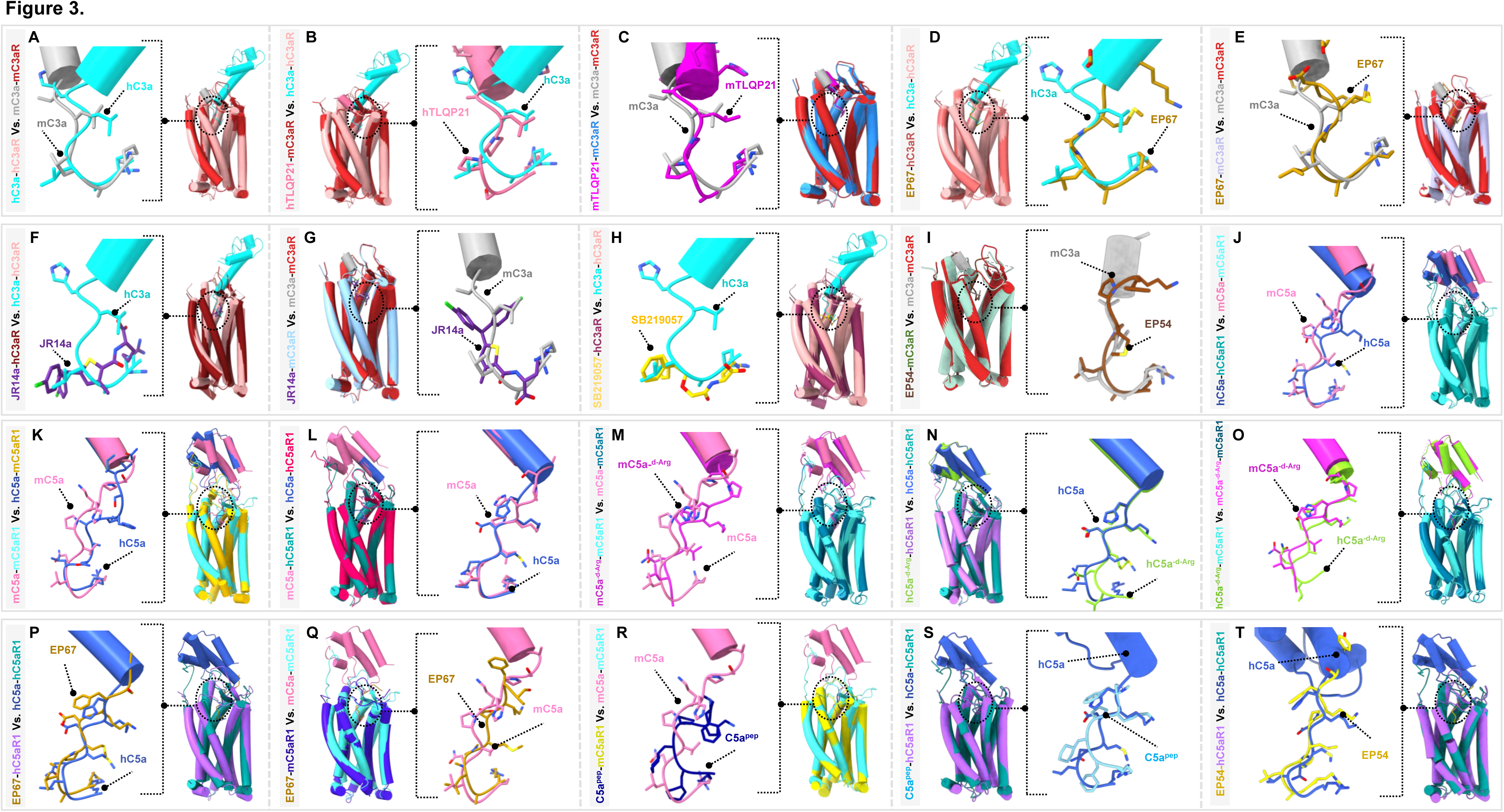
Distinctive binding pose of the ligands in complement receptors. (A) hC3a on hC3aR and mC3a on mC3aR; (B) hTLQP21 on mC3aR and hC3a on hC3aR; (C) mTLQP21 on mC3aR and mC3a on mC3aR; (D) EP67 on hC3aR and hC3a on hC3aR (E) EP67 on mC3aR and mC3a on mC3aR; (F) JR14a on hC3aR and hC3a on hC3aR; (G) JR14a on mC3aR and mC3a on mC3aR; (H) SB290157 on hC3aR and hC3a on hC3aR; (I) EP54 on mC3aR and mC3a on mC3aR; (J) hC5a on hC5aR1 and mC5a on mC5aR1; (K) mC5a on mC5aR1 and hC5a on mC5aR1; (L) mC5a on hC5aR1 and hC5a on hC5aR1; (M) mC5a^-d-Arg^ on mC5aR1 and mC5a on mC5aR1; (N) hC5a^d-Arg^ on hC5aR1 and hC5a on hC5aR1; (O) hC5a^-d-Arg^ on mC5aR1 and mC5a^-d-Arg^ on mC5aR1; (P) EP67 on hC5aR1 and hC5a on hC5aR1; (Q) EP67 on mC5aR1 and mC5a on mC5aR1; (R) C5a^pep^ on mC5aR1 and mC5a on mC5aR1; (S) C5a^pep^ on hC5aR1 and hC5a on hC5aR1; (T) EP54 on hC5aR1 and hC5a on hC5aR1 into the orthosteric pocket of complement receptors.

Taken together, these observations suggest a convergent mechanism of diverse ligand recognition driving agonist efficacy at C3aR and C5aR1, where nearly every single agonist visualized so far, engages a set of five common residues in C3aR and C5aR1. These residues namely Ser/Leu^2.60^, Ile^3.32^/Leu^3.33^, Arg^5.42^/Arg^4.64^, Tyr^6.51^, and Asp/Asn^7.35^ are positioned in a manner to link together TM2-7 in the orthosteric binding pocket, and we referred to them as the “Five-Point-Switch” (**Figure 4A-B)**. Of these, Tyr^6.51^ and Asp/Asn^7.35^ appear to be the most crucial and conserved in the orthosteric binding pocket. In line with structural observations, site-directed mutagenesis of Tyr^6.51^ and Asp/Asn^7.35^ in the human and mouse C3aR and C5aR1 resulted in a dramatic loss of both cAMP response and βarr recruitment further validating a predominant role of these residues in ligand recognition and receptor activation (**Figure 4C-F**). Expectedly, there are additional ligand-specific contacts visualized in the structure, which are likely to drive further fine-tuning of the functional responses including signaling-bias and species-specific functional responses as discussed in the following sections.

**Figure 4.**
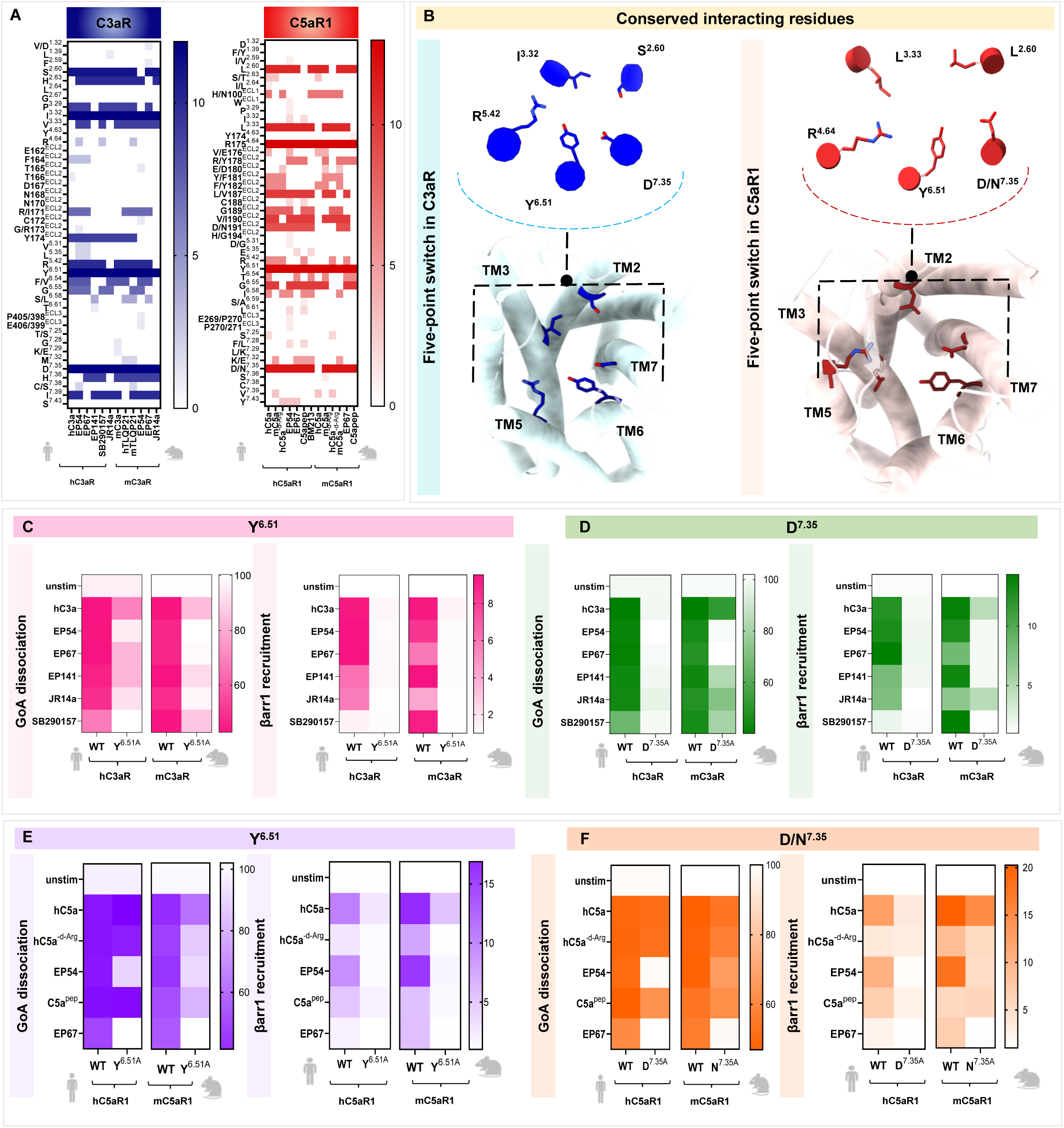
C3aR and C5aR1 exhibit a conserved activation mechanism upon activation with various ligands. (A) The heatmap illustrates common interacting residues among hC3aR, mC3aR, hC5aR1, and mC5aR1 in complex with the specified ligands. Previously reported activated receptor structures were also included in the analysis, such as hC3a-hC3aR (PDB: 8I9L), EP54-hC3aR (PDB: 8I95), EP141-hC3aR (PDB: 8J6D), hC5a-hC5aR1 (PDB: 8IA2), hC5a^-d-Arg^-hC5aR1 (PDB: 8JZZ), hC5a-mC5aR1 (PDB: 8HQC), BM213-hC5aR1 (PDB: 7Y66), and C5a^pep^-mC5aR1 (PDB: 8HPT). Residue conservation across different ligand-induced conformations was quantified and visualised using a colour-coded intensity gradient, where higher intensity corresponds to greater conservation of interaction. (B) Highlighting the conserved residues involved in ligand-binding and receptor activation across the indicated ligands, downstream of anaphylatoxin receptor C3aR and C5aR1. (C-F) Heatmap summarising the maximal responses downstream of C3aR/C5aR1-WT and mutant (Y^6.51A^ and D/N^7.35A^) using NanoBiT-based GαoA dissociation assay (where ligand-induced decrease in the luminescence was measured) and β-arrestin1 recruitment (where ligand-induced increase in luminescence was read). The responses for various tested ligands are normalised with unstimulated (set as 100% for GαoA and 1 for β-arrestin1). Data are presented as mean ± SEM (n = 3).

### Structural basis of species-specific pharmacology and alternate agonism at C3aR

Functional profiling suggests nearly identical response of the mouse and human C5a on the human and mouse C5aR1, however, human and mouse C3a show a modest but significant difference in terms of their potency at the human vs. mouse C3aR. Structural analysis of the human and mouse C3a with the corresponding receptors, show an overall similar hook-like conformation in both, which overlaps well in the orthosteric binding pocket (**Figure 5A**). Interestingly however, we observed that the positioning of human and mouse C3a diverge significantly on their respective receptors at an angle of ∼80° when compared with respect to L72 in hC3a as a pivot point (**Figure 5A**). As a result, there is a discernible difference in the interaction pattern of the two ligands on their corresponding receptors as summarized in **Figure 5B**, despite the conserved engagement of the five key residues as mentioned above.

**Figure 5.**
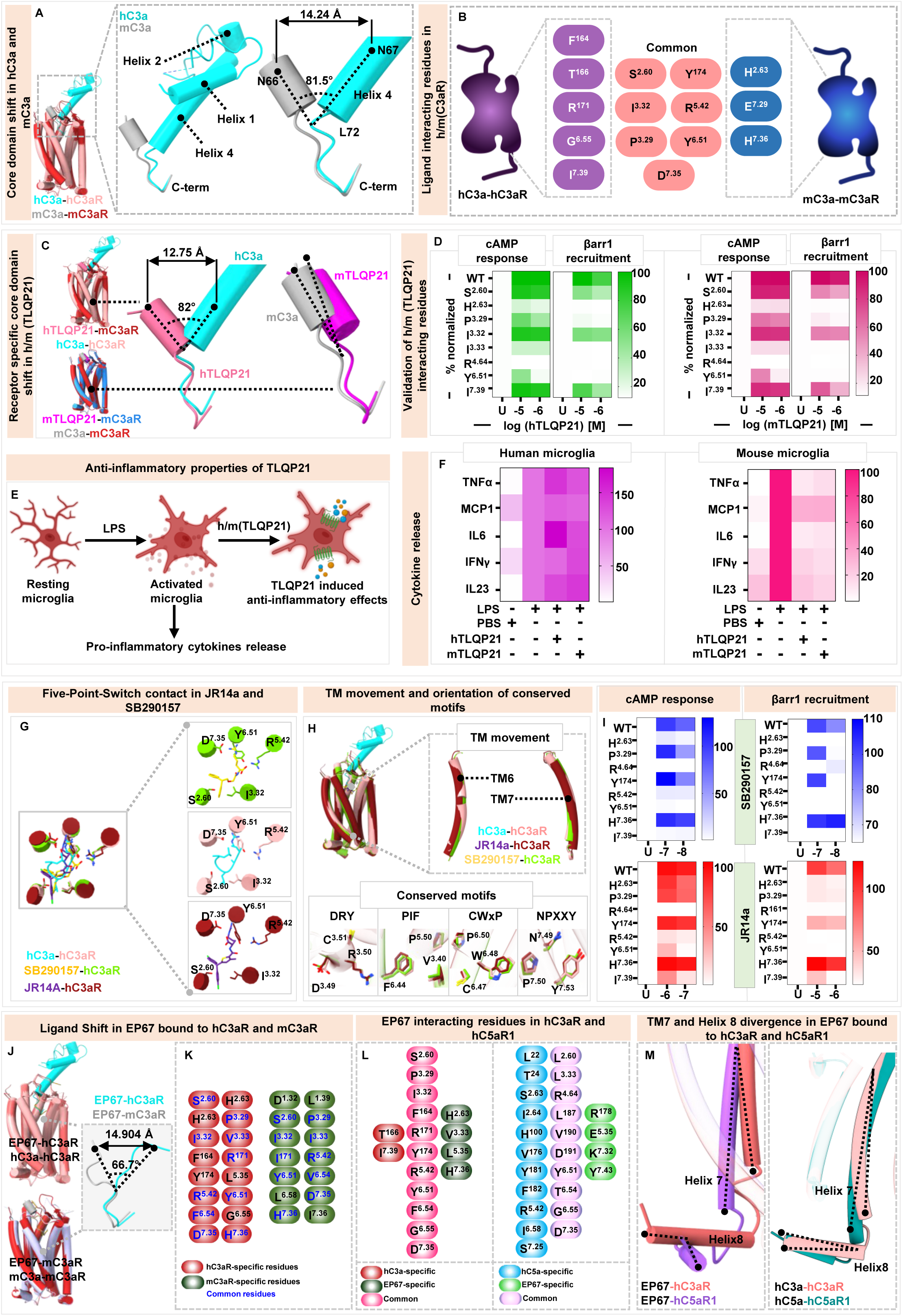
Binding of ligands with C3aR. (A) Structural superimposition of hC3a-hC3aR and mC3a-mC3aR complex structures. Comparison of hC3a and mC3a binding modes on their corresponding receptors reveals major differences in the orientations of the helices of the core domain (*inset*). (B) Interactions of hC3a/mC3a with the orthosteric pocket residues of hC3aR/mC3aR. The schematic representation depicts the residues at the interface of hC3a/mC3a with hC3aR/mC3aR, respectively. (C) Structural superimposition of hTLQP21-mC3aR with hC3a-hC3aR, and mTLQP21-mC3aR with mC3a-mC3aR, respectively. The comparison reveals a receptor-specific, distinct binding pose of the ligand. (D) Structural validation of the ligand-binding site in h/mTLQP21-mC3aR structures was performed by mutating key ligand-interacting residues to alanine. The effects of these mutations were assessed by measuring ligand-induced maximal responses in mC3aR-WT and mutant variants through cAMP response and βarr1 recruitment assays. The results are summarised in a heatmap, with data represented as mean ± SEM (n = 3). The wild-type (WT) receptor response was defined as 100%, and the mutant’s response was normalized against this value (E) Schematic representation of anti-inflammatory effect mediated by h/m TLQP21 on microglial cells. (F) The immunomodulatory effect of hTLQP21 and mTLQP21 on LPS-induced cytokine release in human monocyte-derived microglia-like cells (MDMi) and primary mouse microglia. The ligand-mediated reduction in cytokine levels was assessed, with LPS-induced release set as the 100% baseline. The full panel of cytokine and chemokine release measurement is provided in Figure S20E. (G) Five-Point-Switch contact in JR14a and SB290157. The three critical residues stabilising the conformation of the ligands within the orthosteric pocket of hC3aR are shown. (H) Structural superimposition of JR14a and SB290157-hC3aR with the hC3a-hC3aR structure. The comparison reveals similar TM movements and similar conformational changes in the conserved microswitches upon ligand-mediated activation of hC3aR. (I) Structural validation of the ligand-binding site in SB290157/JR14a-hC3aR structures was conducted by substituting key ligand-interacting residues with alanine. The impact of these mutations was evaluated by assessing ligand-induced maximal responses in hC3aR-WT and mutant variants using cAMP response and βarr1 recruitment assays. The findings are presented in a heatmap, with data expressed as mean ± SEM (n = 3). The wild-type (WT) receptor response was defined as 100%, and the mutant’s response was normalized against this value (J) Receptor-specific binding poses of EP67. Although the C-termini of EP67 align well with each other, a distinct deviation is observed at the proximal site. (K) Common and specific residue interactions between EP67 with hC3aR and mC3aR are shown. (L) Common and specific residue interactions between EP67/hC3a with hC3aR, and EP67/hC5a with hC5aR1 are shown. (M) EP67 induced distinctive helix-8 movement in hC5aR1, which differs from the helix-8 conformations observed in EP67-hC3aR, hC3a-hC3aR, and hC5a-hC5aR1.

Human and mouse TLQP21 display a much weaker response on the human C3aR while robustly activating mouse C3aR. Therefore, we used mouse C3aR activated by human and mouse TLQP21 to investigate their agonism and compare them with the mC3a-mC3aR structure. The binding of TLQP21 primarily involves the orthosteric pocket of the receptor, occupying a buried surface area of ∼1700Å^2^, and the N-terminal segment adopts a helical conformation similar to C3a (**Figure 5C**). Strikingly, human and mouse TLQP21 are also positioned on mC3aR in an orientation that shows a cross-angle of ∼80° compared to human C3a on the human C3aR (**Figure 5C**). Carboxyl-terminal residues adopt a hook-like structure, analogous to C3a, and occupy a similar position in the orthosteric binding pocket in line with a conserved mechanism of agonism (**Figure S17A, K**). As predicted, the overall structural features of C3aR in complex with TLQP21 are similar to that of C3a-bound receptor representing a prototypical active conformation with outward movement of TM6 and 7, and the rearrangement of key structural motifs. Structure-guided mutagenesis of C3aR combined with transducer-coupling assays further validate the key contribution of the interacting residues as visualized in the structures (**Figure 5D and S20A-D**).

In order to confirm the preference of TLQP21 for the mouse C3aR over human C3aR in the native biological context, we measured the effect of TLQP21 stimulation on LPS-induced pro-inflammatory cytokine release from primary human and mouse microglia. These cells express C3aR at endogenous levels^79^, and prior studies support that C3a stimulation can modulate LPS-driven cytokine responses in phagocytic cells^77^. We observed that stimulation of mouse, but not human microglial cells resulted in a robust attenuation of cytokine release upon h/mTLQP21 treatment (**Figure 5E-F and S20E**), confirming the striking preference of TLQP21 for the mouse C3aR in cellular context. Although we do not have the structure of TLQP21 in complex with human C3aR, it is plausible that the divergent positioning of TLQP21 on human vs. mouse C3aR, similar to that observed for C3a, drives this species-specific pharmacology (**Figure 5C**). However, this hypothesis remains to be tested experimentally in future studies, which may also help facilitate the design of new TLQP21 analogs with improved efficacy at the human receptor.

Finally, the analysis of SB290157-bound human C3aR and JR14a-bound human and mouse C3aR structures provides the molecular basis for small molecule agonism at C3aR. These ligands occupy a largely similar orthosteric binding pocket on the receptor, thereby engaging the critical interactions as identified earlier for other agonists (**Figure 5G**). However, unlike peptide agonists, JR14a does not engage Arg^5.42^ in either human or mouse C3aR, whereas SB290157 is able to interact with this residue (**Figure S18D-I**). In addition, the overall structure features of C3aR in complex with these ligands are nearly identical to that of C3a-bound receptor representing a prototypical active conformation with outward movement of TM6 and 7, and the rearrangement of structural motifs (**Figure 5H**). Structure-guided mutagenesis of C3aR combined with transducer-coupling assays further validates the key interaction network of these two ligands (**Figure 5I and S20F-G**). As noted in Figure 1, SB290157 displayed striking selectivity for the human C3aR while JR14a is able to recognize both human and mouse receptors. Although the overall structures of SB290157- and JR14a-bound C3aR are very similar, the spatial positioning of the carboxylate group in SB290157 is marginally deeper in the orthosteric pocket compared to that of JR14a (**Figure S20H**). Moreover, while the overall interaction network of JR14a with human and the mouse receptor is largely conserved within the orthosteric binding pocket, some notable differences are also observed. Specifically, His^7.36^ engages in a π-π interaction with one of the 4-chlorotoluene rings in the JR14a-bound mouse C3aR structure, which is missing in the human counterpart (**Figure S20I)**. Thus, it is possible that the lack of 4-chlorotoluene rings in SB290157 drives its preference for the human C3aR over mouse C3aR although further studies are required to probe this hypothesis.

### Structural insights into subtype selectivity of EP67

As we have determined the structures of EP67 in complex with both human and mouse C3aR and C5aR1, it allows us to systematically explore the molecular details of receptor subtype selectivity and efficacy. As mentioned earlier, EP67 is a decapeptide derived and modified from the distal carboxyl-terminus of human C5a, however, there are confounding reports in the literature about its cross-reactivity and selectivity toward C3aR and C5aR1^48, 49^. Our systematic analysis of cellular responses demonstrates that it is capable of activating both C3aR and C5aR1, with a clearly higher potency and efficacy for human C3aR compared to human C5aR1. On the other hand, its efficacy and potency appeared to be relatively weaker for mouse C3aR compared to human C3aR. The structural snapshots determined here reveal that the distal carboxyl terminus of EP67 adopts a hook-like conformation, occupying a similar orthosteric pocket of the receptors, and has a significantly conserved set of interactions as that of C3a and C5a (**Figure 5J-L**). Interestingly, the carboxyl-terminal segment of EP67 aligns well on human and mouse C3aR, however, its N-terminal segment diverges around a proline kink (P7) making an angle of ∼65° between the two receptors (**Figure 5J)**. This may explain the differences in the efficacy and potency of EP67 observed for the human vs. mouse C3aR although future studies are required to test this hypothesis.

On the other hand, comparison of EP67 on human and mouse C5aR1 does not reflect a significant divergence in their positioning on the orthosteric binding pocket (**Figure S21A)**. A direct comparison of the network of interactions between EP67 and C3a/C5a in C3aR and C5aR1, respectively, underscores a relatively broader gap for C5aR1 compared to C3aR (**Figure 5L**). This is in line with the involvement of the C5aR1 N-terminus in the interaction with C5a that is not observed for C3a-C3aR structures. Therefore, it is likely that the missing interactions in the case of EP67 spanning the N-terminus, and also the ECL2, drive the weaker potency and efficacy of EP67 on C5aR1. As expected, the distal stretch of EP67 makes similar contacts in the orthosteric binding pocket as C3a, including critical interactions of the terminal arginine with Arg^5.42^, Tyr^6.51^, Asp^7.35^ through hydrogen bond, cation-π interaction, and ionic interaction, respectively (**Figure S18A and S21B-C**). Finally, there is a notable difference between EP67-bound C3aR vs. C5aR1 at the level of positioning of the cytoplasmic end of TM7 and Helix8, wherein the TM7 is relatively more outward shifted in C5aR1 and Helix8 is oriented at an angle of 70° compared to C3aR (**Figure 5M**). In contrast, superimposition of C3a-C3aR and C5a-C5aR1 does not reflect an analogous structural difference between the two receptors (**Figure 5M**). These observations underscore EP67-specific conformational fine-tuning of C3aR vs. C5aR1, which is likely to drive the notable differences in its potency and efficacy for the two receptors.

### Species-specific diversity in βarr binding and activation

Inspired by the striking species-specific differences in the pharmacology at these receptors observed here, we compared the primary sequences of the human and mouse receptors with a focus on the putative phosphorylation signatures in their carboxyl-terminus that typically drive βarr interaction. Recent studies have identified a P-X-P-P type phosphorylation motif in GPCRs, where P represents a pSer/pThr, as a broadly conserved mechanism for engaging βarrs leading to their inter-domain rotation and activation^80,81^. Interestingly, we observed that both human and mouse C3aR contain a single P-X-P-P motif while the mouse C5aR1 contains two plausible P-X-P-P motifs unlike human C5aR1 which contains one (**Figure 6A)**. This presented a unique opportunity to test the hypothesis if this difference between the human and mouse receptor imparts an impact on βarr binding and activation. Accordingly, we determined the cryo-EM structures of βarr1 and 2 in complex with a phosphopeptide derived from the carboxyl-terminus of mouse C5aR1 (mC5aR1pp) harboring two juxtaposed P-X-P-P motifs (**Figure 6A)** at an overall resolution of 2.7-2.8Å (**Figure 6B and S6, and S10A-B**). The unambiguous densities allowed the placement of all the components including mC5aR1pp in the map with confidence, and a structural comparison with the basal states confirmed mC5aR1pp-induced activation of βarr1 and 2 as reflected by inter-domain rotation and disruption of the three-element and polar-core network, the hallmarks of βarr activation (**Figure 6C-E and S16C-D**).

**Fig. 6.**
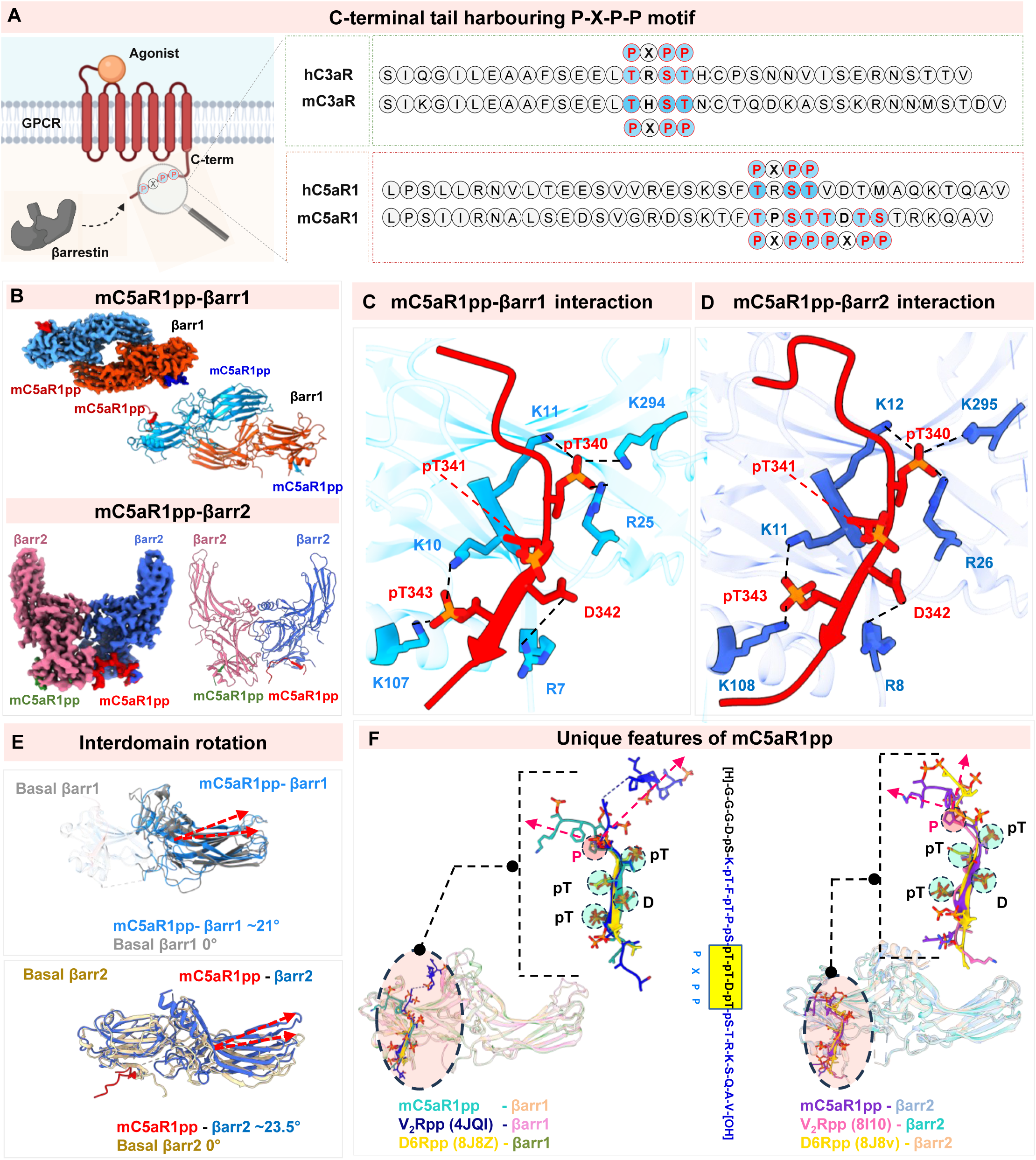
Structural insights into C5aR1 C-tail and βarr1/2. (A) Schematic representation of the C-terminal tail residues of h and mC3aR and C5aR1, highlighting the “P-X-P-P” motifs. MC5aR1 contains two successive P-X-P-P motifs. (B) Cryo-EM density maps and cartoon representations corresponding to the dimeric assemblies of βarr1 (top) and βarr2 (bottom) bound to mC5aR1pp are presented. (C-D) A key charge–charge interaction between mC5aR1pp and a Lys/Arg residue in the N-domain stabilises the positioning of mC5aR1pp within the N-domain groove of βarr1/2. (E) Interdomain rotation of βarr1 (top) and βarr2 (bottom) bound to mC5aR1pp are compared to the basal state structures of βarr1 (PDB: 1G4M) and βarr2 (PDB: 8J9K), respectively. (F) Structural superimposition of βarr1 and βarr2 bound to mC5aR1pp, V2Rpp, and D6Rpp reveals a unique feature of the mC5aR1 receptor: the presence of a proline at the ‘X’ position in the first P-X-P-P motif causes βarrs to slide down and preferentially engage with the second P-X-P-P motif.

Structural superimposition of mC5aR1pp-bound βarr1/2 with that of hC5aR1pp-bound βarr1/2 reported earlier reveal some intriguing and unanticipated differences despite an overall similar mode of binding and activation. For example, it is a variation of P-X-P-P motif in mC5aR1pp (i.e. Thr^340^-Thr^341^-Asp^342^-Thr^343^) that positions itself in the N-domain binding groove of βarr1/2, in an analogous position observed for the P-X-P-P motif in the hC5aR1pp-βarr1/2 structures^81^ (**Figure 6F and 7A)**. Still, this variant P-X-P-P engages an identical set of Lys/Arg residues in βarrs (K-R-K signature) that are recently identified to represent a lock-and-key type mechanism, and the Asp^342^ engages the same residue (i.e. Arg^7^/Arg^8^) in βarrs that is otherwise engaged by a phosphorylated Ser/Thr in a typical P-X-P-P motif (**Figure 7A-B)^81^**. This observation broadens the previously identified repertoire of P-X-P-P signature in GPCRs that plays a decisive role in βarr interaction and activation. Interestingly, a closer analysis of the mC5aR1pp positioning on βarrs reveals that the proline residue present in the first P-X-P-P motif as “X” results in a bending of the phosphorylated peptide, thereby rendering it out-of-place to slide into the binding groove on βarrs, and instead, propels the P-X-P-P variant, which is a part of the extended second P-X-P-P motif, to occupy the binding interface.

**Fig. 7:**
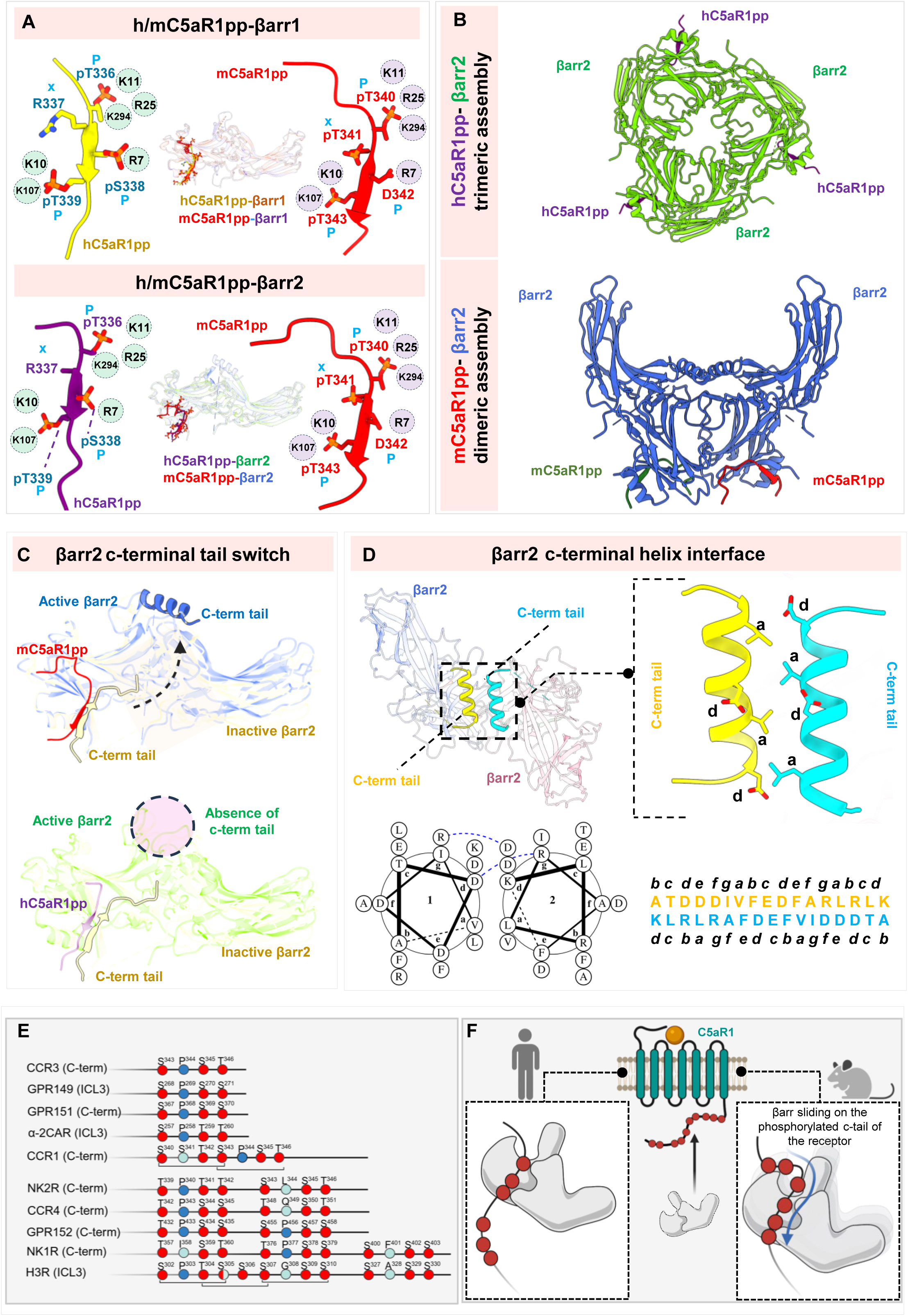
Species-specific diversity in βarr binding and activation. (A) Structural superimposition of βarr1 and βarr2 bound to h and mC5aR1pp (top left and bottom left) and the arrestin residues interacting with the phosphopeptides are shown (top right and bottom right). (B) Cartoon representation of the βarr2 trimeric assembly bound to hC5aR1pp (top) and dimeric assembly bound to mC5aR1pp (bottom). (C) Conformation transition of the βarr2 C-terminal residues in the basal state (PDB: 8J9K) to an α-helical conformation in the mC5aR1pp bound active state (top). This conformational switch is absent in the hC5aR1pp-bound βarr2 (PDB: 8I0Z) (bottom). (D) Dimeric organization of the mC5aR1pp-bound βarr2 structure (top left) and the antiparallel coiled-coil formed by the C-terminal helix of βarr2 at the dimeric interface (top right) are presented. Helical wheel (top left) and heptad representation (bottom left) of the antiparallel coiled-coil formed by the βarr2 C-terminal residues are shown. (E) List of GPCRs harbouring the P-X-P-P sequence at their C-tail or ICL3, where at least one of the P-X-P-P contains a proline residue at the X position. Red colour indicates pS/pT, blue colour indicates the proline residue, and light blue colour indicates any residue other than proline at the “X” position. (F) Diagrammatic representation of the sliding mechanism of βarr on phosphorylated receptor c-tail.

Another key difference observed between mC5aR1pp-βarr2 vs. hC5aR1pp-βarr2 is at the level of their oligomeric state in the corresponding structures. The mC5aR1pp-bound βarr2 exhibits a dimeric assembly in the cryo-EM structure while the hC5aR1pp-βarr2 revealed a trimeric assembly^81^ (**Figure 7C-D)**. The two protomers of βarr2 in mC5aR1pp-bound structure assemble primarily via the finger loop residues harboring an overall large, buried surface area of ∼5,500Å^2^, and they are related to each other through a two-fold symmetry (**Figure 7C-D**). Interestingly, a part of the carboxyl-terminus of βarr2 spanning the residues from 392 to 408, which forms a β-strand in the basal state and contributes to the auto-inhibition of βarr2, adopts an α-helical conformation and positions itself in the central crest of βarr2 (**Figure 7E)**. Both the protomers in the dimeric assembly show the same pattern, and these α-helices of the two protomers align themselves with each other at the dimeric interface to form an anti-parallel coiled-coil conformation (**Figure 7F**). This β-strand to α-helix transition and the specific arrangement is reminiscent of that observed previously for βarr2 in complex with the carboxyl-terminus of an intrinsically-biased, βarr-coupled 7TMR (D6R)^82^.

## Discussion

The complement anaphylatoxin receptors continue to be sought-after therapeutic targets due to their critical role in mediating the physiological responses of the terminal peptides generated following complement cascade activation^3,5^. Moreover, from a receptor pharmacology point of view, they exemplify an intriguing system encoding diverse ligand recognition and signaling-bias at the level for ligands and the receptors^6^. The spectrum of structure and associated pharmacology data presented here uncovers several interesting mysteries encoded by these receptors. In particular, the identification of a “Five-Point-Switch” in the orthosteric binding pocket of C3aR and C5aR1 not only provides a convergent mechanism of diverse ligand recognition but also raises a tantalizing possibility that additional naturally-occurring peptides, or, those provided by the human microbiota, may also serve as endogenous modulators of these receptors. Although such a scenario remains to be systematically explored and experimentally validated, the emerging paradigm of intracellular complement activation with the conceptual framework of the complosome may already provide preliminary support for such a hypothesis^83^.

The striking pharmacology and selectivity observed here for some of the agonists may have significant implications and impact on novel drug discovery focused on these receptors in future studies. For example, TLQP21 has emerged as an interesting lead molecule to engage C3aR from the therapeutic perspective of obesity considering its ability to mediate lipolysis and resting state energy expenditure in mouse models^30, 37^. However, due to its limited ability to activate the human C3aR, native hTLQP21 is unlikely to be a promising therapeutic candidate. However, our structural data now provide an opportunity to further explore the sequence space encoded by TLQP21 to gain reactivity towards hC3aR, and thereby, develop a new lead molecule with therapeutic potential. Similarly, there have been confounding reports in the literature regarding EP67-mediated beneficial outcomes in pathogenic infection models where the specific contribution of C3aR vs. C5aR1 has been unclear^48, 49^. Our study now reveals a clear preference of EP67 for the human C3aR compared to human C5aR1 in terms of transducer-coupling and also provides a structural rationale for this effect. Given recent preclinical studies demonstrating promising outcomes for EP67 therapy in community-acquired methicillin-resistant *S. aureus* (CA-MRSA) mouse models, our study now provides a template to further optimize the efficacy and selectivity of this ligand with potential for an improved therapeutic window^49^. Our study also unequivocally establishes the agonistic nature of SB290157 and JR14a, which were originally and still widely reported as antagonists, and demonstrates a structural explanation for their ability to activate the receptor.

A key finding in our study is the discovery that C5a^-d-Arg^ acts as a G-protein-biased agonist at C5aR1 unlike C3a^-d-Arg^, which nearly completely loses the ability to activate C3aR. This is particularly important to redefine the paradigm of a naturally-encoded mechanism to dampen the inflammatory response drive by C5a to avoid bystander inflammation. For example, a likely scenario is that the C5a^-d-Arg^-induced G-protein response continues to recruit neutrophils and macrophages to help clear pathogenic infections, while the reduction of inflammatory cytokines avoids excessive inflammation. At the molecular level, our study also provides further support that the N-terminus of C5aR1 but not C3aR is critical for anaphylatoxin interaction, and this helps rationalize pharmacology data regarding the relatively better potency and efficacy of peptide agonists at C3aR. Along the same line, it also underscores the challenges associated with small molecule agonists of C5aR1 while the same for C3aR has been rather more straightforward to design^69^.

Finally, the observation on distinct oligomeric state of βarr2 for the human vs. mouse C5aR1, and the engagement via an alternative P-X-P-P motif, further underscores previously unappreciated fine-tuning in the receptor-transducer system. In particular, the preferential engagement of βarrs to a specific phosphorylation motif depending on the adjacent sequence context, even when more than one options may be available, could be an added mechanism to encode conformational and functional diversity. Still, the conserved nature of engagement of the alternate P-X-P-P motif in mC5aR1pp with the K-R-K lock in βarrs reiterates that a certain level of intermolecular interactions must be satisfied for stable βarr binding and activation. As mentioned earlier, the proline residue present at position “X” in the first P-X-P-P of mC5aR1pp renders it ineffective to engage with the K-R-K lock due to a bending of the phosphopeptide, and hence, the adjacent P-X-P-P slides in place. Interestingly, several other human GPCRs also harbor a similar scenario with either a single or multiple P-X-P-P motif with a proline residue at the “X” position (**Figure 7G**), and therefore, it is possible that they also recapitulate a similar mechanism to engage βarrs as observed here for the mC5aR1pp. Based on these observations, we propose a “Sliding Mechanism” to guide βarrs on GPCRs wherein βarrs scan the phosphorylated carboxyl-terminus (or ICL3), and they stall once a P-X-P-P signature (or, a variant thereof) is encountered to effectively engage the K-R-K lock, leading to βarr activation (**Figure 7H**). However, further studies are essential to probe this hypothesis at molecular level and decipher the intricate details.

In summary, we elucidate multiple layers of previously unappreciated functional specialization encoded in the complement anaphylatoxin receptor system using an integrative experimental approach, and we anticipate that these molecular insights will help overcome the existing hurdles to guide novel therapeutic design targeting these receptors.

## Acknowledgements

Research on complement anaphylatoxin receptors in A.K.S.’s laboratory is currently supported by the Senior Fellowship of the DBT Wellcome Trust India Alliance (IA/S/20/1/504916), the Indian Council of Medical research (EMDR/SG/14/2024-01-02127), The Department of Science and Technology (DST/TTI/TC/AMR/COE/2023/5). Part of the cryo-EM data was collected at the National cryo-EM Facility at IIT Kanpur established with the support from ANRF/SERB (IPA/2020/000405). A.K.S. is the Sonu Agrawal Memorial Chair Professor. This research was also supported by the US National Institutes of Health grant R01AI82224 to C.G., and the National Health and Medical Research Council grant 2009957 to T.M.W. R.J.C.’s research is supported by the National Health and Medical Research Council (APP2012661). This research was also supported by the JSPS KAKENHI grant numbers 21H05037 (O.N.) and 23KJ0491 (F.K.S.), the Platform Project for Supporting Drug Discovery and Life Science Research (Basis for Supporting Innovative Drug Discovery and Life Science Research [BINDS]) from the Japan Agency for Medical Research and Development (AMED) grant numbers JP22ama121012 and JP22ama121002 (O.N.). We also acknowledge Australian Red Cross Lifeblood and human donors for providing blood for our research. We also thank Shachie Sinha and Ameesha Nigam for help with the functional assays on C3aR and C5aR1 mutants; and Ashna Reyaz, Debdatta Mukherjee, and Mallika Rani Prusty for help with protein purification.

## Authors’ contributions

SM and AD carried out the functional assays in HEK-293T cells with help from DT, SS, and PS; MKY prepared the complexes for cryo-EM with help from DT, NB and NR; RY collected and processed the cryo-EM data on mTLQP-mC3aR-Go, hTLQP-mC3aR-Go, mC5a-mC5aR1-Go, mC5a-hC5aR1-Go, mC5a^-d-Arg^-mC5aR1-Go, hC5a^-d-Arg^-mC5aR1-Go, EP54-mC3aR-Go, EP67-mC3aR-Go, EP67-hC3aR-Go, mC3a-mC3aR-Go, mC5aR1pp-βarr2-Fab30, mC5aR1pp-βarr1-Fab30 complexes with help from HAK and HZ; KS collected the data and solved the structure for SB290197-C3aR-G-protein complex with FKS; JNF, JM and JDL carried out the microglia experiments with additional experimental support from CSC; RC synthesised TLQP21 peptides with experimental validation by XXL; MKG and RB processed the cryo-EM data on C5a^pep^-hC5aR1-Go, EP54-hC5aR1-Go, JR14a-hC3aR-Go, EP67-mC5aR1-Go, EP67-hC5aR1-Go, JR14a-mC3aR-Go complexes, refined the structures, carried out the analyses, and prepared the figures with DT, NR, SM, AD, MKY and GR; TMW, ON, CG and AKS supervised the overall project. All authors contributed to writing and editing the manuscript.

## MATERIALS AND METHODS

### General reagents and chemicals

Most of the general reagents were purchased from Sigma-Aldrich unless otherwise specified. Dulbecco’s Modified Eagle’s Medium (DMEM), Trypsin-EDTA, Fetal Bovine Serum (FBS), Phosphate-Buffered Saline (PBS), Hank’s Balanced Salt Solution (HBSS), and Penicillin-Streptomycin solution were obtained from Thermo Fisher Scientific. HEK-293T cells (ATCC) were maintained in DMEM (Gibco, Cat. No: 12800-017) supplemented with 10% (v/v) FBS (Gibco, Cat. No: 10270-106), 100 U/mL penicillin, and 100 μg/mL streptomycin (Gibco, Cat. No: 15140122) at 37 °C under 5% CO₂. *Sf*9 cells were cultured in protein-free media (Gibco, Cat. No: 10902-088) at 27 °C at 125 rpm. Constructs for C3aR and C5aR1 expression in HEK-293T and *Sf9* were designed as described previously^77^. The constructs used in the NanoBiT-based βarr1/2 recruitment assay, endosomal localization assays and the bystander mode of the βarr1 recruitment assay were designed as described previously^77, 84, 85^. The G-protein subunit constructs used in the dissociation assays were generously provided by Asuka Inoue. Mutants of hC3aR, mC3aR, hC5aR1, and mC5aR1 were generated using the Q5 Site-Directed Mutagenesis Kit (NEB, Cat. No: E0554S). All DNA constructs were verified by sequencing at Macrogen. Peptides TLQP21, EP54, C5a^pep^ and EP67 were synthesized by GenScript or synthesised as previously described^64^. Small-molecule compounds SB290157 and JR14a were obtained from MedChem Express (Cat. No: HY-101502A for SB290157; Cat. No: HY-138161 for JR14a). M2-HRP coupled anti-FLAG antibodies were purchased from Sigma-Aldrich.

### GloSensor assay to measure cAMP response

Change in intracellular cAMP levels were measured using the GloSensor assay, as previously described^61^, to assess the effect of ligand-induced Gi-mediated signaling. Briefly, HEK-293T cells were transfected with 3.5 μg of either hC3aR, mC3aR, hC5aR1, or mC5aR1 along with 3.5 μg of the F22 plasmid (Promega, Cat. No: E2301). After 14–16 h of transfection, cells were trypsinized and resuspended in assay buffer (20 mM HEPES, pH 7.4, 1 X HBSS, and 0.5 mg/mL D-luciferin (GoldBio, Cat. No: LUCNA-1G). Later, 100 µl of transfected cells were seeded into 96-well plates at a density of 0.2 million cells per well. The cells were incubated at 37 °C for 1.5 h, followed by an additional 30 min incubation at room temperature. Basal luminescence was measured for three cycles. After the basal readings, 5 μM forskolin was added to each well, and luminescence was recorded for eight cycles until the readings stabilized. Subsequently, ligand was added at the indicated concentration, and luminescence was recorded for 15 cycles. The signal obtained was normalized to forskolin values, with the lowest ligand concentration set to 100%. Normalized data was subjected to Nonlinear regression analysis and dose-dependent response was plotted using GraphPad Prism 10 software.

### NanoBiT-based G-protein dissociation assay

Ligand-induced G-protein activation was measured using a previously described NanoBiT-based G-protein dissociation assay^86^. G-protein subunits consisting of the LgBiT-tagged Gα subunit and SmBiT-tagged Gγ2 (C68S mutation) subunit, along with the untagged Gβ1 subunit (pcDNA3.1), were co-expressed with the N-terminal FLAG-tagged C3aR or C5aR1 receptors. The ligand-induced change in luminescence signal was then measured. Briefly, HEK-293T cells were transfected with a plasmid mixture containing 1 µg LgBiT-GαoA, 4 µg Gβ1, 4 µg SmBiT-Gγ2 (C68S), and 1 µg of receptor plasmid (hC3aR, mC3aR, hC5aR1, or mC5aR1). After 14–16 h of transfection, cells were harvested by using Trypsin-EDTA and DMEM supplemented with 10% FBS and resuspended in NanoBiT buffer (5 mM HEPES, pH 7.4, 1 X HBSS, 0.01% BSA, and 10 µM coelenterazine (Gold Bio, Cat. No: CZ2.5). 100 μL of resuspended cells were then seeded into a 96-well plate at a density of 0.1 million cells per well. Plate was then incubated at 37 °C for 1.5 h, followed by an additional 30 min incubation at room temperature. Basal luminescence was recorded for three cycles using a Fluostar Omega plate reader. Following basal luminescence measurements, cells were stimulated with dose-dependent concentrations of ligand, leading to a reduction in luminescence signal. Ligand-induced response at 10 min was taken for analysis, wherein basal corrected response was percentage normalized relative to the lowest ligand concentration. The resulting data were plotted using GraphPad Prism 10 software.

### NanoBiT-based β-arrestin recruitment assay

Ligand-induced recruitment of β-arrestin (βarr1/2) was performed as described previously^81, 87^. Briefly, HEK-293T cells were transfected with 3.5 µg of C-terminal SmBiT-tagged hC3aR, mC3aR, hC5aR1, or mC5aR1, along with N-terminal LgBiT-tagged βarr1/2. For hC3aR and mC3aR mutant constructs, the bystander mode of βarr1 recruitment was employed. In this case, HEK-293T cells were transiently transfected with 3 µg of receptor, 5 µg of an N-terminal LgBiT-fused CAAX motif, and 2 µg of N-terminal SmBiT-tagged βarr1. After 16–18 h of transfection, the cells were trypsinized and harvested. The cell pellets were resuspended in assay buffer containing 5 mM HEPES, pH 7.4, 1 X HBSS, 0.01% BSA, and 10 µM coelenterazine. The cells were seeded into a 96-well plate at a density of 0.1 million cells per well in a 100 µL volume. Seeded plates were incubated at 37 °C for 1.5 h, followed by an additional 30 min incubation at room temperature. After incubation, basal luminescence was recorded for three cycles using the FLUOstar Omega plate reader. Cells were then stimulated with the indicated ligand concentrations, and ligand-induced increases in luminescence were recorded for 15 cycles. The responses from cycles 5–10 was averaged, and fold normalization was performed, considering the lowest ligand concentration as 1. The fold-normalized values were then plotted using GraphPad Prism 10 software.

### NanoBiT-based endosomal trafficking assay

Ligand-induced endosomal localization of the receptor along with βarr1/2 was assessed using a NanoBiT-based complementation assay, similar to the βarr1/2 recruitment assay described above. Briefly, HEK-293T cells were transfected with 3 µg of untagged, hC5aR1, or mC5aR1, along with 5 µg of an N-terminal LgBiT-tagged FYVE domain and N-terminal SmBiT-tagged βarr1/2. The subsequent steps followed the same procedure as the βarr1/2 recruitment assay.

### Receptor surface expression assay

To assess cell surface expression of the corresponding receptors, whole cell surface ELISA was performed, as previously described^88^. Briefly, transfected cells were seeded at a density of 0.2 million cells per well 24 h post-transfection in 24-well plates (pre-coated with 0.01% poly-D-Lysine) and incubated for 24 h at 37 °C in a CO_2_ incubator. Once cells were well adhered, the media was removed by aspiration and the cells were washed once with 1 X TBS. This was followed by treatment with 300 μL of 4% PFA for 20 min to fix the cells on ice. Excess PFA was removed by washing the cells thrice with 400 μL of 1 X TBS. The wells were blocked by incubating with 200 μL 1% BSA for 1.5 h. 200 μL of anti-FLAG M2-HRP antibody (Sigma, Cat. no: A8592) at a dilution of 1:10,000 was added to the wells and incubated for an additional 1.5 h. To develop the signal, 200 μL of TMB-substrate (Thermo Scientific, Cat. no: 34028) was added to the wells and incubated till the development of adequate color. Reaction was quenched by transferring 100 μL of the solution to a 96-well plate containing 100 μL of 1 M H_2_SO_4_ and absorbance was measured at 450 nm using PerkinElmer VICTOR^TM^ X4 plate reader. To estimate the number of cells fixed in each well, excess TMB was first removed by washing once with 400 μL of 1X TBS, and the cells were incubated for 20 min in 0.2% Janus green B stain (Sigma, Cat. no. 201677). After removing excess stain by repeated washing with Milli Q water, a signal was developed by adding 800 μL of 0.1 N HCl to each well and read at 595 nm. Signal was normalized by dividing the reading obtained at 450 nm with the reading obtained at 595 nm. Receptor surface expression was fold normalized to empty pcDNA vector-transfected cells and plotted in the GraphPad Prism 10 software.

### Primary cell assays

#### Cell culture

Primary mouse microglia cultures were isolated from freshly dissected brains of C57 pups at P1-3. Mixed glial cultures were cultured in Dulbecco’s Modified Eagle Medium/Nutrient Mixture 12 (DMEM/F-12; ThermoFisher Scientific, Waltham, MA) with the following supplements: 10% fetal calf serum (FCS), 1% Glutamine (Gibco, Grand Island, NY), 1% Penicillin/Streptomycin (Gibco), 1% Sodium Pyruvate (Gibco), and 1% MEM non-essential Amino Acids (Sigma-Aldrich, St Louis, MO). Glial cultures were seeded onto poly-d-lysine-coated flasks and incubated for 14 days at 37 °C and 5% CO₂. Primary microglia were positively selected from the mixed glial cultures using the EasySep™ Mouse CD11b Positive Selection Kit II (StemCell Technologies, Vancouver, BC) according to the manufacturer’s instructions and resuspended in DMEM, high glucose (ThermoFisher Scientific) supplemented with 10% FCS and 1% Penicillin/Streptomycin. Cells were seeded in 96-well plates at a density of 5,000 cells per well and treated with hTLQP21 and mTLQP21 (1 µM, 10 µM, 100 µM), and LPS (100 ng/mL) for 24 h prior to the collection of cell supernatant.

Peripheral blood mononuclear cells (PBMCs) were taken from healthy participants and differentiated into human monocyte-derived microglial cells (MDMi)^89^. Briefly, peripheral venous blood samples were collected in ethylenediaminetetraacetic acid (EDTA) tubes (Beckton-Dickson, NJ). PBMCs were isolated from blood samples using the SepMate™ tubes (StemCell Technologies) according to the manufacturer’s instructions and seeded onto Matrigel-coated 96-well plates at 200,000 cells/well. Cells were incubated overnight at 37 °C and 5% CO_2_ prior to culturing in serum-free RPMI-1640 GlutaMax medium (ThermoFisher Scientific) supplemented with recombinant human interleukin-34 (IL-34, 100 ng/mL; PeproTech, Cranbury, NJ), recombinant human granulocyte-macrophage colony-stimulating factor (GM-CSF, 10 ng/mL; PeproTech), and 1% Penicillin/Streptomycin (Gibco) for 14 days. MDMi were treated with hTLQP21 and mTLQP21 (1 µM, 10 µM, 100 µM), and LPS (100 ng/mL) for 24 h prior to the collection of cell supernatant.

### Cytokine and chemokine measurement for mouse primary microglia

Cytokine and chemokine levels in the mouse primary microglia cell supernatants were quantified using the BioLegend LEGENDplex Mouse 13-plex Inflammation Panel Kit (BioLegend, San Diego, CA). Similarly, cytokine and chemokine levels in MDMi cell supernatants were quantified using the BioLegend LEGENDplex™ Human 13-plex Inflammation panel kit (BioLegend). Protocols were followed according to the manufacturer’s instructions.

### Data collection, processing and analysis

All experiments were conducted in triplicate and repeated on three separate occasions or using cells from at least 3 independent human donors and (for MDMi) mice (primary microglia), unless otherwise specified. Data were analysed using GraphPad Prims software (Prism 10.4). Data from each individual repeat was normalised accordingly (as specified in the figure legends) before being combined and expressed as mean ± standard error of the mean (S.E.M.) unless otherwise described. Concentration-response curves were plotted using combined data and analysed to determine the respective potency values.

### Expression and purification of C3a, C5a, and C5a^-d-Arg^

Gene encoding C3a, C5a and C3a^-d-Arg^/C5a^-d-Arg^ (Human and mouse) were cloned in pET-32a(+) vector with a Trx-6X-His tag at the N-terminal end and purified following the previously described protocol with some modifications^90^. Briefly, freshly transformed *E. coli* Shuffle cells were inoculated in 50 mL of LB media containing 100 µg/mL ampicillin for the starter culture at 30 °C. The overnight-grown primary culture was then inoculated into 1.5 L of LB media with 100 µg/mL ampicillin, and the culture was allowed to grow at 30 °C. When the optical density (O.D.) reached approximately 0.8-1.0, the culture was induced with 1 mM IPTG and then shifted to 16 °C for overnight induction. The cells were harvested and incubated with 1 mg/mL lysozyme in a buffer containing 50 mM HEPES, pH 7.4, 300 mM NaCl, 20 mM imidazole, 1 mM PMSF, and 2 mM benzamidine for 2 h at 4 °C. Following this, the cells were disrupted using ultrasonication, and the cell debris was removed by high-speed centrifugation. C3a/C5a/C3a^-d-Arg^/C5a^-d-Arg^ were enriched on Ni-NTA resins using gravity flow. Non-specific proteins were removed through extensive washing with a buffer containing 50 mM HEPES, pH 7.4, 1 M NaCl, and 20 mM imidazole. The fusion protein(s) was then eluted using a buffer composed of 50 mM HEPES, pH 7.4, 150 mM NaCl, and 300 mM imidazole. The Trx-His tag was cleaved by treating the protein with TEV protease (1:20 w/w, TEV: protein) for 8-10 h at room temperature. The purified proteins were further cleaned using cation-exchange chromatography and stored at -80 °C with a final glycerol concentration of 10%.

### Expression and purification of C3aR and C5aR1

The human and mouse, C3aR and C5aR1 genes were codon-optimized and cloned into the pVL1393 vector with an N-terminal HA signal sequence, a FLAG tag, and a 21-amino-acid stretch of the M4 receptor N-terminal region (amino acids 2–23, ANFTPVNGSSGNQSVRLVTSSS) for purification. The receptor was expressed and purified from *Spodoptera frugiperda* (*Sf9*) cells using a baculovirus-mediated expression system. Briefly, the insect cells were harvested 72 h post-infection and lysed through sequential douncing in the following buffers: a low-salt buffer (20 mM HEPES, pH 7.4, 10 mM MgCl_2_, 20 mM KCl, 1 mM PMSF, and 2 mM benzamidine), a high-salt buffer (20 mM HEPES, pH 7.4, 1 M NaCl, 10 mM MgCl_2_, 20 mM KCl, 1 mM PMSF, and 2 mM benzamidine), and a solubilisation buffer (20 mM HEPES, pH 7.4, 450 mM NaCl, 2 mM CaCl_2_, 1 mM PMSF, 2 mM benzamidine, and 2 mM iodoacetamide). After lysis, the receptor was solubilized for 2 h at 4 °C with continuous stirring in a solution containing 0.5% L-MNG (Anatrace, Cat. no: NG310) and 0.01% cholesteryl hemisuccinate (Sigma, Cat. no: C6512). Following solubilization, the salt concentration was reduced to 150 mM, and the receptor was purified using an M1-FLAG column. To remove non-specific proteins from the FLAG beads, three washes with a low-salt buffer (20 mM HEPES, pH 7.4, 2 mM CaCl_2_, 0.01% CHS, 0.01% L-MNG) were alternated with two washes using a high-salt buffer (20 mM HEPES, pH 7.4, 450 mM NaCl, 2 mM CaCl_2_, 0.01% L-MNG). The bound receptor was eluted using a FLAG elution buffer containing 20 mM HEPES, pH 7.4, 150 mM NaCl, 0.01% LMNG, 2 mM EDTA, and 250 µg/mL FLAG peptide. The purified receptor was incubated on ice with 2 mM iodoacetamide for 30 min. Subsequently, 2 mM L-cysteine was added, and the mixture was incubated for 30 min. The purified receptor was concentrated using a 30 kDa MWCO concentrator and stored at -80 °C in 10% glycerol till further use. 100 nM of h/mC3a, mC5a, h/mC5a^-d-Arg^ or 1 µM of hTLQP21, mTLQP21, EP67, EP54, SB290157, JR14a, or C5a^pep^ were kept in all steps of receptor purification.

### Expression and purification of G-proteins

The miniGαo construct was engineered by incorporating an ScFv16 binding sequence at the N-terminus of the previously described miniGαo^91, 92^. The miniGαo1 gene was cloned into a pET-15b(+) vector with an N-terminal 6X-His tag for expression. The engineered construct was transformed into *E. coli* BL21(DE3) cells for protein expression. A starter culture was grown in LB media and incubated at 37 °C with shaking at 160 rpm for 6-8 h. This starter culture was used to inoculate an overnight culture in LB media supplemented with 0.2% glucose and grown at 30 °C. 15 mL of the overnight culture was transferred to 1.5 L of Terrific Broth (TB) media. Protein expression was induced with 50 μM IPTG when the culture reached an optical density (O.D_600_) of 0.8, followed by incubation at 25 °C for 18-20 h. Cells were harvested and lysed in buffer containing 40 mM HEPES, pH 7.4, 100 mM NaCl, 10 mM imidazole, 10% glycerol, 5 mM MgCl₂, 1 mM PMSF, and 2 mM benzamidine, supplemented with 1 mg/mL lysozyme, 50 μM GDP, and 100 μM DTT. Cell debris was removed by centrifugation at 18,000 rpm for 30 min at 4 °C. The protein was purified using Ni-NTA affinity chromatography. The bound protein was washed extensively with buffer containing 20 mM HEPES, pH 7.4, 500 mM NaCl, 40 mM imidazole, 10% glycerol, 50 μM GDP, and 1 mM MgCl₂. The protein was then eluted using 20 mM HEPES, pH 7.4, 100 mM NaCl, 10% glycerol, and 500 mM imidazole. The purified protein fractions were pooled and stored at -80 °C in glycerol until further use. The genes encoding the Gβ1 and Gγ2 subunits were cloned into a bicistronic pVL1392-based vector with an N-terminal histidine tag on Gβ1. These constructs were expressed in *Sf9* cells using the baculovirus expression system. The cells were harvested 72 h post-infection and resuspended in a lysis buffer containing 20 mM Tris-Cl, pH 8.0, 150 mM NaCl, 10% glycerol, 1 mM PMSF, 2 mM benzamidine, and 1 mM MgCl_2_. The cells were disrupted using a dounce homogenizer and then centrifuged at 4 °C for 40 min at 18,000 rpm. The resulting pellet was resuspended and dounced again in a solubilization buffer (20 mM Tris-Cl, pH 8.0, 150 mM NaCl, 10% glycerol, 1% DDM, 5 mM β-ME (β-mercaptoethanol), 10 mM imidazole, 1 mM PMSF, and 2 mM benzamidine) and solubilized at 4 °C under constant stirring for 2 h. Cell debris was pelleted down by centrifuging at 20,000 rpm for 60 min at 4 °C. The protein was enriched on Ni-NTA resin. After extensive washing with a buffer containing 20 mM Tris-Cl, pH 8.0, 150 mM NaCl, 30 mM imidazole, and 0.02% DDM, the protein was eluted with 300 mM imidazole in a buffer composed of 20 mM Tris-Cl, pH 8.0, 150 mM NaCl, and 0.01% LMNG. Eluted protein was concentrated with a 10 kDa MWCO concentrator (Cytiva Cat no: GE28-9322-96) and stored at -80 °C with 10% glycerol.

### Expression and purification of ScFv16

The ScFv16 gene was cloned into a pET-42a(+) vector with an in-frame N-terminal 10X-His-MBP tag followed by a TEV protease cleavage site. The construct was transformed into *E. coli* Rosetta (DE3) cells for protein expression^93^. An overnight culture was inoculated in 1 L of 2XYT media supplemented with 0.5% glucose and 5 mM MgSO₄. When the culture reached an optical density (O.D_600_) ∼ 1.0, protein expression was induced with 250 µM IPTG, and cells were grown at 18 °C for 16-18 h. The harvested cells were resuspended in lysis buffer containing 20 mM HEPES, pH 7.4, 200 mM NaCl, 2 mM benzamidine, and 1 mM PMSF, followed by incubation at 4 °C for 1 h with continuous stirring. The cells were lysed by ultrasonication, and cellular debris was removed by centrifugation at 18,000 rpm for 40 min at 4 °C. Protein was enriched on Ni-NTA resins, and non-specifically bound proteins were removed by extensive washing with buffer containing 20 mM HEPES, pH 7.4, 200 mM NaCl, and 50 mM imidazole. The protein was then eluted using the same buffer containing 300 mM imidazole. Further purification was performed using amylose resin (NEB, Cat. no: E8021L), where the Ni-NTA eluate was applied and washed with buffer containing 20 mM HEPES, pH 7.4, and 200 mM NaCl to remove any remaining non-specific proteins. Protein was eluted with 10 mM maltose (prepared in 20 mM HEPES, pH 7.4, 200 mM NaCl), and the His-MBP tag was removed by overnight treatment with TEV protease. Following TEV protease cleavage, the tag-free ScFv16 was separated from the cleaved tag by passing the mixture through Ni-NTA resin. The purified protein was then concentrated and subjected to final polishing by size exclusion chromatography using a Hi Load Superdex 200 PG 16/600 column (Cytiva Life Sciences, Cat. no: 17517501). Purified protein was flash-frozen and stored at -80 °C with 10% glycerol.

### Reconstitution of the receptor G-protein complexes

The complexes were reconstituted following the previously published protocols^94–96^. Briefly, the purified receptors were incubated with 1.2 molar excess of Gαo, Gβ1γ2, and ScFv16 at room temperature for 2 h in the presence of 25 mU/mL apyrase (NEB, Cat. no: M0398S) and either 1 µM of mC3a, mC5a, hC5a^-d-Arg^, mC5a^-d-Arg^ or 10 µM hTLQP21, mTLQP21, EP67, EP54, SB290157, JR14a, or C5a^pep^ for complex formation. The G-protein complex was separated from unbound components by loading on Superose 6 increase 10/300 GL SEC column and analyzed by SDS-PAGE. Complex fractions were pooled and concentrated to ∼10-15 mg/ml using a 100 MWCO concentrator (Cytiva, Cat. no: GE28-9323-19) and stored at -80 °C until further use.

### Fab30 Purification

The Fab fragment production methodology followed a multi-step protocol^97^. *E. coli* M55244 cells (ATCC) transformed with a Fab plasmid, where an initial 50 mL seed culture was grown overnight at 30 °C, subsequently expanded to 1 L of 2XYT media inoculated with 5% seed culture and incubated for 8 h at 30 °C. Following this, cells were harvested and resuspended in 1 L of CRAP medium containing 100 μg/mL ampicillin, then further incubated for 24 h at 30°C. Cell lysis was performed using sonication with a specific buffer composition (50 mM HEPES, pH 8.0, 500 mM NaCl, 0.5% Triton X-100, 0.5 mM MgCl2), after which the lysate was heated to 65 °C for 30 min and immediately chilled on ice for 5 min. To obtain a clear supernatant, the lysate was centrifuged at 20,000 rpm for 30 min and loaded onto a Protein-L resin column at room temperature, with subsequent washing using a 50 mM HEPES, pH 8.0, 500 mM NaCl buffer. Protein elution was conducted using 100 mM acetic acid in tubes containing 1 M HEPES, pH 8.0, at 10% column volume to ensure rapid neutralization, followed by buffer exchange into 20 mM HEPES, pH 8.0, 100 mM NaCl using pre-packed de-salting columns (GE Healthcare Cat. no. 17085101), and final storage at -80 °C

### Expression and purification of βarrs

For the expression and purification of β-arrestins (βarrs), a previously established protocol was utilized^98^. In summary, the cDNAs of rat βarr1 and bovine βarr2^DM,81^ (referred to as βarr2 henceforth) were inserted into the pGEX-4T-3 vector, which includes a GST tag and a thrombin cleavage site. A single colony of *E. coli* BL21(DE3) was inoculated into 50 mL of TB medium containing 100 µg/mL ampicillin. After culturing until the optical density at 600 nm (OD600) reached 0.8-1.0, a secondary culture of 1.5 L Terrific Broth was inoculated and grown to the same OD_600_ range. The expression of βarrs was induced with 25 µM IPTG and incubated for an additional 16 h at 18 °C. The cultures were then harvested and stored at -80 °C for later use. Cell lysis was performed by sonicating the resuspended pellets in a lysis buffer containing 25 mM Tris, pH 8.5, 150 mM NaCl, 1 mM PMSF, 2 mM benzamidine, 1 mM EDTA, 5% glycerol, 2 mM dithiothreitol (DTT), and 1 mg/mL Lysozyme. The lysate was centrifuged at 18,000-20,000 rpm at 4 °C, followed by filtration through a 0.45 µm filter to yield a clear supernatant. Overnight batch binding was conducted with Glutathione Sepharose resin at 4 °C. The beads containing the bound proteins were thoroughly washed with a buffer comprising 25 mM Tris, pH 8.5, 150 mM NaCl, 2 mM DTT, and 0.02% n-dodecyl-β-D-maltopyranoside (DDM) after being transferred to Econo columns. Thrombin was added to the resin slurry at a concentration of 1 unit/µL in a ratio of dry resin to cleavage buffer of 1:1, where the cleavage buffer consisted of 25 mM Tris, pH 8.5, 350 mM NaCl, and 0.02% DDM, and incubated for 2 h at room temperature for on-column cleavage. The resulting pure, tag-free βarrs were eluted and further purified using Hi-Load Superdex 200 PG 16/600 column (Cytiva Life sciences, Cat. no: 17517501) with a running buffer of 25 mM Tris, pH 8.5, 350 mM NaCl, 2 mM DTT, and 0.02% DDM. The fractions containing βarrs were pooled and stored at -80 °C with the addition of 10% glycerol.

### Reconstitution of phosphopeptide-βarr-Fab complexes

A previously established protocol was utilized for reconstituting the phosphopeptide-βarr complex^81^. In summary, phosphopeptides were added in three-fold molar excess to βarrs and incubated for 45 min at room temperature to facilitate activation. After incubation, the Fab30 was mixed in a 1:1.5 ratio (βarr: Fab) and allowed to form complexes for 90 min at room temperature. The resulting complexes were purified using a Superose 6 Increase 10/300 GL gel-filtration column with a running buffer consisting of 20 mM HEPES, pH 7.4, 300 mM NaCl, 0.00075% LMNG, 0.00025% CHS, and 2 mM DTT, following concentration with 30,000 Da MWCO concentrators. Fractions containing the complex were pooled, concentrated to the desired level (8-10 mg/mL), and subsequently used for negative-staining electron microscopy (EM) and cryo-EM grid preparation.

### Negative stain electron microscopy

In order to judge homogeneity and purity of the samples prior to grid freezing for cryogenic electron microscopy, negative staining was performed on the complexes using uranyl formate (Polysciences) in accordance with previously published protocols^77, 99^. All the staining steps were performed at room temperature. In brief, 3.5 µL of the protein samples at a concentration of ∼0.02 mg/mL were dispensed onto freshly glow-discharged carbon/formvar coated Cu grids (PELCO, *Ted Pella Inc.*), allowed to adsorb for 1 min, and blotted off using a Whatman No.1 filter paper. The grid was then touched onto a drop of freshly prepared uranyl formate (Cat. No: 24762-1) stain (0.75%) and immediately blotted off using a filter paper to remove excess sample solution. This was followed by staining of the adhered samples by touching the grid onto a second drop of stain for 30 s and left on the bench for drying. Data collection of the samples was performed at 30,000 x magnification on a FEI Tecnai G2 12 Twin TEM (LaB_6_) operating at 120 kV and equipped with a Gatan CCD camera (4k x 4k). Data processing of the collected micrographs was performed in Relion 3.1.2^100–102^. Approximately 10,000 particles were automatically picked using the Gaussian blob picker, extracted and subjected to reference-free 2D classification. 2D class averages with clear secondary features were selected for preparing the figures.

### Cryo-EM sample preparation and data collection

The purified ligand-receptor-Go complexes (mTLQP-mC3aR-Go, hTLQP-mC3aR-Go, mC5a-mC5aR1-Go, mC5a-hC5aR1-Go, mC5a^-d-Arg^-mC5aR1-Go, hC5a^-d-Arg^-mC5aR1-Go, EP67-mC3aR-Go, EP67-hC3aR-Go, mC3a-mC3aR-Go, EP54-mC3aR-Go) were incubated with excess ligand at 4 °C for 1 h before grid freezing. Quantifoil R1.2/1.3 Au 200-mesh grids were glow discharged at 20 mA current for 60 s glow and 10 s hold using a PELCO easiGlow glow discharge system (Ted Pella). For vitrification, a 2.5 µL sample of the above-mentioned ligand-receptor-Go and phosphopeptide-βarrestin complexes (mC5aR1pp-βarr1-Fab30 and mC5aR1pp-βarr2-Fab30) were applied onto glow-discharged grids and blotted for 5-6 s with blot-force 0 at 4 °C and 100% humidity and plunged into liquid ethane using a Mark IV Vitrobot (Thermo Fisher Scientific).

Cryo-EM data were collected using an FEI Titan Krios microscope (Thermo Fisher Scientific) operating at 300 kV, equipped with a BioQuantum GIF energy filter (Gatan) with 20 eV slit width, with a K3 direct electron detector (Gatan) operating in super-resolution counting mode at 130,000 magnifications with a pixel size of 0.647 Å (2-fold hardware binning) using EPU data acquisition software (Thermo Fisher Scientific). Movies were recorded with 1.8 s exposure time, a defocus range of -0.8 to -3.4 µm, and a total dose of 40-60e^-^/Å^2^. The cryo-EM data for the mC5a-hC5aR1-Go and and EP54-mC3aR-Go samples were collected on a Glacios microscope (Thermo Fisher Scientific) operating at 200 kV with a Falcon 4 direct electron detector in counting mode. The movies were recorded at 150,000 magnification (0.92 Å pixel size) with an exposure time of 7.74 s, a defocus range of -0.8 to -3.0 µm, and a total dose of 58.6 e^-^/Å^2^ using EPU data acquisition software. The cryo-EM data for EP67-mC3aR-Go, EP67-hC3aR-Go, and mC3a-mC3aR-Go were collected on a Glacios microscope operating in Fringe-Free Imaging mode, enabling multiple shots per hole. The single-particle movies for EP67-mC3aR-Go and EP67-hC3aR-Go were collected at 190,000x magnification (0.73 Å pixel size) with an exposure time of 2.77 s, a defocus range of -0.8 to -3.2 µm, and a total dose of 50 e^-^/Å^2^. The data for mC3a-mC3aR-Go was collected at 150,000 magnification (0.92 Å pixel size) with an exposure time of 4.66 s, a defocus range of -0.8 to -3.2 µm, and a total dose of 50 e^-^/Å^2^.

Grids for JR14a-hC3aR-Go, JR14a-mC3aR-Go, EP67-mC5aR1-Go, EP67-hC5aR1-Go, C5a^pep^-hC5aR1-Go, and EP54-hC5aR1-Go were prepared by dispensing 3 μl of the complex samples (∼10-14 mg/mL) onto glow-discharged holey grids (Quantifoil Au or Cu 1.2/1.3 300 mesh). The excess sample was blotted off for 4 s with Whatman no. 1 filter paper and plunge-frozen in liquid ethane (-181 °C) using a FEI Vitrobot Mark IV at 100% relative humidity and 4 °C. The grids were clipped and stored in liquid nitrogen for data collection later. Data collection was carried out on a TFS Titan Krios microscope operating at 300 kV using a GIF Quantum energy filter and K3 detector (Gatan) in counting mode. Movies were recorded across a defocus range of -0.8 to -1.8 μm, at a nominal magnification of 105,000 x, pixel size of 0.86 Å/pix using EPU software. Movies were fractionated into 50 frames with a total dose of ∼55 e^-^/A^2^. For the SB219057-hC3aR-Go complex sample, 3 μL of the sample were dispensed onto glow discharged Quantifoil holey carbon grids (Au R1.2/1.3) using a PIB-10 glow-discharge (Vacuum Device Co., Ltd., https://www.shinkuu.co.jp/) and blotted using Ø55/20 mm, Grade 595 (Ted Pella, Inc.) filter paper. The grids were subsequently flash frozen in liquid ethane (-181 °C) using a Vitrobot Mark IV maintained at 100% humidity and 4 °C. Data collection was performed on a 300 kV Titan Krios microscope (G3i, Thermo Fisher Scientific) equipped with a K3 direct electron detector (Gatan) and BioQuantum K3 imaging filter. Movies were recorded in counting mode across a defocus range of -0.8 to -1.6 μm at a pixel size of 0.83 Å/px using EPU software (Thermo Fisher Scientific). Movies were fractionated into 48 frames with electron exposures of 47.4 e^-^/Å^2^.

### Data processing

Data processing of all the datasets was performed using cryoSPARC v4^103^ unless otherwise stated. Movies were imported and aligned using Patch motion correction (multi), followed by estimation of contrast transfer function using Patch CTF (multi). All the datasets of C3aR and C5aR1 and phosphopeptide-βarrestin complexes were processed using a similar strategy as outlined in Supplementary Figure 5-7. Briefly, particles were picked with the blob picker sub-program within the cryoSPARC suite followed by particle extraction with the appropriate box size. The extracted particle stacks were subjected to multiple rounds of 2D classification to discard junk particles. A clean particle stack against the 2D class averages with clear secondary features were used to generate ab-initio maps. Multiple rounds of heterogenous refinement were performed to clean the particle stacks further using the ab-initio models as reference. The 3D class with features of GPCR-G protein complex were subjected to non-uniform refinement followed by local refinement with a mask on the micelle. The resolution of all maps was calculated internally in cryoSPARC against the half-sets using the gold standard fourier shell correlation at 0.143.

For the mTLQP21-mC3aR-Go dataset, 8,573 micrographs with CTF resolution better than 4 Å were selected for further processing. The initial ab-initio model with features of a receptor-G protein complex exhibited anisotropy. To remove orientation bias and anisotropy, TOPAZ was used to train on a particle set with particles picked from denoised micrographs using the ResNet8 convolutional neural network model using a down-sampling factor of 8, and the expected number of particles per micrograph set to 500. The particles picked with TOPAZ were extracted and cleaned through iterative rounds of 2D classification, followed by multiclass ab-initio/hetero classification. Particles corresponding to the best class were subjected to non-uniform refinement followed by local refinement with a mask on the complex to eliminate noise around the reconstruction. Data collection, processing and model refinement statistics have been included as **Supplementary table S2.** Estimation of local resolution were performed with the Blocres sub-program, and map sharpening was performed with either an inbuilt program within the cryoSPARC v4 suite or deepEMhancer included in the Cosmic cryo-EM webserver^104^ to improve features for model building.

### Model building and refinement

For the mC3aR-Go complex, the starting model of mC3aR was generated with Swiss-model^105^ using the active hC3aR structure as the template. For the hC3aR, hC5aR1 and mC5aR1-Go complexes, the coordinates of PDB ID: 8I9L, 8IA2, and 8HPT, respectively, were used as the starting models for model building. For the phosphopeptide-βarrestin complexes, the initial coordinates of βarr1 and βarr2 were obtained from the cryo-EM structures of C5aR1pp-βarr-1 (PDB:8GO8) and D6Rpp-βarr2 (PDB:8GO9), respectively. Initial coordinates of Gαo, Gβ1 and Gγ2 were obtained from the cryo-EM structure of EP54-C3aR-Go (PDB: 8I95). These initial models were used to rigidly dock into the sharpened map using Chimera^106,107^, followed by flexible fitting of the coordinates using the “all-atom-refine” sub-module in COOT^108-110^. This was followed by iterative manual adjustments in COOT and *phenix.real_space_refine* within the Phenix suite^111,112^. The final models showed good statistics, with most of the residues lying in the most favored region of the Ramachandran plot (**Table S2**). All structural figures used in the manuscript have been prepared with Chimera and ChimeraX softwares^106,107^.

### Accession number

The cryo-EM maps and structures have been deposited in the EMDB and PDB with accession numbers PDB ID-9KV8, EMD-6258 (hTLQP21-mC3aR-Go); PDB ID-9KV6, EMD-62586 (mTLQP21-mC3aR-Go); PDB ID-9KWG, EMD-62610 (mC5a-mC5aR1-Go); PDB ID - 8JZP, EMD-36750 (mC5a-hC5aR1-Go); PDB ID-9KZK, EMD-62665 (mC3a-mC3aR-Go); PDB ID-9KWX, EMD-62619 (mC5a^-d-Arg^-mC5aR1-Go); PDB ID-9KX6, EMD-62624 (hC5a^-d-Arg^ mC5aR1-Go); PDB ID-9KZ2, EMD-62651 (EP67-hC3aR-Go); PDB ID-9KUG, EMD-62578 (EP67-hC5aR1-Go); PDB ID-9KZ8, EMD-62654 (EP67-mC3aR-Go); PDB ID-9KXS, EMD-62626 (EP67-mC5aR1-Go); PDB ID-9KUT, EMD-62580 (JR14a-hC3aR-Go); PDB ID-9L0H, EMD-62712 (JR14a-mC3aR-Go), PDB ID-9KVP, EMD-62598 (SB290157-hC3aR-Go-Full map), and PDB ID-9KVP, EMD-62598 (SB290157-hC3aR-Go ligand-receptor-focused map), PDB: 9UMR, EMD-64291 (C5a^pep^-hC5aR1-Go), PDB: 9UMJ, EMD-64276 (EP54-mC3aR-Go), PDB: 9UMX, EMD-64325 (EP54-hC5aR1-Go), PDB: 9KYU, EMD-62649 (mC5aR1pp-βarr1-Fab30), PDB: 9KY22, EMD-62636, (mC5aR1pp-βarr2-Fab30).

## References

1. Ricklin, D., Hajishengallis, G., Yang, K., and Lambris, J.D. (2010). Complement: a key system for immune surveillance and homeostasis. Nat Immunol 11, 785–797. 10.1038/ni.1923.

2. Mastellos, D.C., Hajishengallis, G., and Lambris, J.D. (2024). A guide to complement biology, pathology and therapeutic opportunity. Nat Rev Immunol 24, 118–141. 10.1038/s41577-023-00926-1.

3. Merle, N.S., Church, S.E., Fremeaux-Bacchi, V., and Roumenina, L.T. (2015). Complement System Part I - Molecular Mechanisms of Activation and Regulation. Front Immunol 6, 262. 10.3389/fimmu.2015.00262.

4. Esser, A.F. (1994). The membrane attack complex of complement. Assembly, structure and cytotoxic activity. Toxicology 87, 229–247. 10.1016/0300-483x(94)90253-4.

5. Bajic, G., Degn, S.E., Thiel, S., and Andersen, G.R. (2015). Complement activation, regulation, and molecular basis for complement-related diseases. EMBO J 34, 2735–2757. 10.15252/embj.201591881.

6. Klos, A., Wende, E., Wareham, K.J., and Monk, P.N. (2013). International Union of Basic and Clinical Pharmacology. [corrected]. LXXXVII. Complement peptide C5a, C4a, and C3a receptors. Pharmacol Rev 65, 500–543. 10.1124/pr.111.005223.

7. Ames, R.S., Li, Y., Sarau, H.M., Nuthulaganti, P., Foley, J.J., Ellis, C., Zeng, Z., Su, K., Jurewicz, A.J., Hertzberg, R.P., et al. (1996). Molecular cloning and characterization of the human anaphylatoxin C3a receptor. J Biol Chem 271, 20231–20234. 10.1074/jbc.271.34.20231.

8. Boulay, F., Mery, L., Tardif, M., Brouchon, L., and Vignais, P. (1991). Expression cloning of a receptor for C5a anaphylatoxin on differentiated HL-60 cells. Biochemistry 30, 2993–2999. 10.1021/bi00226a002.

9. Ohno, M., Hirata, T., Enomoto, M., Araki, T., Ishimaru, H., and Takahashi, T.A. (2000). A putative chemoattractant receptor, C5L2, is expressed in granulocyte and immature dendritic cells, but not in mature dendritic cells. Molecular Immunology 37, 407–412. Doi 10.1016/S0161-5890(00)00067-5.

10. Pandey, S., Kumari, P., Baidya, M., Kise, R., Cao, Y., Dwivedi-Agnihotri, H., Banerjee, R., Li, X.X., Cui, C.S., Lee, J.D., et al. (2021). Intrinsic bias at non-canonical, beta-arrestin-coupled seven transmembrane receptors. Mol Cell 81, 4605–4621 e4611. 10.1016/j.molcel.2021.09.007.

11. Van Lith, L.H., Oosterom, J., Van Elsas, A., and Zaman, G.J. (2009). C5a-stimulated recruitment of beta-arrestin2 to the nonsignaling 7-transmembrane decoy receptor C5L2. J Biomol Screen 14, 1067–1075. 10.1177/1087057109341407.

12. Okinaga, S., Slattery, D., Humbles, A., Zsengeller, Z., Morteau, O., Kinrade, M.B., Brodbeck, R.M., Krause, J.E., Choe, H.R., Gerard, N.P., and Gerard, C. (2003). C5L2, a nonsignaling C5A binding protein. Biochemistry 42, 9406–9415. 10.1021/bi034489v.

13. Gerard, N.P., and Gerard, C. (1991). The chemotactic receptor for human C5a anaphylatoxin. Nature 349, 614–617. 10.1038/349614a0.

14. Füreder, W., Agis, H., Willheim, M., Bankl, H.C., Maier, U., Kishi, K., Müller, M.R., Czerwenka, K., Radaszkiewicz, T., Butterfield, J.H., et al. (1995). Differential expression of complement receptors on human basophils and mast cells. Evidence for mast cell heterogeneity and CD88/C5aR expression on skin mast cells. The Journal of Immunology 155, 3152–3160. 10.4049/jimmunol.155.6.3152.

15. Li, R., Coulthard, L.G., Wu, M.C., Taylor, S.M., and Woodruff, T.M. (2013). C5L2: a controversial receptor of complement anaphylatoxin, C5a. Faseb J 27, 855–864. 10.1096/fj.12-220509.

16. Martin, U., Bock, D., Arseniev, L., Tornetta, M.A., Ames, R.S., Bautsch, W., Köhl, J., Ganser, A., and Klos, A. (1997). The human C3a receptor is expressed on neutrophils and monocytes, but not on B or T lymphocytes. J Exp Med 186, 199–207. 10.1084/jem.186.2.199.

17. Carvelli, J., Demaria, O., Vely, F., Batista, L., Chouaki Benmansour, N., Fares, J., Carpentier, S., Thibult, M.L., Morel, A., Remark, R., et al. (2020). Association of COVID-19 inflammation with activation of the C5a-C5aR1 axis. Nature 588, 146–150. 10.1038/s41586-020-2600-6.

18. Guo, R.F., and Ward, P.A. (2005). Role of C5a in inflammatory responses. Annu Rev Immunol 23, 821–852. 10.1146/annurev.immunol.23.021704.115835.

19. Rittirsch, D., Flierl, M.A., Nadeau, B.A., Day, D.E., Huber-Lang, M., Mackay, C.R., Zetoune, F.S., Gerard, N.P., Cianflone, K., Kohl, J., et al. (2008). Functional roles for C5a receptors in sepsis. Nat Med 14, 551–557. 10.1038/nm1753.

20. Ricklin, D., Reis, E.S., and Lambris, J.D. (2016). Complement in disease: a defence system turning offensive. Nat Rev Nephrol 12, 383–401. 10.1038/nrneph.2016.70.

21. Matthews, K.W., Mueller-Ortiz, S.L., and Wetsel, R.A. (2004). Carboxypeptidase N: a pleiotropic regulator of inflammation. Mol Immunol 40, 785–793. 10.1016/j.molimm.2003.10.002.

22. Bokisch, V.A., and Muller-Eberhard, H.J. (1970). Anaphylatoxin inactivator of human plasma: its isolation and characterization as a carboxypeptidase. J Clin Invest 49, 2427–2436. 10.1172/JCI106462.

23. Croker, D.E., Halai, R., Fairlie, D.P., and Cooper, M.A. (2013). C5a, but not C5a-des Arg, induces upregulation of heteromer formation between complement C5a receptors C5aR and C5L2. Immunol Cell Biol 91, 625–633. 10.1038/icb.2013.48.

24. Croker, D.E., Halai, R., Fairlie, D.P., and Cooper, M.A. (2012). Ligand-induced dimerisation of the complement C5aR and C5L2 receptors by C5a but not C5a-des Arg. Immunobiology 217, 1181–1182. 10.1016/j.imbio.2012.08.152.

25. Wilken, H.C., Götze, O., Werfel, T., and Zwirner, J. (1999). C3a(desArg) does not bind to and signal through the human C3a receptor. Immunol Lett 67, 141–145. 10.1016/s0165-2478(99)00002-4.

26. Polley, M.J., and Nachman, R.L. (1983). Human platelet activation by C3a and C3a des-arg. J Exp Med 158, 603–615. 10.1084/jem.158.2.603.

27. Sayah, S., Jauneau, A.C., Patte, C., Tonon, M.C., Vaudry, H., and Fontaine, M. (2003). Two different transduction pathways are activated by C3a and C5a anaphylatoxins on astrocytes. Brain Res Mol Brain Res 112, 53–60. 10.1016/s0169-328x(03)00046-9.

28. Hannedouche, S., Beck, V., Leighton-Davies, J., Beibel, M., Roma, G., Oakeley, E.J., Lannoy, V., Bernard, J., Hamon, J., Barbieri, S., et al. (2013). Identification of the C3a receptor (C3AR1) as the target of the VGF-derived peptide TLQP-21 in rodent cells. J Biol Chem 288, 27434–27443. 10.1074/jbc.M113.497214.

29. Cero, C., Vostrikov, V.V., Verardi, R., Severini, C., Gopinath, T., Braun, P.D., Sassano, M.F., Gurney, A., Roth, B.L., Vulchanova, L., et al. (2014). The TLQP-21 peptide activates the G-protein-coupled receptor C3aR1 via a folding-upon-binding mechanism. Structure 22, 1744–1753. 10.1016/j.str.2014.10.001.

30. Cero, C., Razzoli, M., Han, R., Sahu, B.S., Patricelli, J., Guo, Z., Zaidman, N.A., Miles, J.M., O’Grady, S.M., and Bartolomucci, A. (2017). The neuropeptide TLQP-21 opposes obesity via C3aR1-mediated enhancement of adrenergic-induced lipolysis. Mol Metab 6, 148–158. 10.1016/j.molmet.2016.10.005.

31. Possenti, R., Muccioli, G., Petrocchi, P., Cero, C., Cabassi, A., Vulchanova, L., Riedl, M.S., Manieri, M., Frontini, A., Giordano, A., et al. (2012). Characterization of a novel peripheral pro-lipolytic mechanism in mice: role of VGF-derived peptide TLQP-21. Biochem J 441, 511–522. 10.1042/BJ20111165.

32. Sahu, B.S., Razzoli, M., McGonigle, S., Pallais, J.P., Nguyen, M.E., Sadahiro, M., Jiang, C., Lin, W.J., Kelley, K.A., Rodriguez, P., et al. (2023). Targeted and selective knockout of the TLQP-21 neuropeptide unmasks its unique role in energy homeostasis. Mol Metab 76, 101781. 10.1016/j.molmet.2023.101781.

33. Sahu, B.S., Rodriguez, P., Nguyen, M.E., Han, R., Cero, C., Razzoli, M., Piaggi, P., Laskowski, L.J., Pavlicev, M., Muglia, L., et al. (2019). Peptide/Receptor Co-evolution Explains the Lipolytic Function of the Neuropeptide TLQP-21. Cell Rep 28, 2567–2580 e2566. 10.1016/j.celrep.2019.07.101.

34. El Gaamouch, F., Audrain, M., Lin, W.J., Beckmann, N., Jiang, C., Hariharan, S., Heeger, P.S., Schadt, E.E., Gandy, S., Ehrlich, M.E., and Salton, S.R. (2020). VGF-derived peptide TLQP-21 modulates microglial function through C3aR1 signaling pathways and reduces neuropathology in 5xFAD mice. Mol Neurodegener 15, 4. 10.1186/s13024-020-0357-x.

35. Cho, K., Jang, Y.J., Lee, S.J., Jeon, Y.N., Shim, Y.L., Lee, J.Y., Lim, D.S., Kim, D.H., and Yoon, S.Y. (2020). TLQP-21 mediated activation of microglial BV2 cells promotes clearance of extracellular fibril amyloid-β. Biochem Biophys Res Commun 524, 764–771. 10.1016/j.bbrc.2020.01.111.

36. Rodriguez, P., Laskowski, L.J., Pallais, J.P., Bock, H.A., Cavalco, N.G., Anderson, E.I., Calkins, M.M., Razzoli, M., Sham, Y.Y., McCorvy, J.D., and Bartolomucci, A. (2024). Functional profiling of the G protein-coupled receptor C3aR1 reveals ligand-mediated biased agonism. J Biol Chem 300, 105549. 10.1016/j.jbc.2023.105549.

37. Bartolomucci, A., La Corte, G., Possenti, R., Locatelli, V., Rigamonti, A.E., Torsello, A., Bresciani, E., Bulgarelli, I., Rizzi, R., Pavone, F., et al. (2006). TLQP-21, a VGF-derived peptide, increases energy expenditure and prevents the early phase of diet-induced obesity. Proc Natl Acad Sci U S A 103, 14584–14589. 10.1073/pnas.0606102103.

38. Scully, C.C., Blakeney, J.S., Singh, R., Hoang, H.N., Abbenante, G., Reid, R.C., and Fairlie, D.P. (2010). Selective hexapeptide agonists and antagonists for human complement C3a receptor. J Med Chem 53, 4938–4948. 10.1021/jm1003705.

39. Finch, A.M., Wong, A.K., Paczkowski, N.J., Wadi, S.K., Craik, D.J., Fairlie, D.P., and Taylor, S.M. (1999). Low-molecular-weight peptidic and cyclic antagonists of the receptor for the complement factor C5a. J Med Chem 42, 1965–1974. 10.1021/jm9806594.

40. Finch, A.M., Vogen, S.M., Sherman, S.A., Kirnarsky, L., Taylor, S.M., and Sanderson, S.D. (1997). Biologically active conformer of the effector region of human C5a and modulatory effects of N-terminal receptor binding determinants on activity. J Med Chem 40, 877–884. 10.1021/jm960727r.

41. Ember, J.A., Johansen, N.L., and Hugli, T.E. (1991). Designing synthetic superagonists of C3a anaphylatoxin. Biochemistry 30, 3603–3612. 10.1021/bi00229a003.

42. Karuturi, B.V., Tallapaka, S.B., Phillips, J.A., Sanderson, S.D., and Vetro, J.A. (2015). Preliminary evidence that the novel host-derived immunostimulant EP67 can act as a mucosal adjuvant. Clin Immunol 161, 251–259. 10.1016/j.clim.2015.06.006.

43. Hung, C.Y., Hurtgen, B.J., Bellecourt, M., Sanderson, S.D., Morgan, E.L., and Cole, G.T. (2012). An agonist of human complement fragment C5a enhances vaccine immunity against Coccidioides infection. Vaccine 30, 4681–4690. 10.1016/j.vaccine.2012.04.084.

44. Phillips, J.A., Morgan, E.L., Dong, Y., Cole, G.T., McMahan, C., Hung, C.Y., and Sanderson, S.D. (2009). Single-step conjugation of bioactive peptides to proteins via a self-contained succinimidyl bis-arylhydrazone. Bioconjug Chem 20, 1950–1957. 10.1021/bc9002794.

45. Morgan, E.L., Morgan, B.N., Stein, E.A., Vitrs, E.L., Thoman, M.L., Sanderson, S.D., and Phillips, J.A. (2009). Enhancement of in vivo and in vitro immune functions by a conformationally biased, response-selective agonist of human C5a: implications for a novel adjuvant in vaccine design. Vaccine 28, 463–469. 10.1016/j.vaccine.2009.10.029.

46. Tifrea, D.F., Pal, S., Le Bon, C., Giusti, F., Popot, J.L., Cocco, M.J., Zoonens, M., and de la Maza, L.M. (2018). Co-delivery of amphipol-conjugated adjuvant with antigen, and adjuvant combinations, enhance immune protection elicited by a membrane protein-based vaccine against a mucosal challenge with Chlamydia. Vaccine 36, 6640–6649. 10.1016/j.vaccine.2018.09.055.

47. Parriott, J.E., Stewart, J.P., Smith, D.D., Curran, S.M., Bauer, C.D., Wyatt, T.A., Phillips, J.A., Lyden, E., Thiele, G.M., and Vetro, J.A. (2022). Surface Modification of Biodegradable Microparticles with the Novel Host-Derived Immunostimulant CPDI-02 Significantly Increases Short-Term and Long-Term Mucosal and Systemic Antibodies against Encapsulated Protein Antigen in Young Naive Mice after Respiratory Immunization. Pharmaceutics 14. 10.3390/pharmaceutics14091843.

48. Cavaco, C.K., Patras, K.A., Zlamal, J.E., Thoman, M.L., Morgan, E.L., Sanderson, S.D., and Doran, K.S. (2013). A novel C5a-derived immunobiotic peptide reduces Streptococcus agalactiae colonization through targeted bacterial killing. Antimicrob Agents Chemother 57, 5492–5499. 10.1128/AAC.01590-13.

49. Sheen, T.R., Cavaco, C.K., Ebrahimi, C.M., Thoman, M.L., Sanderson, S.D., Morgan, E.L., and Doran, K.S. (2011). Control of methicillin resistant Staphylococcus aureus infection utilizing a novel immunostimulatory peptide. Vaccine 30, 9–13. 10.1016/j.vaccine.2011.10.054.

50. Hanke, M.L., Heim, C.E., Angle, A., Sanderson, S.D., and Kielian, T. (2013). Targeting macrophage activation for the prevention and treatment of Staphylococcus aureus biofilm infections. J Immunol 190, 2159–2168. 10.4049/jimmunol.1202348.

51. Rowley, J.A., Reid, R.C., Poon, E.K.Y., Wu, K.C., Lim, J., Lohman, R.J., Hamidon, J.K., Yau, M.K., Halili, M.A., Durek, T., et al. (2020). Potent Thiophene Antagonists of Human Complement C3a Receptor with Anti-Inflammatory Activity. J Med Chem 63, 529–541. 10.1021/acs.jmedchem.9b00927.

52. Ames, R.S., Lee, D., Foley, J.J., Jurewicz, A.J., Tornetta, M.A., Bautsch, W., Settmacher, B., Klos, A., Erhard, K.F., Cousins, R.D., et al. (2001). Identification of a selective nonpeptide antagonist of the anaphylatoxin C3a receptor that demonstrates antiinflammatory activity in animal models. J Immunol 166, 6341–6348. 10.4049/jimmunol.166.10.6341.

53. Tang, J., Maihemuti, N., Fang, Y., Tan, J., Jia, M., Mu, Q., Huang, K., Gan, H., and Zhao, J. (2024). JR14a: A novel antagonist of C3aR attenuates neuroinflammation in cerebral ischemia-reperfusion injury. Brain Res Bull 213, 110986. 10.1016/j.brainresbull.2024.110986.

54. Proctor, L.M., Arumugam, T.V., Shiels, I., Reid, R.C., Fairlie, D.P., and Taylor, S.M. (2004). Comparative anti-inflammatory activities of antagonists to C3a and C5a receptors in a rat model of intestinal ischaemia/reperfusion injury. Br J Pharmacol 142, 756–764. 10.1038/sj.bjp.0705819.

55. Hutamekalin, P., Takeda, K., Tani, M., Tsuga, Y., Ogawa, N., Mizutani, N., and Yoshino, S. (2010). Effect of the C3a-receptor antagonist SB 290157 on anti-OVA polyclonal antibody-induced arthritis. J Pharmacol Sci 112, 56–63. 10.1254/jphs.09180fp.

56. Mathieu, M.C., Sawyer, N., Greig, G.M., Hamel, M., Kargman, S., Ducharme, Y., Lau, C.K., Friesen, R.W., O’Neill, G.P., Gervais, F.G., and Therien, A.G. (2005). The C3a receptor antagonist SB 290157 has agonist activity. Immunol Lett 100, 139–145. 10.1016/j.imlet.2005.03.003.

57. Li, X.X., Kumar, V., Clark, R.J., Lee, J.D., and Woodruff, T.M. (2020). The “C3aR Antagonist” SB290157 is a Partial C5aR2 Agonist. Front Pharmacol 11, 591398. 10.3389/fphar.2020.591398.

58. Luo, P., Xin, W., Guo, S., Li, X., Zhang, Q., Xu, Y., He, X., Wang, Y., Fan, W., Yuan, Q., et al. (2025). Structural insights into the agonist activity of the nonpeptide modulator JR14a on C3aR. Cell Discov 11, 7. 10.1038/s41421-024-00765-x.

59. Kim, J., Ko, S.B., Choi, C., Bae, J., Byeon, H., Seok, C., and Choi, H.J. (2025). Structural insights into small-molecule agonist recognition and activation of complement receptor C3aR. Embo Journal 44, 2803–2826. 10.1038/s44318-025-00429-w.

60. Baidya, M., Chaturvedi, M., Dwivedi-Agnihotri, H., Ranjan, A., Devost, D., Namkung, Y., Stepniewski, T.M., Pandey, S., Baruah, M., Panigrahi, B., et al. (2022). Allosteric modulation of GPCR-induced beta-arrestin trafficking and signaling by a synthetic intrabody. Nat Commun 13, 4634. 10.1038/s41467-022-32386-x.

61. Kumar, B.A., Kumari, P., Sona, C., and Yadav, P.N. (2017). GloSensor assay for discovery of GPCR-selective ligands. Methods Cell Biol 142, 27–50. 10.1016/bs.mcb.2017.07.012.

62. Baidya, M., Kumari, P., Dwivedi-Agnihotri, H., Pandey, S., Chaturvedi, M., Stepniewski, T.M., Kawakami, K., Cao, Y., Laporte, S.A., Selent, J., et al. (2020). Key phosphorylation sites in GPCRs orchestrate the contribution of beta-Arrestin 1 in ERK1/2 activation. EMBO Rep 21, e49886. 10.15252/embr.201949886.

63. Reis, E.S., Chen, H., Sfyroera, G., Monk, P.N., Kohl, J., Ricklin, D., and Lambris, J.D. (2012). C5a receptor-dependent cell activation by physiological concentrations of desarginated C5a: insights from a novel label-free cellular assay. J Immunol 189, 4797–4805. 10.4049/jimmunol.1200834.

64. Li, X.X., Lee, J.D., Lee, H.S., Clark, R.J., and Woodruff, T.M. (2023). TLQP-21 is a low potency partial C3aR activator on human primary macrophages. Front Immunol 14, 1086673. 10.3389/fimmu.2023.1086673.

65. Rizzi, R., Bartolomucci, A., Moles, A., D’Amato, F., Sacerdote, P., Levi, A., La Corte, G., Ciotti, M.T., Possenti, R., and Pavone, F. (2008). The VGF-derived peptide TLQP-21: a new modulatory peptide for inflammatory pain. Neurosci Lett 441, 129–133. 10.1016/j.neulet.2008.06.018.

66. Bartolomucci, A., Possenti, R., Mahata, S.K., Fischer-Colbrie, R., Loh, Y.P., and Salton, S.R. (2011). The extended granin family: structure, function, and biomedical implications. Endocr Rev 32, 755–797. 10.1210/er.2010-0027.

67. Jethwa, P.H., Warner, A., Nilaweera, K.N., Brameld, J.M., Keyte, J.W., Carter, W.G., Bolton, N., Bruggraber, M., Morgan, P.J., Barrett, P., and Ebling, F.J. (2007). VGF-derived peptide, TLQP-21, regulates food intake and body weight in Siberian hamsters. Endocrinology 148, 4044–4055. 10.1210/en.2007-0038.

68. Mamane, Y., Chung Chan, C., Lavallee, G., Morin, N., Xu, L.J., Huang, J., Gordon, R., Thomas, W., Lamb, J., Schadt, E.E., et al. (2009). The C3a anaphylatoxin receptor is a key mediator of insulin resistance and functions by modulating adipose tissue macrophage infiltration and activation. Diabetes 58, 2006–2017. 10.2337/db09-0323.

69. Halai, R., Bellows-Peterson, M.L., Branchett, W., Smadbeck, J., Kieslich, C.A., Croker, D.E., Cooper, M.A., Morikis, D., Woodruff, T.M., Floudas, C.A., and Monk, P.N. (2014). Derivation of ligands for the complement C3a receptor from the C-terminus of C5a. Eur J Pharmacol 745, 176–181. 10.1016/j.ejphar.2014.10.041.

70. Gorman, D.M., Li, X.X., Lee, J.D., Fung, J.N., Cui, C.S., Lee, H.S., Rolfe, B.E., Woodruff, T.M., and Clark, R.J. (2021). Development of Potent and Selective Agonists for Complement C5a Receptor 1 with In Vivo Activity. J Med Chem 64, 16598–16608. 10.1021/acs.jmedchem.1c01174.

71. Woodruff, T.M., Strachan, A.J., Sanderson, S.D., Monk, P.N., Wong, A.K., Fairlie, D.P., and Taylor, S.M. (2001). Species dependence for binding of small molecule agonist and antagonists to the C5a receptor on polymorphonuclear leukocytes. Inflammation 25, 171–177. 10.1023/a:1011036414353.

72. Kawai, M., Quincy, D.A., Lane, B., Mollison, K.W., Or, Y.S., Luly, J.R., and Carter, G.W. (1992). Structure-function studies in a series of carboxyl-terminal octapeptide analogues of anaphylatoxin C5a. J Med Chem 35, 220–223. 10.1021/jm00080a004.

73. Siciliano, S.J., Rollins, T.E., DeMartino, J., Konteatis, Z., Malkowitz, L., Van Riper, G., Bondy, S., Rosen, H., and Springer, M.S. (1994). Two-site binding of C5a by its receptor: an alternative binding paradigm for G protein-coupled receptors. Proc Natl Acad Sci U S A 91, 1214–1218. 10.1073/pnas.91.4.1214.

74. Taylor, S.M., Sherman, S.A., Kirnarsky, L., and Sanderson, S.D. (2001). Development of response-selective agonists of human C5a anaphylatoxin: conformational, biological, and therapeutic considerations. Curr Med Chem 8, 675–684. 10.2174/0929867013373156.

75. Wang, Y., Liu, W., Xu, Y., He, X., Yuan, Q., Luo, P., Fan, W., Zhu, J., Zhang, X., Cheng, X., et al. (2023). Revealing the signaling of complement receptors C3aR and C5aR1 by anaphylatoxins. Nat Chem Biol 19, 1351–1360. 10.1038/s41589-023-01339-w.

76. Feng, Y., Zhao, C., Deng, Y., Wang, H., Ma, L., Liu, S., Tian, X., Wang, B., Bin, Y., Chen, P., et al. (2023). Mechanism of activation and biased signaling in complement receptor C5aR1. Cell Res 33, 312–324. 10.1038/s41422-023-00779-2.

77. Yadav, M.K., Maharana, J., Yadav, R., Saha, S., Sarma, P., Soni, C., Singh, V., Saha, S., Ganguly, M., Li, X.X., et al. (2023). Molecular basis of anaphylatoxin binding, activation, and signaling bias at complement receptors. Cell 186, 4956–4973.e4921. 10.1016/j.cell.2023.09.020.

78. Yadav, M.K., Maharana, J., Yadav, R., Saha, S., Sarma, P., Soni, C., Singh, V., Saha, S., Ganguly, M., Li, X.X., et al. (2023). Molecular basis of anaphylatoxin binding, activation, and signaling bias at complement receptors. Cell 186, 4956–4973 e4921. 10.1016/j.cell.2023.09.020.

79. Gedam, M., Comerota, M.M., Propson, N.E., Chen, T., Jin, F., Wang, M.C., and Zheng, H. (2023). Complement C3aR depletion reverses HIF-1alpha-induced metabolic impairment and enhances microglial response to Abeta pathology. J Clin Invest 133. 10.1172/JCI167501.

80. Isaikina, P., Petrovic, I., Jakob, R.P., Sarma, P., Ranjan, A., Baruah, M., Panwalkar, V., Maier, T., Shukla, A.K., and Grzesiek, S. (2023). A key GPCR phosphorylation motif discovered in arrestin2⋅CCR5 phosphopeptide complexes. Mol Cell 83, 2108–2121 e2107. 10.1016/j.molcel.2023.05.002.

81. Maharana, J., Sarma, P., Yadav, M.K., Saha, S., Singh, V., Saha, S., Chami, M., Banerjee, R., and Shukla, A.K. (2023). Structural snapshots uncover a key phosphorylation motif in GPCRs driving beta-arrestin activation. Mol Cell 83, 2091–2107 e2097. 10.1016/j.molcel.2023.04.025.

82. Maharana, J., Sano, F.K., Sarma, P., Yadav, M.K., Duan, L., Stepniewski, T.M., Chaturvedi, M., Ranjan, A., Singh, V., Saha, S., et al. (2024). Molecular insights into atypical modes of βarrestin interaction with seven transmembrane receptors. Science 383, 101–108. 10.1126/science.adj3347.

83. West, E.E., and Kemper, C. (2023). Complosome - the intracellular complement system. Nat Rev Nephrol 19, 426–439. 10.1038/s41581-023-00704-1.

84. Kawakami, K., Yanagawa, M., Hiratsuka, S., Yoshida, M., Ono, Y., Hiroshima, M., Ueda, M., Aoki, J., Sako, Y., and Inoue, A. (2022). Heterotrimeric Gq proteins act as a switch for GRK5/6 selectivity underlying beta-arrestin transducer bias. Nat Commun 13, 487. 10.1038/s41467-022-28056-7.

85. Dijon, N.C., Nesheva, D.N., and Holliday, N.D. (2021). Luciferase Complementation Approaches to Measure GPCR Signaling Kinetics and Bias. In G Protein-Coupled Receptor Screening Assays: Methods and Protocols, S.A.M. Martins, and D.M.F. Prazeres, eds. (Springer US), pp. 249-274. 10.1007/978-1-0716-1221-7_17.

86. Inoue, A., Raimondi, F., Kadji, F.M.N., Singh, G., Kishi, T., Uwamizu, A., Ono, Y., Shinjo, Y., Ishida, S., Arang, N., et al. (2019). Illuminating G-Protein-Coupling Selectivity of GPCRs. Cell 177, 1933–1947 e1925. 10.1016/j.cell.2019.04.044.

87. Dwivedi-Agnihotri, H., Sarma, P., Deeksha, S., Kawakami, K., Inoue, A., and Shukla, A.K. (2022). Chapter 12 - An intrabody sensor to monitor conformational activation of β-arrestins. In Methods in Cell Biology, A.K. Shukla, ed. (Academic Press), pp. 267–278. 10.1016/bs.mcb.2021.12.023.

88. Pandey, S., Roy, D., and Shukla, A.K. (2019). Measuring surface expression and endocytosis of GPCRs using whole-cell ELISA. Methods Cell Biol 149, 131–140. 10.1016/bs.mcb.2018.09.014.

89. Quek, H., Cuni-Lopez, C., Stewart, R., Lim, Y.C., Roberts, T.L., and White, A.R. (2022). A robust approach to differentiate human monocyte-derived microglia from peripheral blood mononuclear cells. STAR Protoc 3, 101747. 10.1016/j.xpro.2022.101747.

90. Bajic, G., Yatime, L., Klos, A., and Andersen, G.R. (2013). Human C3a and C3a desArg anaphylatoxins have conserved structures, in contrast to C5a and C5a desArg. Protein Sci 22, 204–212. 10.1002/pro.2200.

91. Nehme, R., Carpenter, B., Singhal, A., Strege, A., Edwards, P.C., White, C.F., Du, H., Grisshammer, R., and Tate, C.G. (2017). Mini-G proteins: Novel tools for studying GPCRs in their active conformation. PLoS One 12, e0175642. 10.1371/journal.pone.0175642.

92. Carpenter, B., and Tate, C.G. (2017). Expression and Purification of Mini G Proteins from Escherichia coli. Bio Protoc 7. 10.21769/BioProtoc.2235.

93. Hong, C., Byrne, N.J., Zamlynny, B., Tummala, S., Xiao, L., Shipman, J.M., Partridge, A.T., Minnick, C., Breslin, M.J., Rudd, M.T., et al. (2021). Structures of active-state orexin receptor 2 rationalize peptide and small-molecule agonist recognition and receptor activation. Nat Commun 12, 815. 10.1038/s41467-021-21087-6.

94. Saha, S., Sano, F.K., Sharma, S., Ganguly, M., Mishra, S., Dalal, A., Akasaka, H., Kobayashi, T.A., Zaidi, N., Tiwari, D., et al. (2025). Molecular basis of promiscuous chemokine binding and structural mimicry at the C-X-C chemokine receptor, CXCR2. Mol Cell 85, 976–988 e979. 10.1016/j.molcel.2025.01.024.

95. Saha, S., Sano, F.K., Sharma, S., Ganguly, M., Dalal, A., Mishra, S., Tiwari, D., Akasaka, H., Kobayashi, T.A., Roy, N., et al. (2025). Structural visualization of small molecule recognition by CXCR3 uncovers dual-agonism in the CXCR3-CXCR7 system. Nat Commun 16, 3047. 10.1038/s41467-025-58264-w.

96. Saha, S., Khanppnavar, B., Maharana, J., Kim, H., Carino, C.M.C., Daly, C., Houston, S., Sharma, S., Zaidi, N., Dalal, A., et al. (2024). Molecular mechanism of distinct chemokine engagement and functional divergence of the human Duffy antigen receptor. Cell 187, 4751–4769 e4725. 10.1016/j.cell.2024.07.005.

97. Ghosh, E., Srivastava, A., Baidya, M., Kumari, P., Dwivedi, H., Nidhi, K., Ranjan, R., Dogra, S., Koide, A., Yadav, P.N., et al. (2017). A synthetic intrabody-based selective and generic inhibitor of GPCR endocytosis. Nat Nanotechnol 12, 1190-+. 10.1038/Nnano.2017.188.

98. Yadav, M.K., Singh, V., Saha, S., and Shukla, A.K. (2023). A streamlined protocol for expression and purification of wild-type beta-arrestins. Methods Enzymol 682, 465–475. 10.1016/bs.mie.2022.12.006.

99. Yadav, M.K., Sarma, P., Maharana, J., Ganguly, M., Mishra, S., Zaidi, N., Dalal, A., Singh, V., Saha, S., Mahajan, G., et al. (2024). Structure-guided engineering of biased-agonism in the human niacin receptor via single amino acid substitution. Nature Communications 15, 1939. 10.1038/s41467-024-46239-2.

100. Zivanov, J., Nakane, T., Forsberg, B.O., Kimanius, D., Hagen, W.J., Lindahl, E., and Scheres, S.H. (2018). New tools for automated high-resolution cryo-EM structure determination in RELION-3. Elife 7. 10.7554/eLife.42166.

101. Zivanov, J., Nakane, T., and Scheres, S.H.W. (2020). Estimation of high-order aberrations and anisotropic magnification from cryo-EM data sets in RELION-3.1. IUCrJ 7, 253–267. 10.1107/S2052252520000081.

102. Zivanov, J., Oton, J., Ke, Z., von Kugelgen, A., Pyle, E., Qu, K., Morado, D., Castano-Diez, D., Zanetti, G., Bharat, T.A.M., et al. (2022). A Bayesian approach to single-particle electron cryo-tomography in RELION-4.0. Elife 11. 10.7554/eLife.83724.

103. Punjani, A., Rubinstein, J.L., Fleet, D.J., and Brubaker, M.A. (2017). cryoSPARC: algorithms for rapid unsupervised cryo-EM structure determination. Nat Methods 14, 290–296. 10.1038/nmeth.4169.

104. Cianfrocco, M.A., Wong-Barnum, M., Youn, C., Wagner, R., and Leschziner, A. (2017). COSMIC2: A Science Gateway for Cryo-Electron Microscopy Structure Determination. Practice and Experience in Advanced Research Computing 2017: Sustainability, Success and Impact. Association for Computing Machinery.

105. Waterhouse, A., Bertoni, M., Bienert, S., Studer, G., Tauriello, G., Gumienny, R., Heer, F.T., de Beer, T.A.P., Rempfer, C., Bordoli, L., et al. (2018). SWISS-MODEL: homology modelling of protein structures and complexes. Nucleic Acids Res 46, W296–W303. 10.1093/nar/gky427.

106. Pettersen, E.F., Goddard, T.D., Huang, C.C., Couch, G.S., Greenblatt, D.M., Meng, E.C., and Ferrin, T.E. (2004). UCSF Chimera--a visualization system for exploratory research and analysis. J Comput Chem 25, 1605–1612. 10.1002/jcc.20084.

107. Pettersen, E.F., Goddard, T.D., Huang, C.C., Meng, E.C., Couch, G.S., Croll, T.I., Morris, J.H., and Ferrin, T.E. (2021). UCSF ChimeraX: Structure visualization for researchers, educators, and developers. Protein Sci 30, 70–82. 10.1002/pro.3943.

108. Emsley, P., and Cowtan, K. (2004). Coot: model-building tools for molecular graphics. Acta Crystallogr D Biol Crystallogr 60, 2126–2132. 10.1107/S0907444904019158.

109. Emsley, P., Lohkamp, B., Scott, W.G., and Cowtan, K. (2010). Features and development of Coot. Acta Crystallogr D Biol Crystallogr 66, 486–501. 10.1107/S0907444910007493.

110. Casanal, A., Lohkamp, B., and Emsley, P. (2020). Current developments in Coot for macromolecular model building of Electron Cryo-microscopy and Crystallographic Data. Protein Sci 29, 1069–1078. 10.1002/pro.3791.

111. Adams, P.D., Afonine, P.V., Bunkoczi, G., Chen, V.B., Davis, I.W., Echols, N., Headd, J.J., Hung, L.W., Kapral, G.J., Grosse-Kunstleve, R.W., et al. (2010). PHENIX: a comprehensive Python-based system for macromolecular structure solution. Acta Crystallogr D Biol Crystallogr 66, 213–221. 10.1107/S0907444909052925.

112. Liebschner, D., Afonine, P.V., Baker, M.L., Bunkoczi, G., Chen, V.B., Croll, T.I., Hintze, B., Hung, L.W., Jain, S., McCoy, A.J., et al. (2019). Macromolecular structure determination using X-rays, neutrons and electrons: recent developments in Phenix. Acta Crystallogr D Struct Biol 75, 861–877. 10.1107/S2059798319011471.

