## Supplemental Figures for "Molecular fingerprints of a convergent mechanism orchestrating diverse ligand recognition and species-specific pharmacology at the complement anaphylatoxin receptors"

#### Figure S1

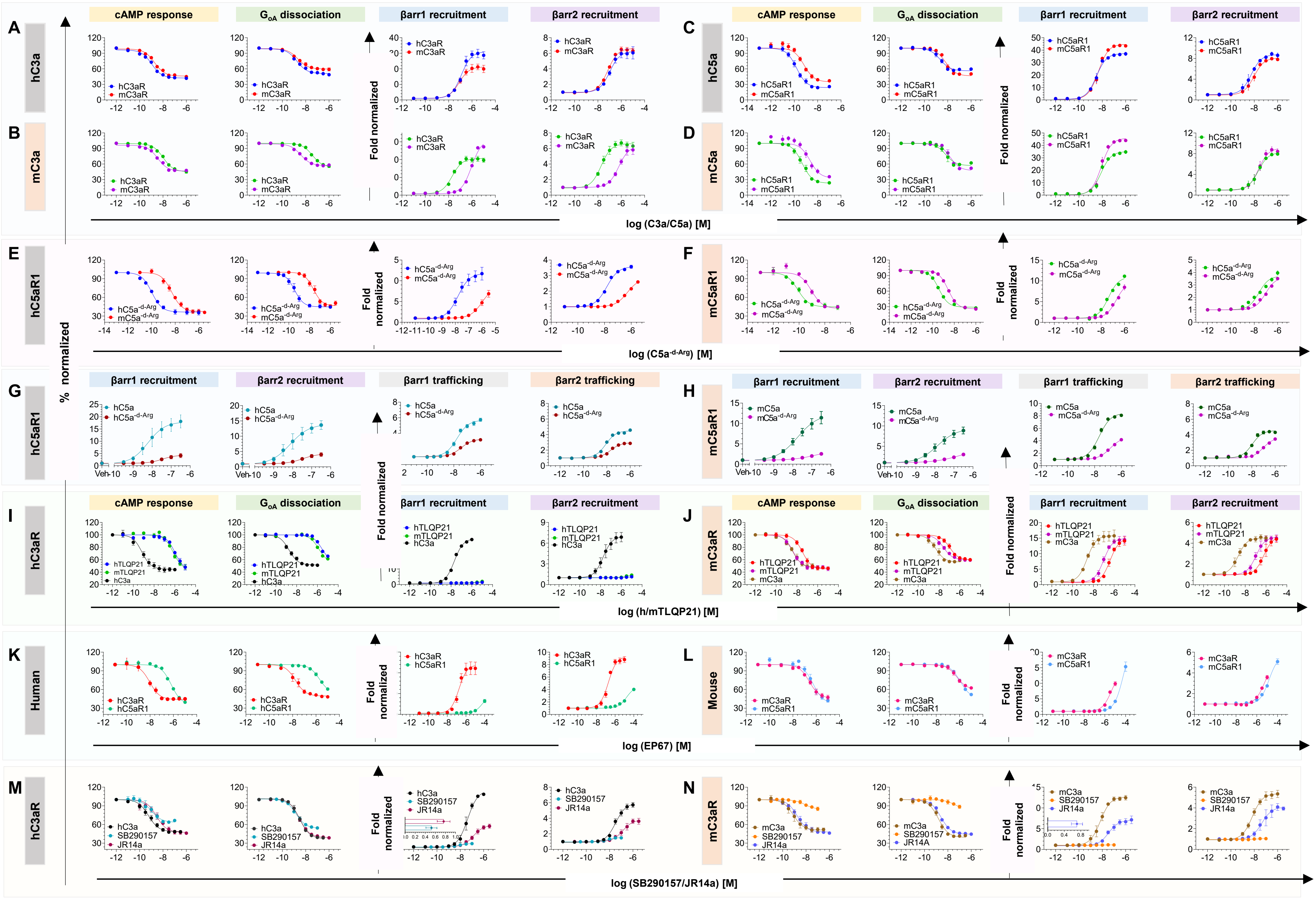
A
B
C
D
E
F
G
H
I
J
K
L
M
N
hC3a
mC3a
hC5a
mC5a
hC5aR1
mC5aR1
hC3aR
mC3aR
Human
Mouse
cAMP response
GoA dissociation
βarr1 recruitment
βarr2 recruitment
βarr1 trafficking
βarr2 trafficking
% normalized
Fold normalized
hC3aR
mC3aR
hC5aR1
mC5aR1
hC5a-d-Arg
mC5a-d-Arg
hC5a
mC5a
hTLQP21
mTLQP21
hC3a
mC3a
SB290157
JR14a
log (C3a/C5a) [M]
log (C5a-d-Arg) [M]
log (h/mTLQP21) [M]
log (EP67) [M]
log (SB290157/JR14a) [M]

**Figure S1. Species-specific pharmacology downstream of complement receptors C3aR and C5aR1.**

(A) hC3a-mediated functional response downstream of hC3aR and mC3aR corresponding to Figure 1 (E and F).

(B) mC3a-mediated functional response downstream of hC3aR and mC3aR corresponding to Figure 1 (E and F).

(C) hC5a mediated functional response downstream of hC5aR1 and mC5aR1 corresponding to Figure 1 (G and H).

(D) mC5a-mediated functional response downstream of hC5aR1 and mC5aR1 corresponding to Figure 1 (G and H).

(E and F) Pharmacological characterisation of C5a$^{\text{-d-Arg}}$ in a cross-species manner at h and mC5aR1 receptors was performed. Data are presented as mean ± SEM (n = 3) and dose-dependent response normalized with the lowest ligand concentration. For cAMP accumulation and GoA dissociation assays, the response at the lowest ligand concentration was set to 100%, while for βarr1/2 recruitment assays, it was normalized to 1.

(G and H) βarr1/2 recruitment was assessed using a NanoBiT-based complementation assay employing the reverse Bit configuration (receptor_LgBiT + SmBiT_βarr1/2) following stimulation with C5a and C5a$^{\text{-d-Arg}}$. Receptor endosomal localisation was evaluated using a NanoBiT-based assay, in which LgBiT was fused to the N-terminus of FYVE and SmBiT to the N-terminus of βarr1/2. Data are presented as mean ± SEM (n = 3) and normalized to the response at the lowest ligand concentration, which was set to 1.

(I and J) Functional response of h and mTLQP21 compared with h and mC3a downstream of hC3aR and mC3aR, respectively (corresponding to Figures 1M and 1N).

(K and L) Comparison between hC3aR vs. C5aR1 and mC3aR vs. C5aR1 (corresponding to Figure 1O–1R).

(M and N) Functional response to SB290157 and JR14a downstream of hC3aR and mC3aR, respectively (corresponding to Figure 1M and 1N). Bias factors for JR14a and SB290157 were calculated relative to the reference ligand C3a using the online bias calculator tool (https://biasedcalculator.shinyapps.io/calc/), based on efficacy and potency data from Gi activation and βarr1/2 recruitment assays.

#### Figure S2

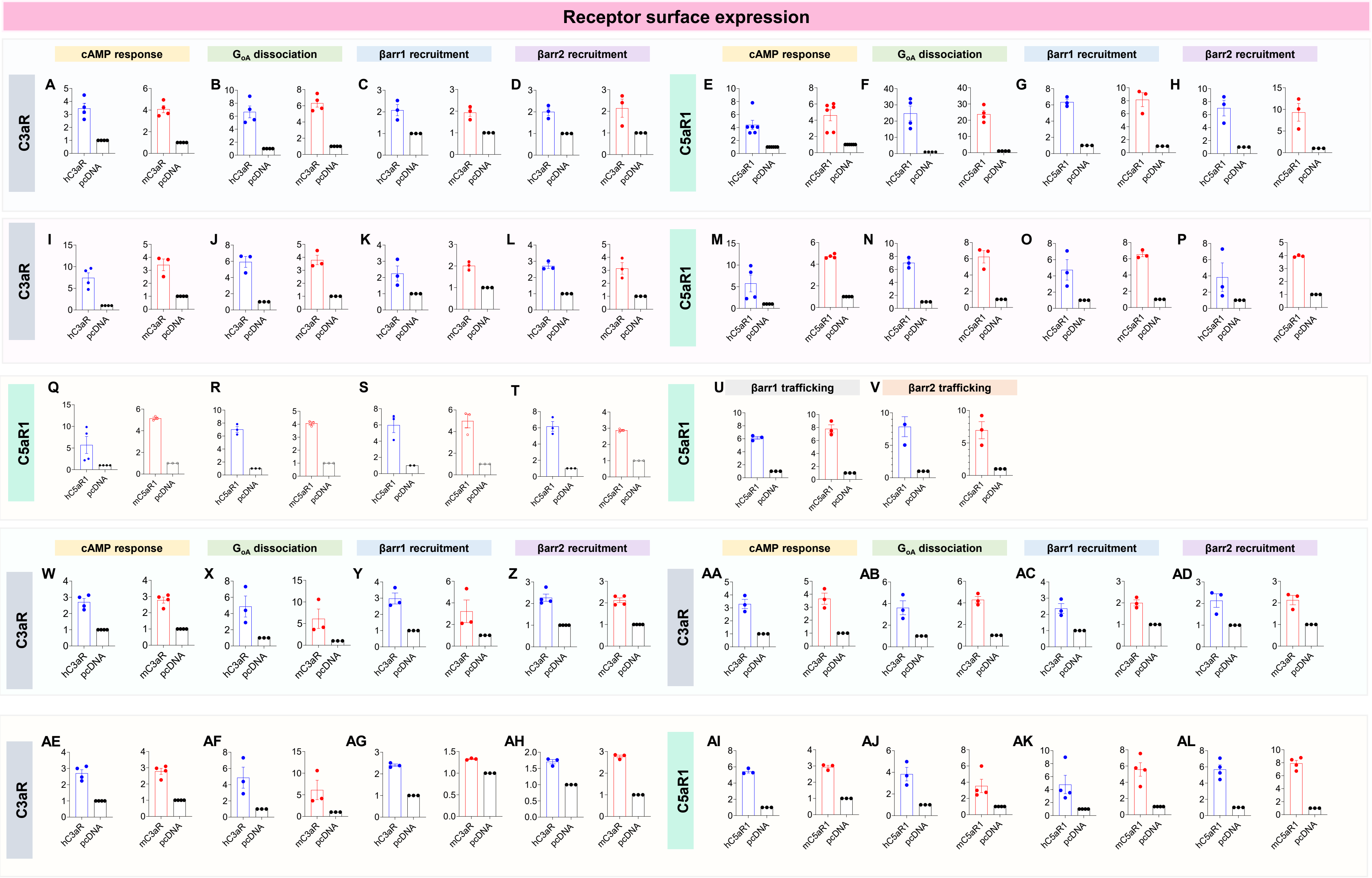

Receptor surface expression
cAMP response
G$_{oA}$ dissociation
βarr1 recruitment
βarr2 recruitment
C3aR
C5aR1
A
B
C
D
E
F
G
H
I
J
K
L
M
N
O
P
Q
R
S
T
U
V
βarr1 trafficking
βarr2 trafficking
W
X
Y
Z
AA
AB
AC
AD
AE
AF
AG
AH
AI
AJ
AK
AL
hC3aR
mC3aR
hC5aR1
mC5aR1
pcDNA

**Figure S2. Receptor surface expression.**

(A-D) Receptor surface expression of h and mC3aR, corresponding to Figure 1 (E, F) and S1 (A, B). Measured via whole-cell ELISA in GloSensor, NanoBiT-based GoA dissociation, and βarr1/2 recruitment assays. Data are mean ± SEM (n = 3–4), normalised to mock-transfected cells (values set to 1).

(E-H) Receptor surface expression of h and mC5aR1, corresponding to Figure 1 (G, H) and S1 (C, D). Measured via whole-cell ELISA in GloSensor, NanoBiT-based GoA dissociation, and βarr1/2 recruitment assays. Data are mean ± SEM (n = 3–4), normalised to mock-transfected cells (values set to 1).

(I-L) Receptor surface expression of h and mC3aR, corresponding to Figure 1 (I, J). Measured via whole-cell ELISA in GloSensor, NanoBiT-based GoA dissociation, and βarr1/2 recruitment assays. Data are mean ± SEM (n = 3–4), normalised to mock-transfected cells (values set to 1).

(M-P) Receptor surface expression of h and mC5aR1, corresponding to Figure 1 (K, L). Measured via whole-cell ELISA in GloSensor, NanoBiT-based GoA dissociation, and βarr1/2 recruitment assays. Data are mean ± SEM (n = 3–4), normalised to mock-transfected cells (values set to 1).

(Q-V) Receptor surface expression of h and mC5aR1, corresponding to Figure S1 (E, H). Measured via whole-cell ELISA in GloSensor, NanoBiT-based GoA dissociation, and βarr1/2 recruitment assays. Data are mean ± SEM (n = 3), normalised to mock-transfected cells (values set to 1).

(W-Z) Receptor surface expression of h and mC3aR, corresponding to Figure 1 (M, N) and S1 (I, J). Measured via whole-cell ELISA in GloSensor, NanoBiT-based GoA dissociation, and βarr1/2 recruitment assays. Data are mean ± SEM (n = 3), normalised to mock-transfected cells (values set to 1).

(AA-AD) Receptor surface expression of h and mC3aR, corresponding to Figure 1 (S, T) and S1 (M, N). Measured via whole-cell ELISA in GloSensor, NanoBiT-based GoA dissociation, and βarr1/2 recruitment assays. Data are mean ± SEM (n = 3), normalised to mock-transfected cells (values set to 1).

(AE-AHReceptor surface expression of h and mC3aR, corresponding to Figure 1 (O, P) and S1 (K, L). Measured via whole-cell ELISA in GloSensor, NanoBiT-based GoA dissociation, and βarr1/2 recruitment assays. Data are mean ± SEM (n = 3), normalised to mock-transfected cells (values set to 1).

(AI-AL) Receptor surface expression of h and mC5aR1, corresponding to Figure 1 (Q, R) and S1 (K, L). Measured via whole-cell ELISA in GloSensor, NanoBiT-based GoA dissociation, and βarr1/2 recruitment assays. Data are mean ± SEM (n = 3-4), normalised to mock-transfected cells (values set to 1).

**Figure S3**

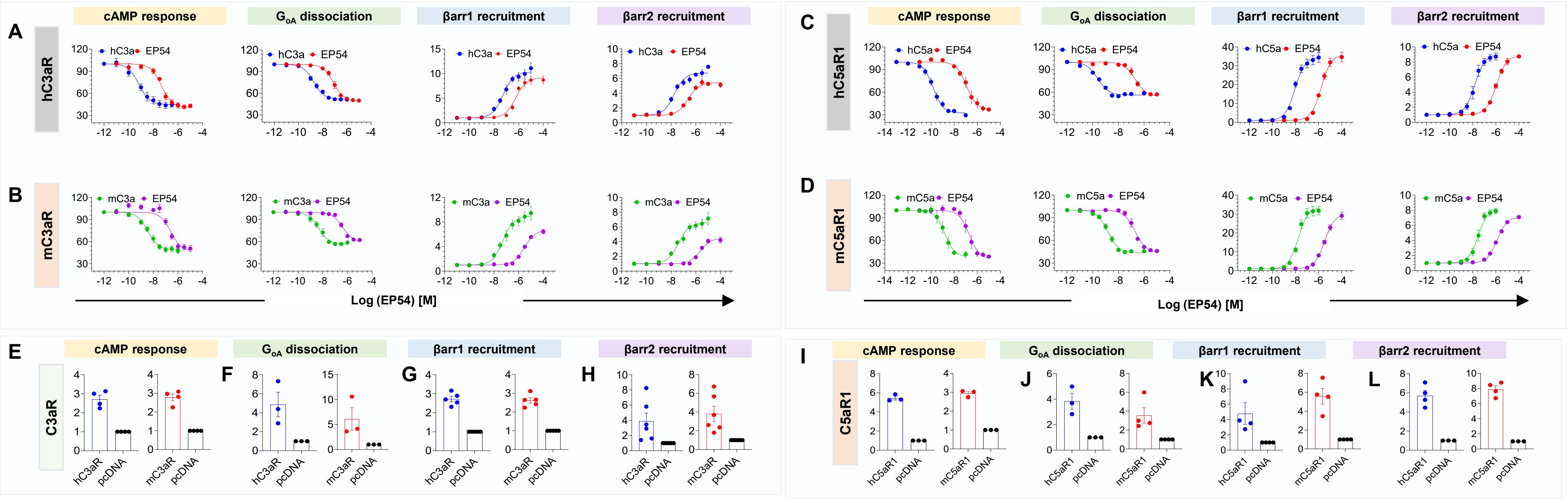

A
hC3aR
cAMP response
$G_{oA}$ dissociation
βarr1 recruitment
βarr2 recruitment
hC3a
EP54
B
mC3aR
mC3a
EP54
Log (EP54) [M]
C
hC5aR1
hC5a
EP54
D
mC5aR1
mC5a
EP54
Log (EP54) [M]
E
C3aR
F
G
H
hC3aR
mC3aR
pcDNA
I
C5aR1
J
K
L
hC5aR1
mC5aR1
pcDNA

**Figure S3. Species-specific pharmacology of EP54 downstream of complement receptors C3aR and C5aR1**

(A-D) Gi activation was assessed using the GloSensor assay, measuring ligand-induced inhibition of forskolin-stimulated cAMP accumulation (mean ± SEM; n=3; normalized with the lowest ligand concentration considered to 100% after forskolin correction). GoA activation was evaluated using a NanoBiT-based heterotrimeric G-protein dissociation assay (mean ± SEM; n=3; normalized with the lowest ligand concentration considered to 100% after basal correction). βarr1/2 recruitment was measured via a NanoBiT-based bystander mode (Receptor + Lg_CAAX + Smβarr1/2) for C3aR and NanoBiT-based direct mode for C5aR1 (mean ± SEM; n=3; normalised with the lowest ligand concentration considered to 1). A and B**)** EP54 mediated functional response downstream of hC3aR and mC3aR; C and D) Functional response downstream of hC5aR1 and mC5aR1 after EP54 stimulation.

(E-L) Receptor surface expression of C3aR and C5aR1 across various assays was measured using a whole-cell ELISA. Data are presented as mean ± SEM (n = 3) and normalized to the signal from mock-transfected cells, which was set to 1.

Figure S4

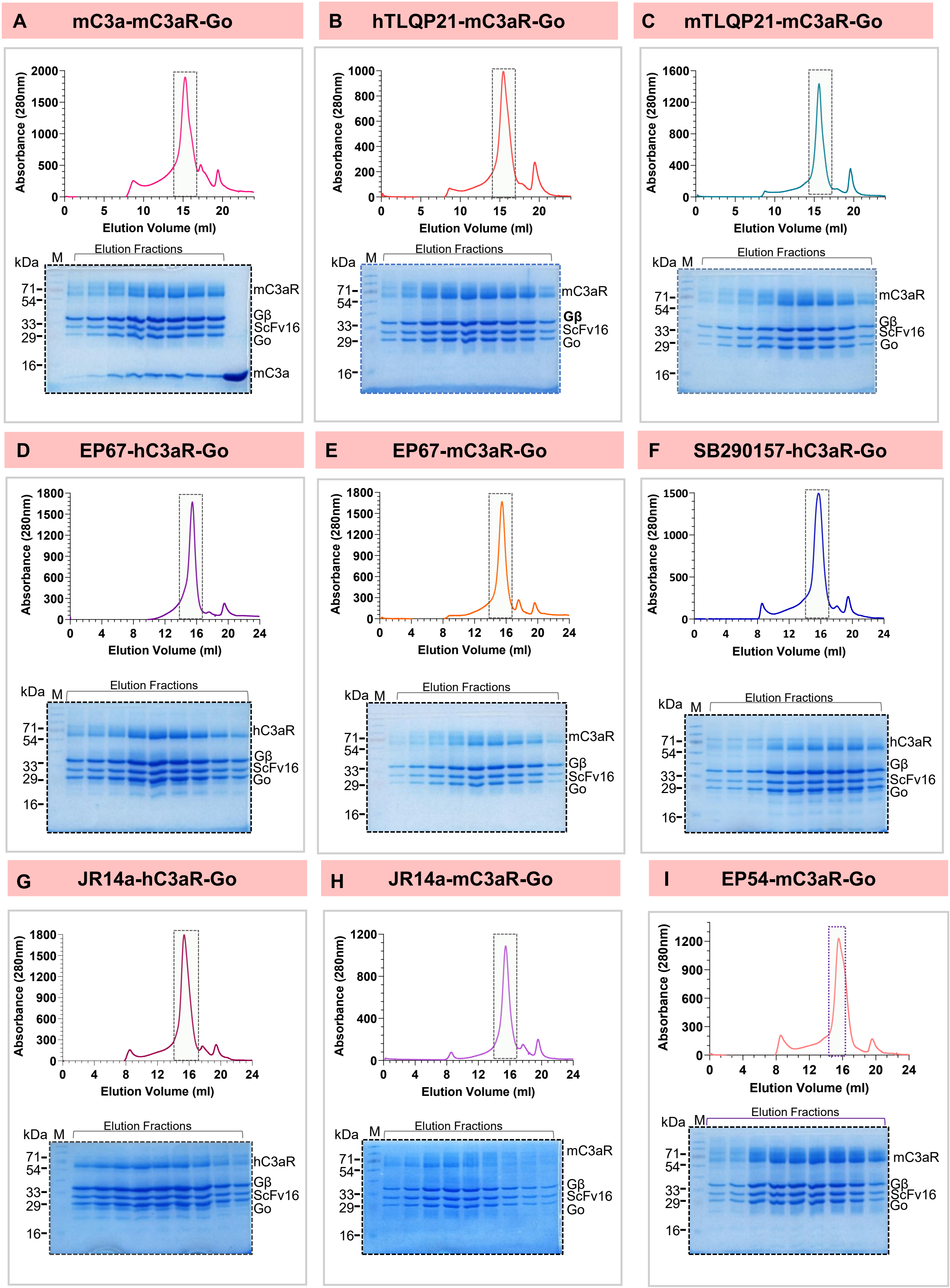


**Figure S4. Purification and preparation of C3aR-G-protein complexes**.

(A-I) Size exclusion chromatograph and SDS-PAGE gel images of mC3a-mC3aR-Go, hTLQP21-mC3aR-Go, mTLQP21-mC3aR-Go, EP67-hC3aR-Go, EP67-mC3aR-Go, SB290157-hC3aR-Go, JR14a-hC3aR, JR14a-mC3aR-Go and EP54-mC3aR-Go complexes respectively.

Figure S5

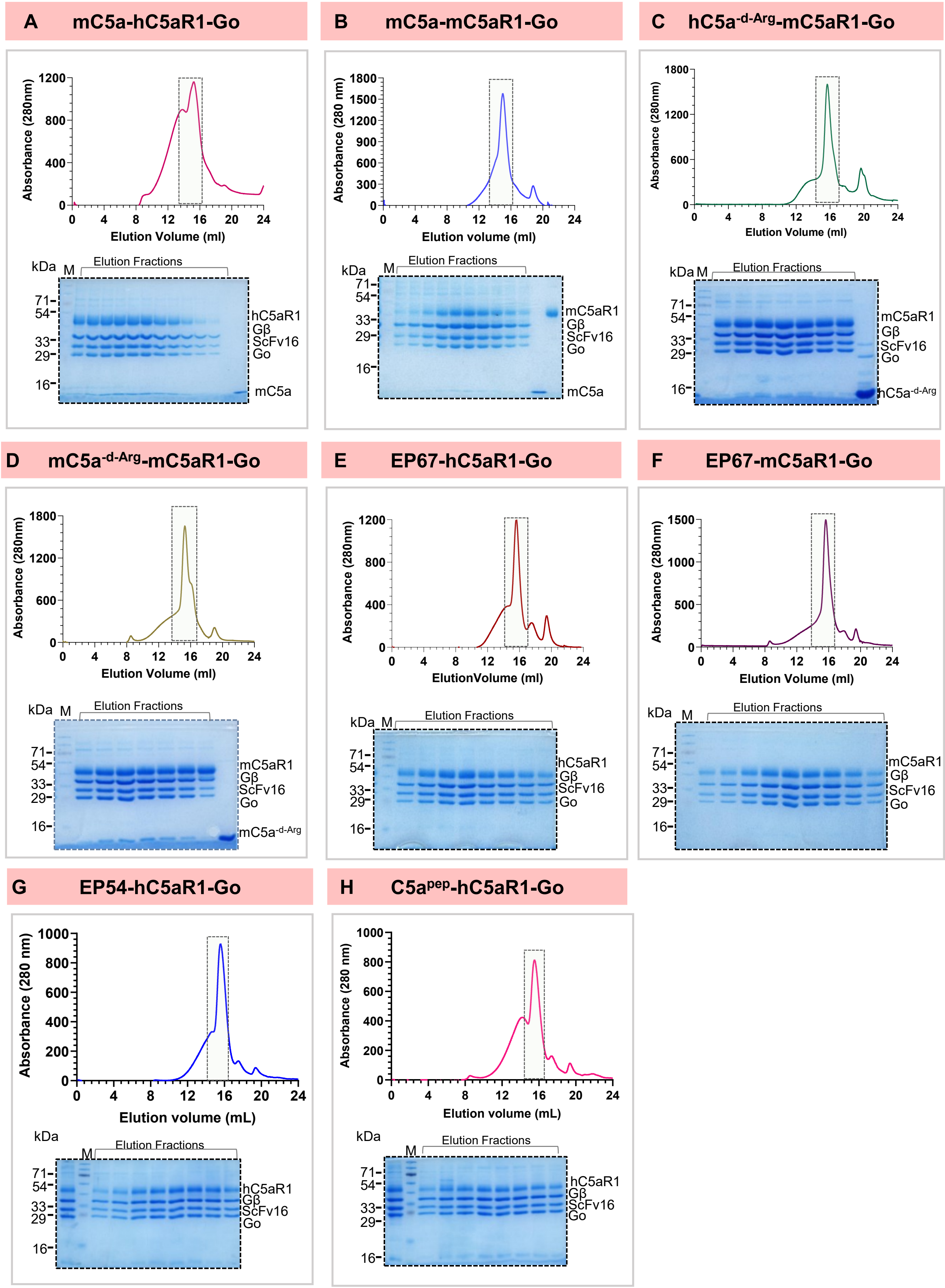


**Figure S5. Purification and preparation of C5aR1-G protein complexes**.

(A-H) Size exclusion chromatograph and SDS-PAGE gel images of mC5a-hC5aR1-Go, mC5a-mC5aR1-Go, hC5a$^{-d-Arg}$-mC5aR1-Go, and mC5a$^{-d-Arg}$-mC5aR1-Go, EP67-hC5aR1-Go, EP67-mC5aR1-Go, EP54-hC5aR1-Go and C5a$^{pep}$-hC5aR1-Go complexes respectively.

Figure S6

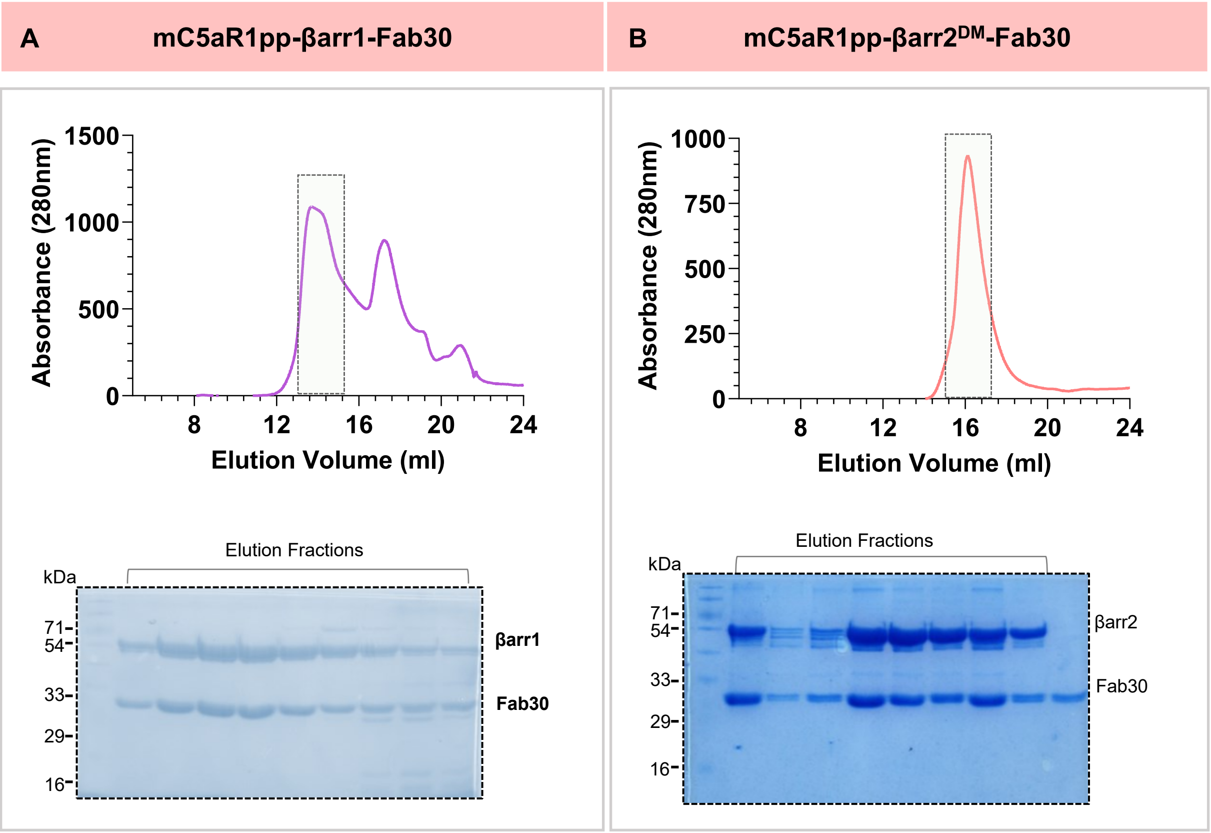


**Figure S6. C5aR1pp-βarr1/2 complex preparation**.

(A-B) Size exclusion chromatograph and SDS-PAGE gel images of mC5aR1pp-βarr1-Fab30 and mC5aR1pp-βarr2$^{DM}$-Fab30 complexes, respectively.

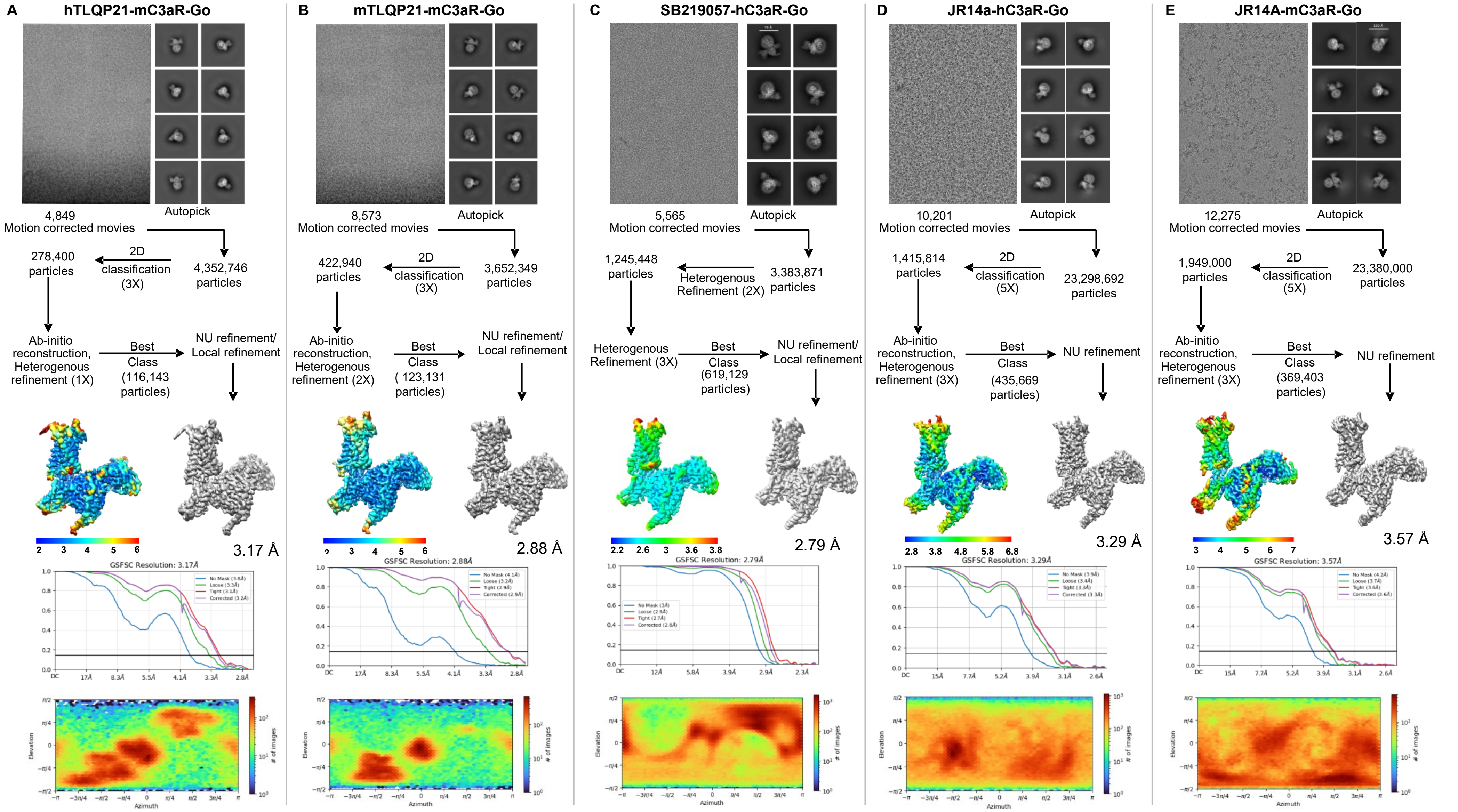
A
hTLQP21-mC3aR-Go
4,849
Motion corrected movies
Autopick
4,352,746
particles
2D
classification
(3X)
278,400
particles
Ab-initio
reconstruction,
Heterogenous
refinement (1X)
Best
Class
(116,143
particles)
NU refinement/
Local refinement
3.17 Å
GSFSC Resolution: 3.17Å
B
mTLQP21-mC3aR-Go
8,573
Motion corrected movies
Autopick
3,652,349
particles
2D
classification
(3X)
422,940
particles
Ab-initio
reconstruction,
Heterogenous
refinement (2X)
Best
Class
( 123,131
particles)
NU refinement/
Local refinement
2.88 Å
GSFSC Resolution: 2.88Å
C
SB219057-hC3aR-Go
5,565
Motion corrected movies
Autopick
3,383,871
particles
Heterogenous
Refinement (2X)
1,245,448
particles
Heterogenous
Refinement (3X)
Best
Class
(619,129
particles)
NU refinement/
Local refinement
2.79 Å
GSFSC Resolution: 2.79Å
D
JR14a-hC3aR-Go
10,201
Motion corrected movies
Autopick
23,298,692
particles
2D
classification
(5X)
1,415,814
particles
Ab-initio
reconstruction,
Heterogenous
refinement (3X)
Best
Class
(435,669
particles)
NU refinement
3.29 Å
GSFSC Resolution: 3.29Å
E
JR14A-mC3aR-Go
12,275
Motion corrected movies
Autopick
23,380,000
particles
2D
classification
(5X)
1,949,000
particles
Ab-initio
reconstruction,
Heterogenous
refinement (3X)
Best
Class
(369,403
particles)
NU refinement
3.57 Å
GSFSC Resolution: 3.57Å
Elevation
Azimuth
### of images

**Figure S7: Workflow for cryo-EM data processing of C3aR complexes.**

(A-E) Representative cryo-EM micrograph, selected 2D class averages representing different orientations, schematic representation of cryo-EM data processing workflow, local resolution map of the 3D reconstruction, gold standard fourier shell correlation curve (GSFSC) at 0.143 threshold, and angular distribution of the particles against the final reconstruction of hTLQP21-mC3aR-Go, mTLQP21-mC3aR-Go, SB219057-hC3aR-Go, JR14a-hC3aR-Go and JR14a-mC3aR-Go complexes, respectively.

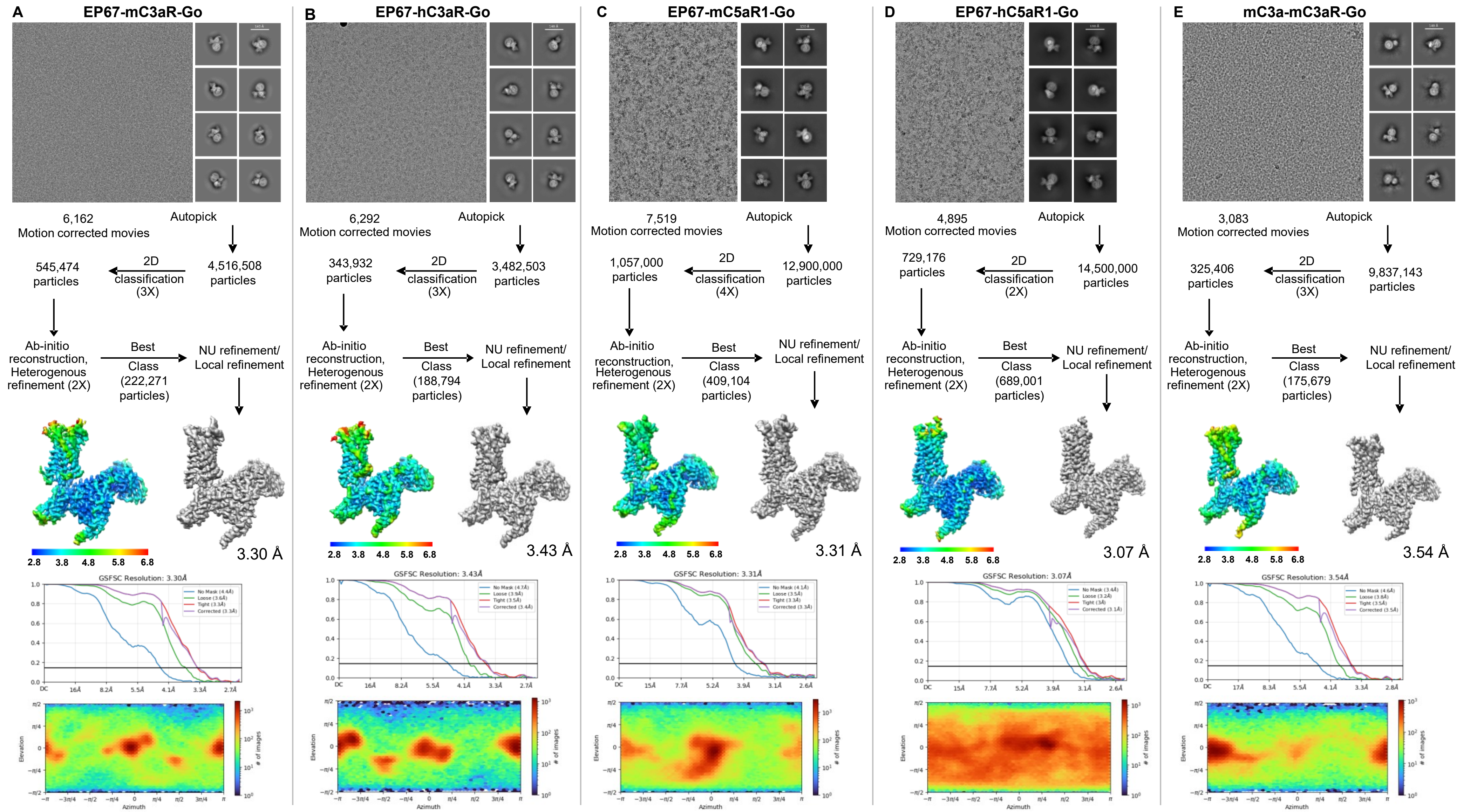

A
EP67-mC3aR-Go
6,162
Motion corrected movies
Autopick
4,516,508
particles
2D
classification
(3X)
545,474
particles
Ab-initio
reconstruction,
Heterogenous
refinement (2X)
Best
Class
(222,271
particles)
NU refinement/
Local refinement
2.8 3.8 4.8 5.8 6.8
3.30 Å
GSFSC Resolution: 3.30Å
B
EP67-hC3aR-Go
6,292
Motion corrected movies
Autopick
3,482,503
particles
2D
classification
(3X)
343,932
particles
Ab-initio
reconstruction,
Heterogenous
refinement (2X)
Best
Class
(188,794
particles)
NU refinement/
Local refinement
2.8 3.8 4.8 5.8 6.8
3.43 Å
GSFSC Resolution: 3.43Å
C
EP67-mC5aR1-Go
7,519
Motion corrected movies
Autopick
12,900,000
particles
2D
classification
(4X)
1,057,000
particles
Ab-initio
reconstruction,
Heterogenous
refinement (2X)
Best
Class
(409,104
particles)
NU refinement/
Local refinement
2.8 3.8 4.8 5.8 6.8
3.31 Å
GSFSC Resolution: 3.31Å
D
EP67-hC5aR1-Go
4,895
Motion corrected movies
Autopick
14,500,000
particles
2D
classification
(2X)
729,176
particles
Ab-initio
reconstruction,
Heterogenous
refinement (2X)
Best
Class
(689,001
particles)
NU refinement/
Local refinement
2.8 3.8 4.8 5.8 6.8
3.07 Å
GSFSC Resolution: 3.07Å
E
mC3a-mC3aR-Go
3,083
Motion corrected movies
Autopick
9,837,143
particles
2D
classification
(3X)
325,406
particles
Ab-initio
reconstruction,
Heterogenous
refinement (2X)
Best
Class
(175,679
particles)
NU refinement/
Local refinement
2.8 3.8 4.8 5.8 6.8
3.54 Å
GSFSC Resolution: 3.54Å
Elevation
Azimuth
### of images

**Figure S8: Workflow for cryo-EM data processing of C3aR and C5aR1 complexes.**

(A-E) Representative cryo-EM micrograph, selected 2D class averages representing different orientations, schematic representation of cryo-EM data processing workflow, local resolution map of the 3D reconstruction, gold standard fourier shell correlation curve (GSFSC) at 0.143 threshold, and angular distribution of the particles against the final reconstruction of EP67-mC3aR-Go, EP67-hC3aR-Go, EP67-mC5aR1-Go, EP67-hC5aR1-Go and mC3a-mC3aR-Go complexes, respectively.

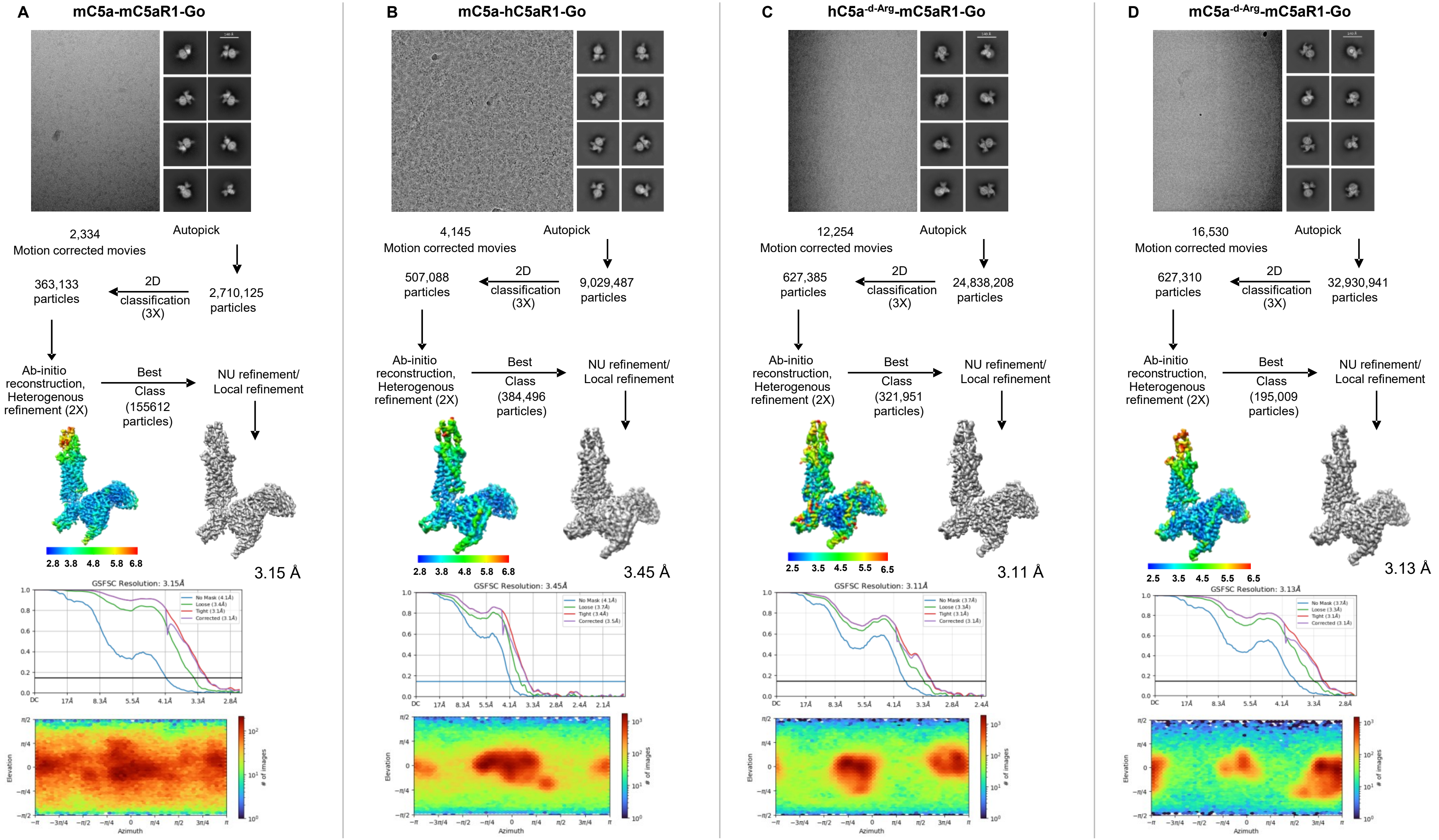

A mC5a-mC5aR1-Go
2,334
Motion corrected movies
Autopick
2,710,125 particles
2D classification (3X)
363,133 particles
Ab-initio reconstruction, Heterogenous refinement (2X)
Best Class (155612 particles)
NU refinement/ Local refinement
2.8 3.8 4.8 5.8 6.8
3.15 Å
GSFSC Resolution: 3.15Å
B mC5a-hC5aR1-Go
4,145
Motion corrected movies
Autopick
9,029,487 particles
2D classification (3X)
507,088 particles
Ab-initio reconstruction, Heterogenous refinement (2X)
Best Class (384,496 particles)
NU refinement/ Local refinement
2.8 3.8 4.8 5.8 6.8
3.45 Å
GSFSC Resolution: 3.45Å
C hC5a-d-Arg-mC5aR1-Go
12,254
Motion corrected movies
Autopick
24,838,208 particles
2D classification (3X)
627,385 particles
Ab-initio reconstruction, Heterogenous refinement (2X)
Best Class (321,951 particles)
NU refinement/ Local refinement
2.5 3.5 4.5 5.5 6.5
3.11 Å
GSFSC Resolution: 3.11Å
D mC5a-d-Arg-mC5aR1-Go
16,530
Motion corrected movies
Autopick
32,930,941 particles
2D classification (3X)
627,310 particles
Ab-initio reconstruction, Heterogenous refinement (2X)
Best Class (195,009 particles)
NU refinement/ Local refinement
2.5 3.5 4.5 5.5 6.5
3.13 Å
GSFSC Resolution: 3.13Å
Elevation
Azimuth
### of images

**Figure S9: Workflow for cryo-EM data processing of C5aR1 complexes.**

(A-D) Representative cryo-EM micrograph, selected 2D class averages representing different orientations, schematic representation of cryo-EM data processing workflow, local resolution map of the 3D reconstruction, gold standard fourier shell correlation curve (GSFSC) at 0.143 threshold, and angular distribution of the particles against the final reconstruction of mC5a-mC5aR1-Go, mC5a-hC5aR1-Go, hC5a$^{\text{-d-Arg}}$-mC5aR1-Go and mC5a$^{\text{-d-Arg}}$-mC5aR1-Go complexes, respectively.

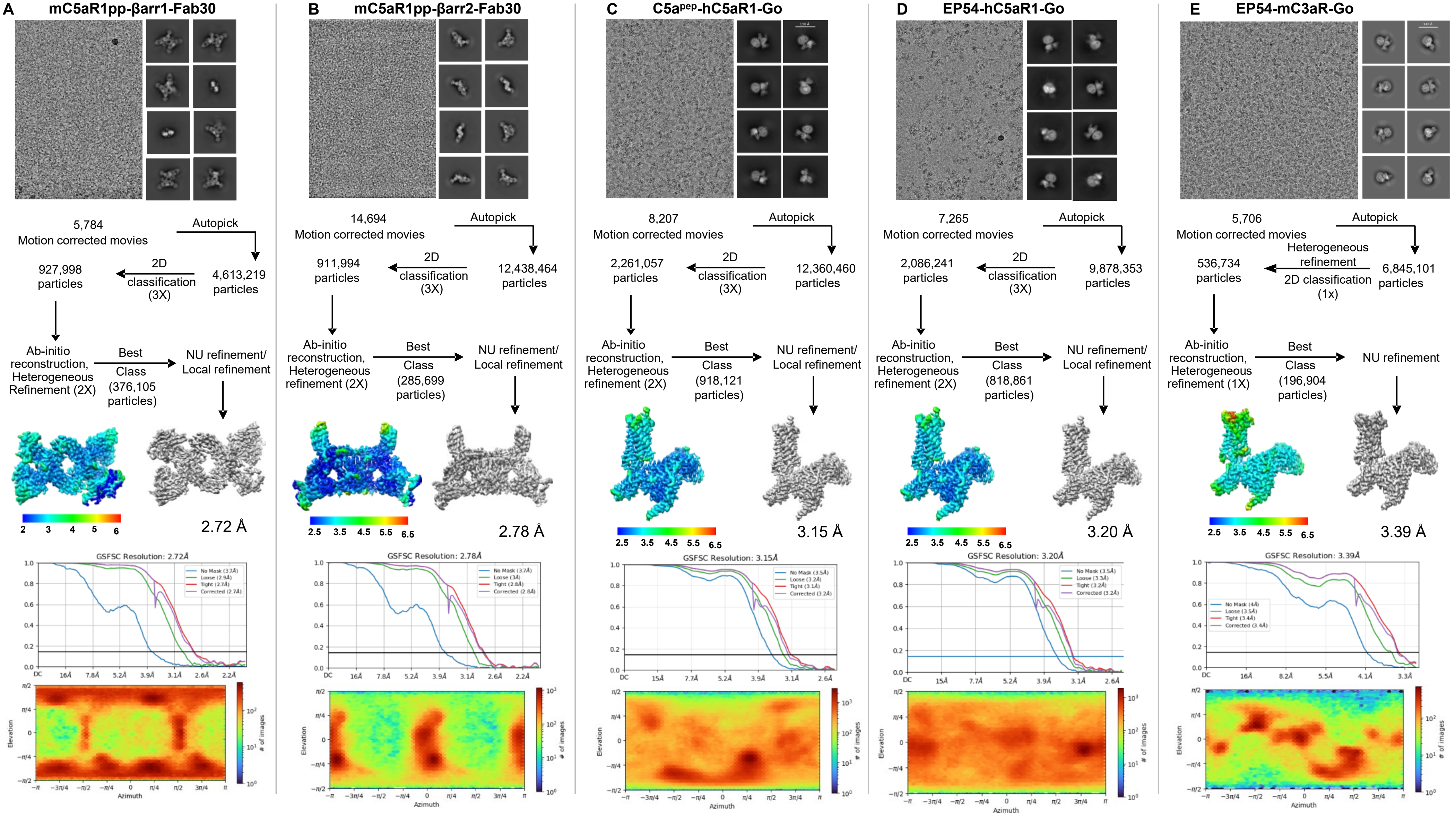
A mC5aR1pp-βarr1-Fab30
5,784
Motion corrected movies
Autopick
4,613,219 particles
2D classification (3X)
927,998 particles
Ab-initio reconstruction, Heterogeneous Refinement (2X)
Best Class (376,105 particles)
NU refinement/ Local refinement
2.72 Å
GSFSC Resolution: 2.72Å
B mC5aR1pp-βarr2-Fab30
14,694
Motion corrected movies
Autopick
12,438,464 particles
2D classification (3X)
911,994 particles
Ab-initio reconstruction, Heterogeneous refinement (2X)
Best Class (285,699 particles)
NU refinement/ Local refinement
2.78 Å
GSFSC Resolution: 2.78Å
C C5apep-hC5aR1-Go
8,207
Motion corrected movies
Autopick
12,360,460 particles
2D classification (3X)
2,261,057 particles
Ab-initio reconstruction, Heterogeneous refinement (2X)
Best Class (918,121 particles)
NU refinement/ Local refinement
3.15 Å
GSFSC Resolution: 3.15Å
D EP54-hC5aR1-Go
7,265
Motion corrected movies
Autopick
9,878,353 particles
2D classification (3X)
2,086,241 particles
Ab-initio reconstruction, Heterogeneous refinement (2X)
Best Class (818,861 particles)
NU refinement/ Local refinement
3.20 Å
GSFSC Resolution: 3.20Å
E EP54-mC3aR-Go
5,706
Motion corrected movies
Autopick
6,845,101 particles
Heterogeneous refinement
2D classification (1x)
536,734 particles
Ab-initio reconstruction, Heterogeneous refinement (1X)
Best Class (196,904 particles)
NU refinement
3.39 Å
GSFSC Resolution: 3.39Å
Elevation
Azimuth
### of images

**Figure S10: Workflow for cryo-EM data processing of βarr-phosphopeptide complexes and C3aR and C5aR1 complexes.**

(A-E) Representative cryo-EM micrograph, selected 2D class averages representing different orientations, schematic representation of cryo-EM data processing workflow, local resolution map of the 3D reconstruction, gold standard fourier shell correlation curve (GSFSC) at 0.143 threshold, and angular distribution of the particles against the final reconstruction of mC5aR1pp-βarr1-Fab30, mC5aR1pp-βarr2-Fab30, $C5a^{pep}$-hC5aR1-Go, EP54-hC5aR1-Go and EP54-mC3aR-Go complexes, respectively.

Figure S11

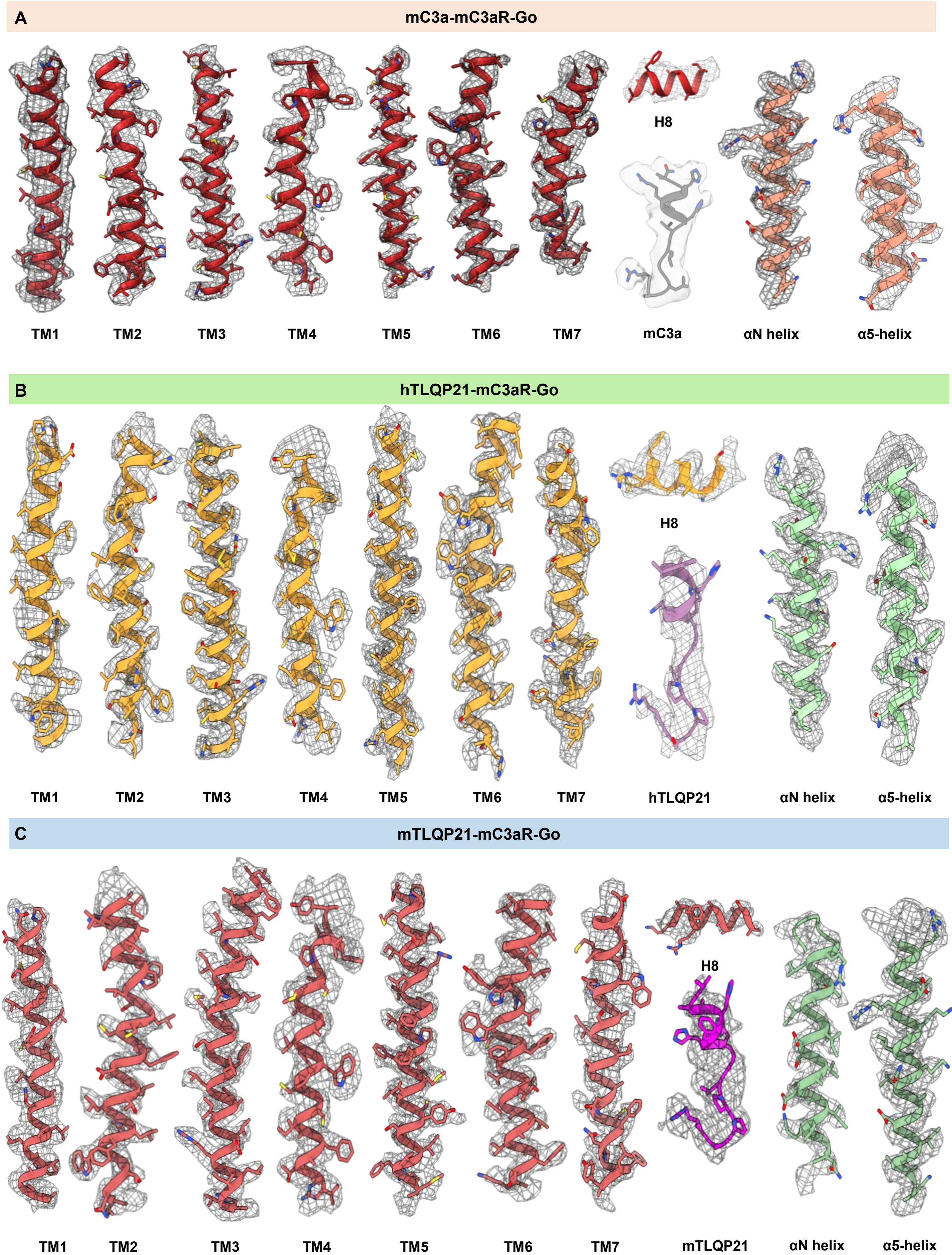


**Figure S11. Representative electron density maps**.

(A-C) EM densities of TM1 to TM7, helix-8, α5-helix, αN helix, and ligands of mC3a-mC3aR-Go, hTLQp21-mC3aR-Go, mTLQP21-mC3aR-Go complexes respectively.

**Figure S12**

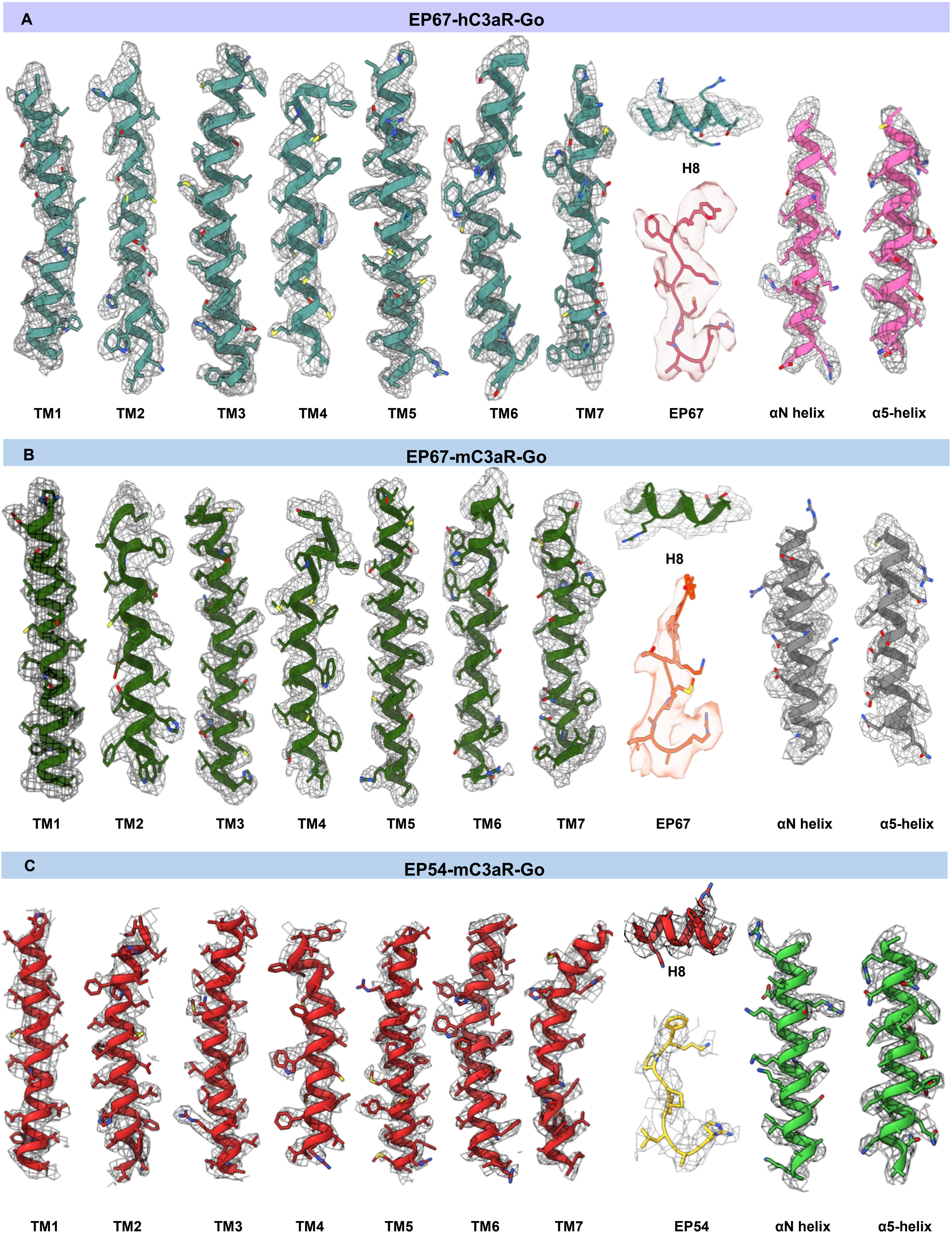


**Figure S12. Representative electron density maps.**

(A-C) EM densities of TM1 to TM7, helix 8, α5-helix, αN helix, and ligands of EP67-hC3aR-Go, EP67-mC3aR-Go and EP54-mC3aR-Go complexes respectively.

Figure S13

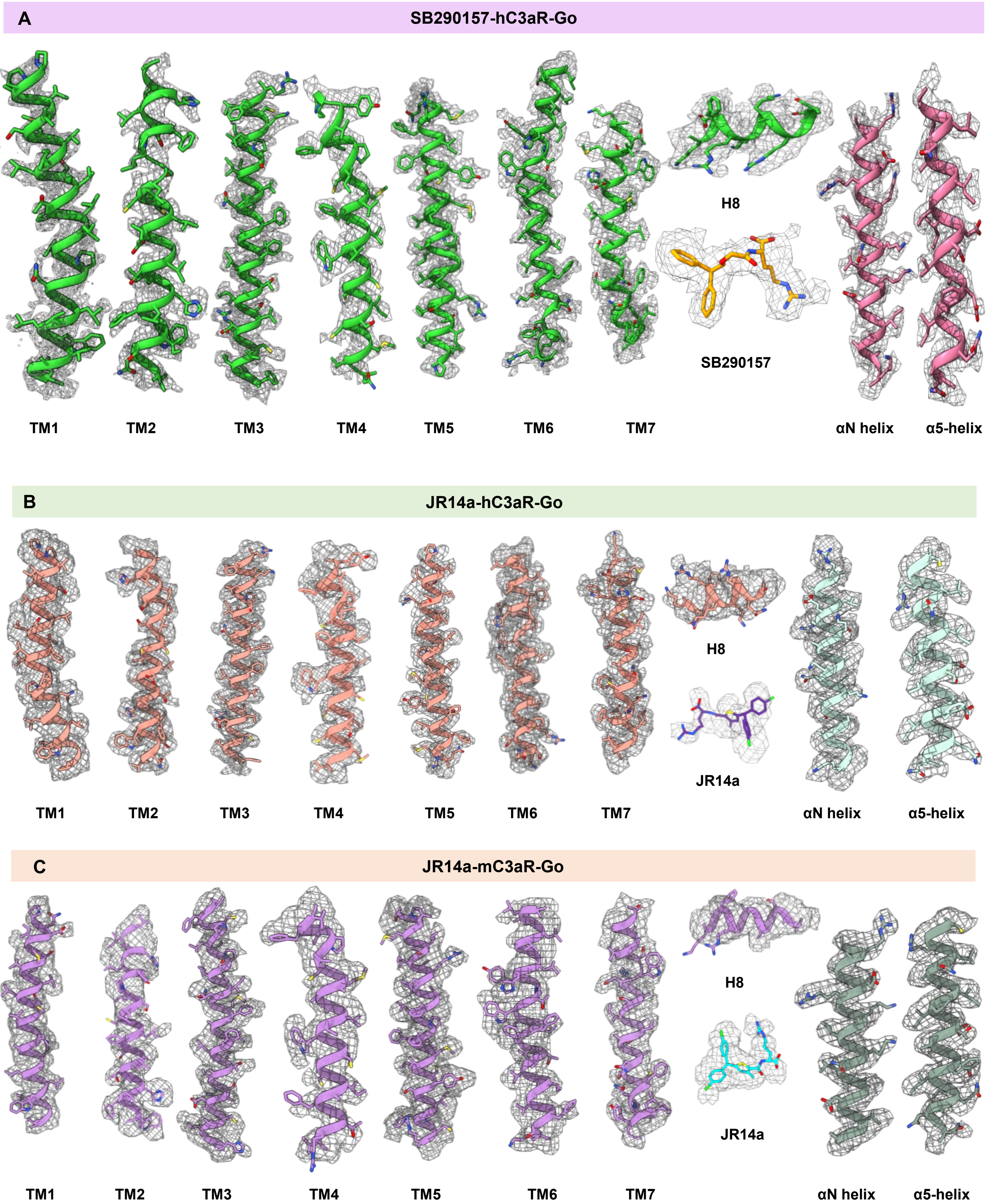


**Figure S13**. **Representative electron density maps**.

(A-C) EM densities of TM1 to TM7, helix 8, α5-helix, αN helix, and ligands SB290157-hC3aR-Go, JR14a-hC3aR-Go, JR14a-mC3aR-Go complexes respectively.

**Figure S14**

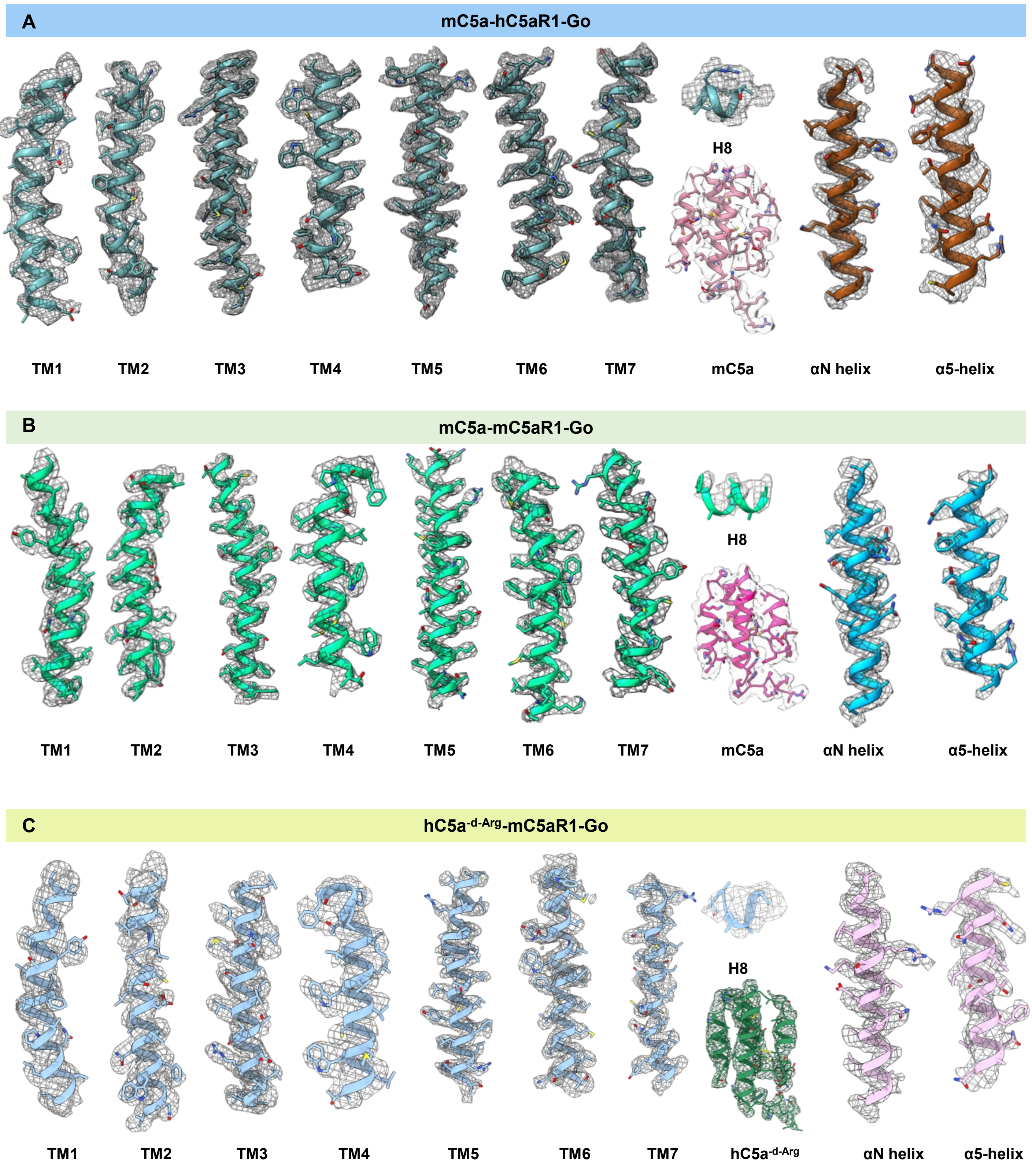


**Figure S14. Representative electron density maps.**

(A-C) EM densities of TM1 to TM7, helix 8, α5-helix, αN helix, and ligands of mC5a-hC5aR1-Go, mC5a-mC5aR1-Go hC5a$^{\text{-d-Arg}}$-mC5aR1-Go, complexes respectively.

**Figure S15**

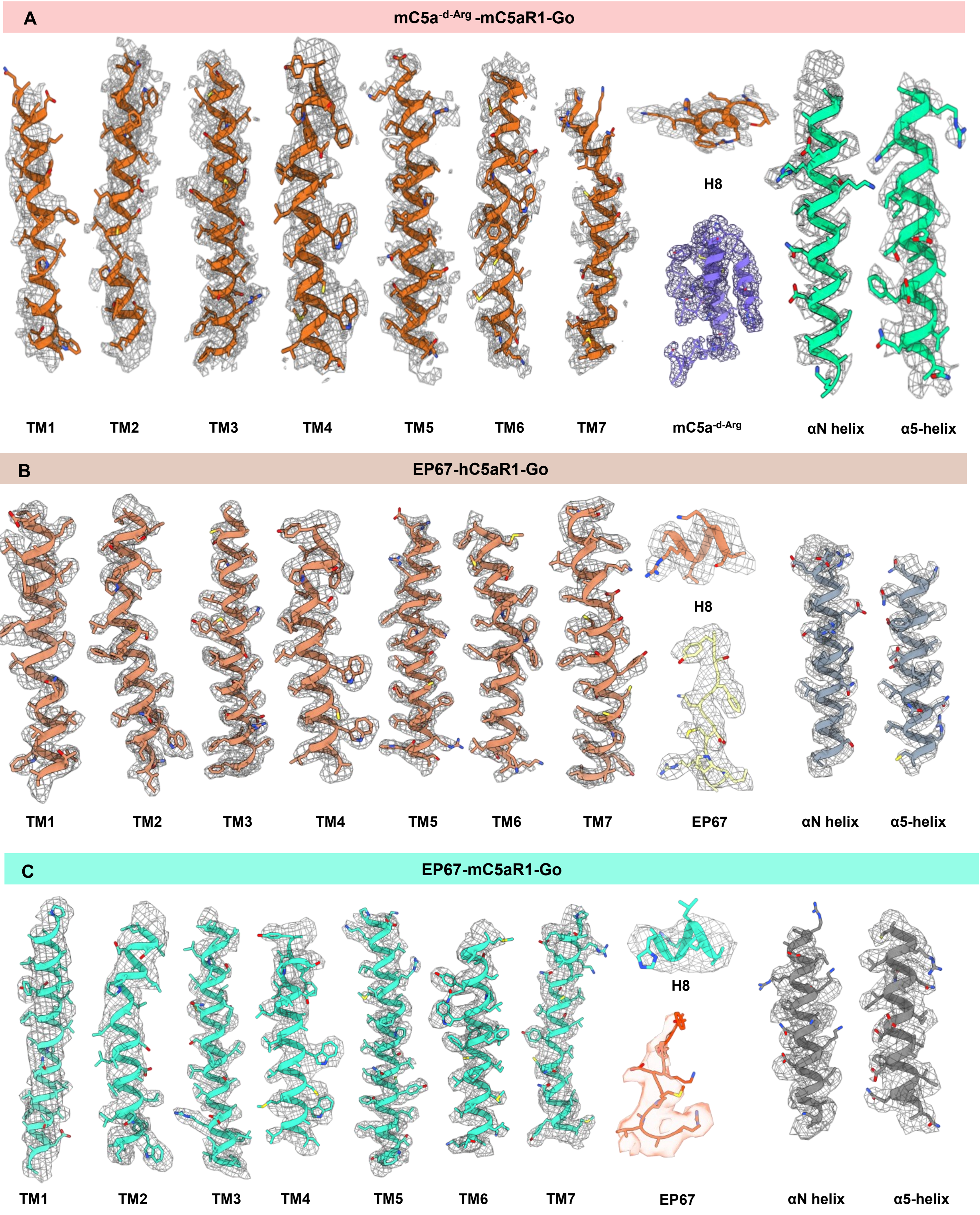


**Figure S15 Representative electron density maps.**

(A-C) EM densities of TM1 to TM7, helix 8, α5-helix, αN helix, and ligands of mC5a$^{-d-Arg}$-mC5aR1-Go, EP67-hC5aR1-Go, EP67-mC5aR1-Go complexes respectively.

**Figure S16**

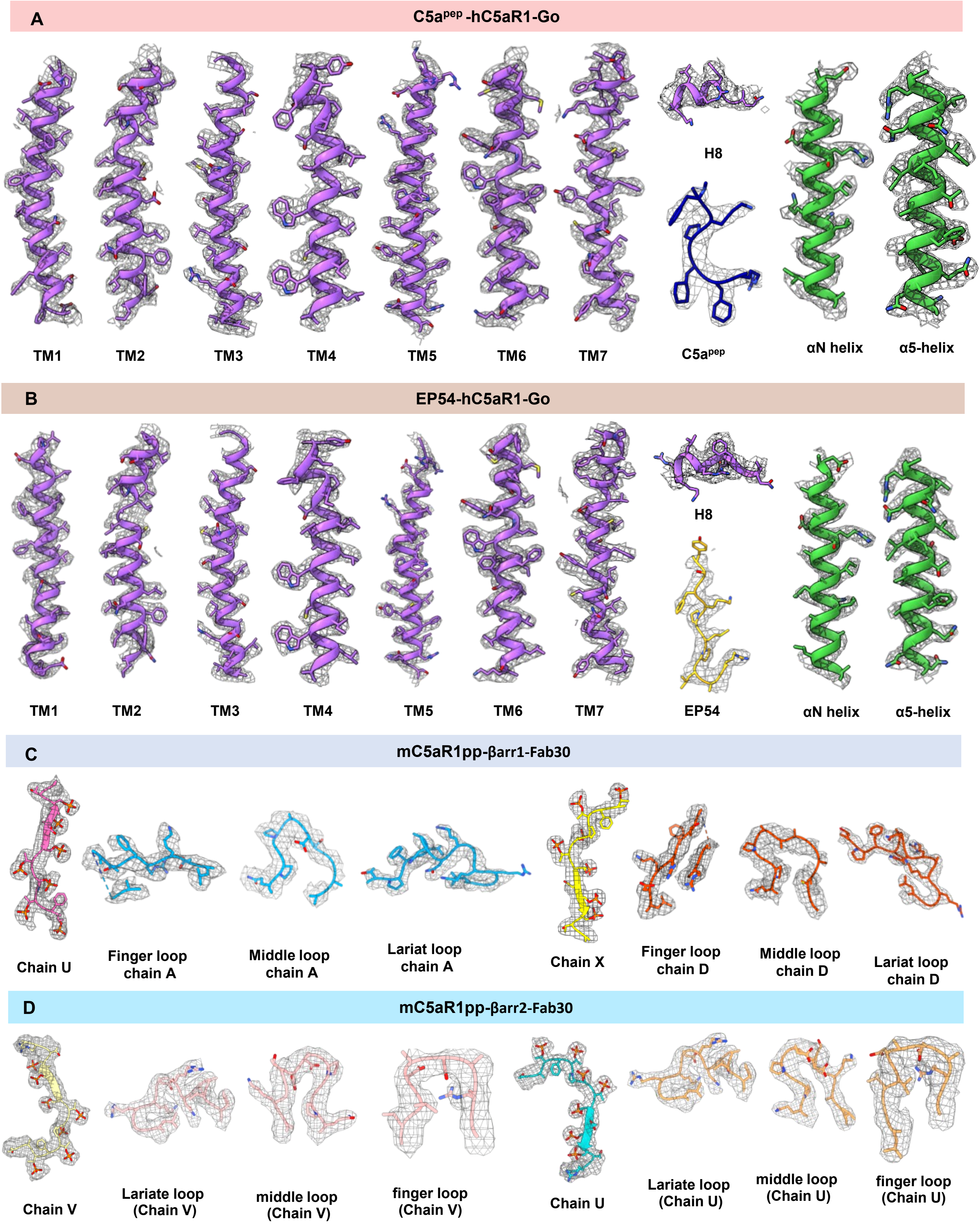


**Figure S16 Representative electron density maps.**
(A-B) EM densities of TM1 to TM7, helix 8, α5-helix, αN helix, and ligands of C5a$^{pep}$-hC5aR1-Go and EP54-hC5aR1-Go, complexes respectively; **C-D**. EM densities of phosphopeptides and key loop regions of mC5aR1pp-βarr1-Fab30 and mC5aR1pp-βarr2-Fab30 complexes respectively.

**Figure S17**

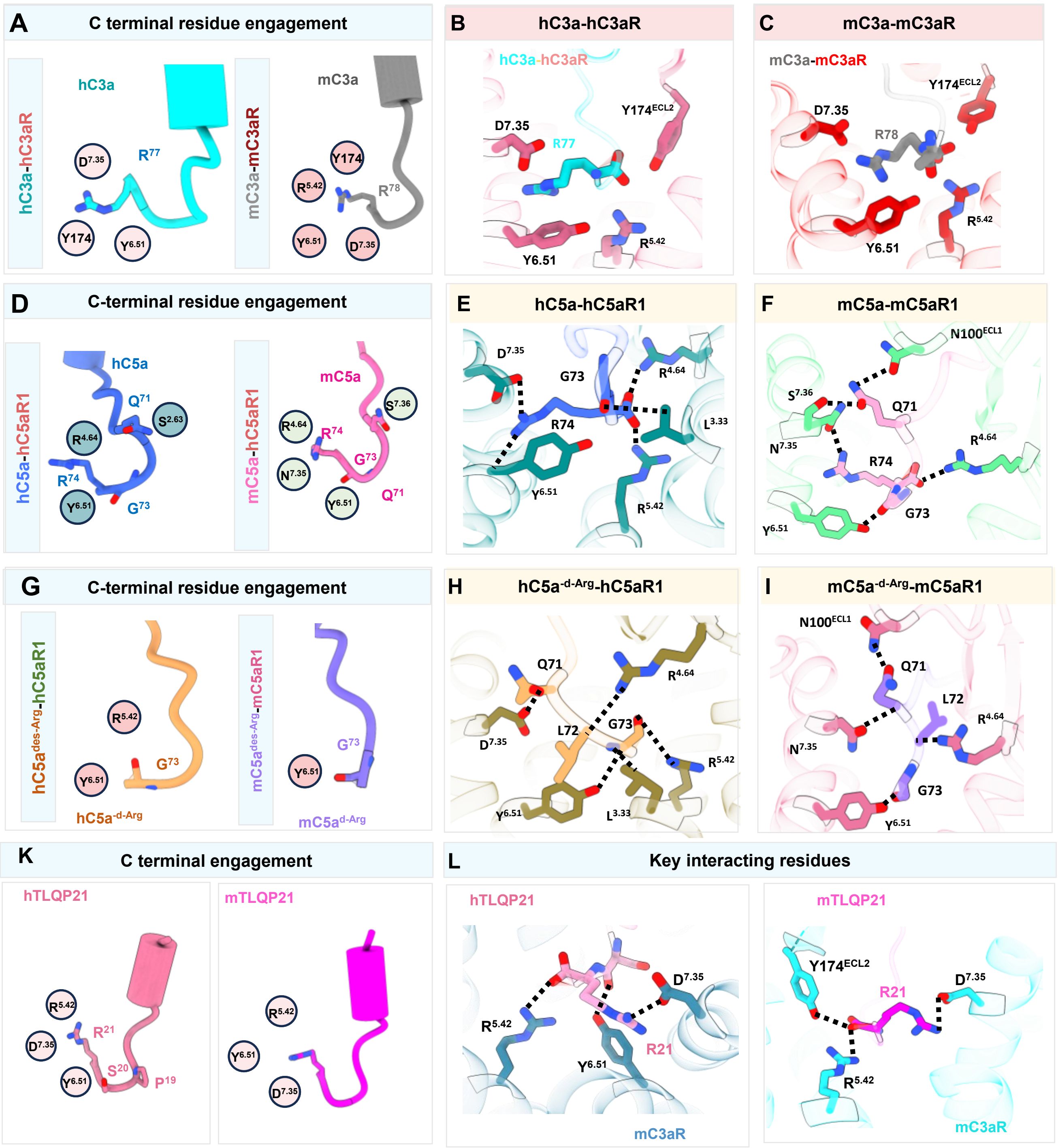


**Figure S17. C-terminal engagement of various ligands in the orthosteric pocket of the receptor**.

(A-F) Comparison of the terminal arginine residue interactions within the orthosteric binding pocket of hC3a-hC3aR vs. mC3a-mC3aR and hC5a-hC5aR1 vs. mC5a-mC5aR1.

(G-I) Interaction of G73 residue within the orthosteric binding pocket of hC5a$^{\text{-d-Arg}}$-hC5aR1 and mC5a$^{\text{-d-Arg}}$-mC5aR1, compensating for the absence of the terminal arginine residue interaction.

(K-L) Key ligand-binding interactions in the orthosteric binding site of hTLQP21 and mTLQP21 with mC3aR, detailing the molecular interactions and structural determinants of ligand specificity

**Figure S18**

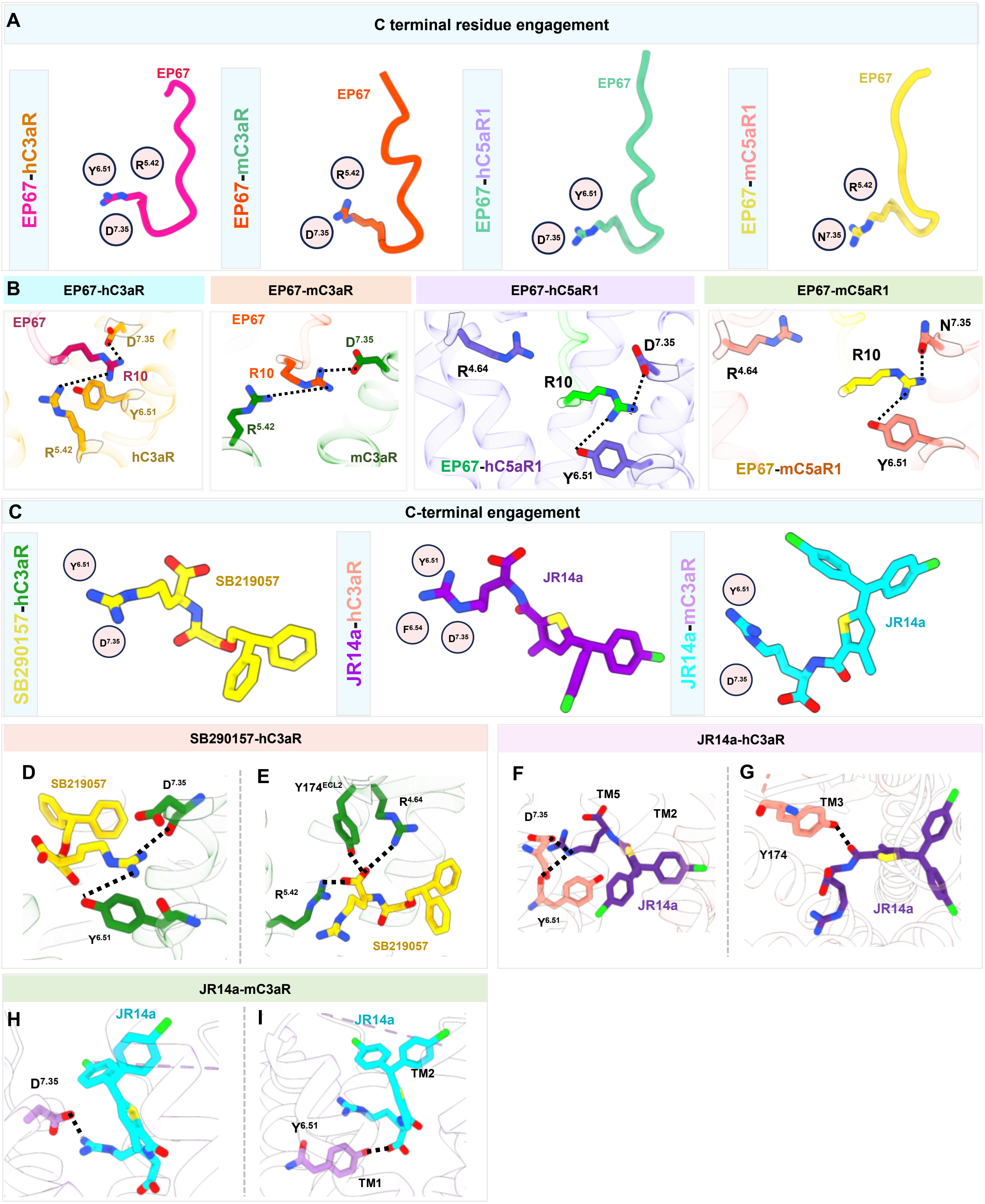


**Figure S18. C-terminal engagement of synthetic peptide EP67 and small molecule agonist SB290157 and JR14a within the orthosteric pocket of the complement receptor.**

(A-B) Key ligand-binding interactions of EP67 with hC3aR, mC3aR, hC5aR1, and mC5aR1, illustrating critical contacts that promote stable binding and receptor activation,

(C-I) Comparison of the terminal arginine residue interactions within the orthosteric binding pocket SB290157-hC3aR, JR14a-hC3aR, JR14a-mC3aR, emphasising structural variation in receptor-ligand interactions. Snapshots focusing on the guanidinium group interactions within the orthosteric binding pocket for SB290157-hC3aR, JR14a-hC3aR, and JR14a-mC3aR structures, respectively.

Figure S19

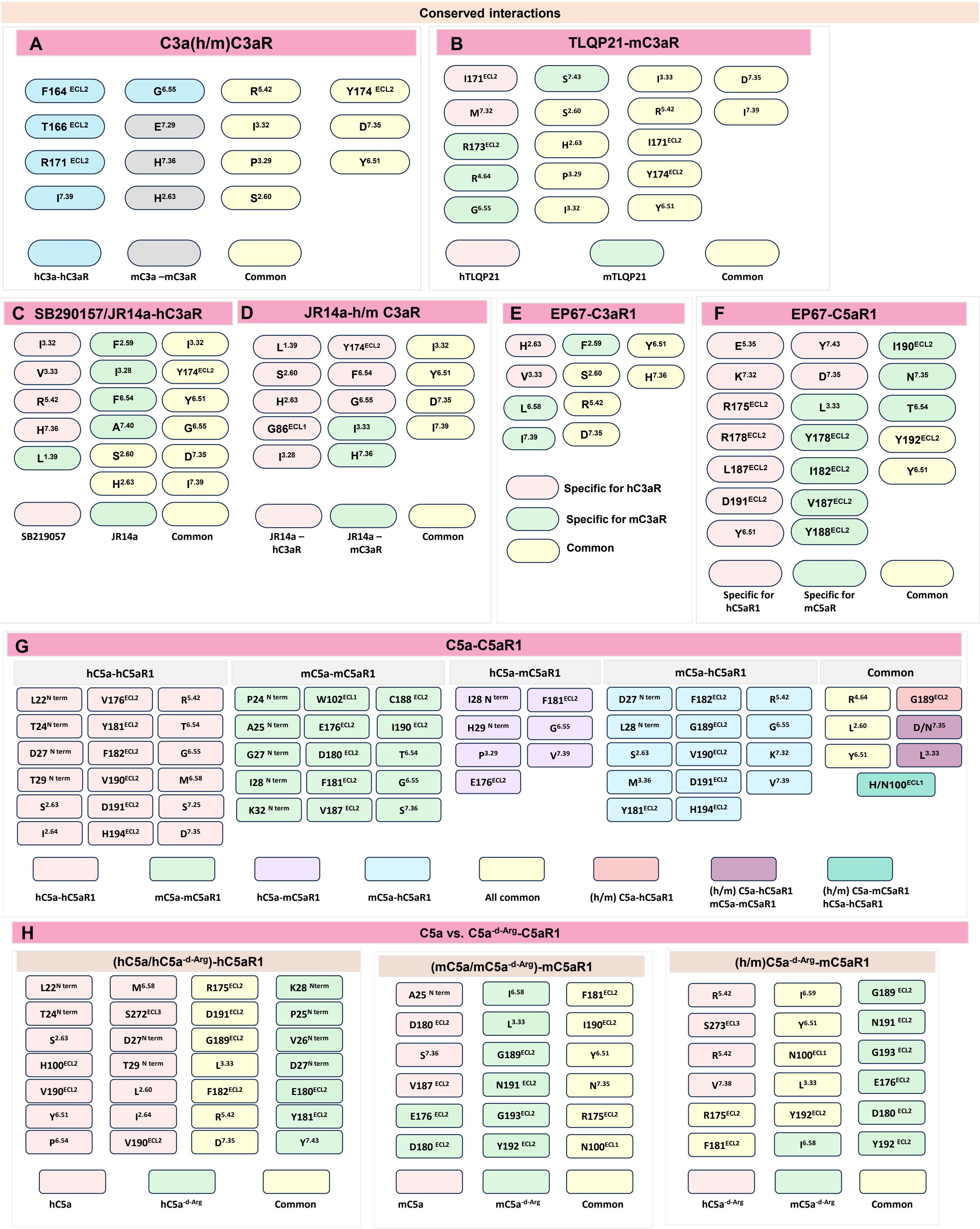


**Figure S19 List of conserved ligand-interacting residues within orthosteric pocket of complement receptor** (A-H) List of conserved ligand-interacting residues between C3a-C3aR, TLQP21-C3aR, SB290157 vs. JR14a-hC3aR, JR14a-hC3aR/mC3aR, EP67-C3aR, EP67-C5aR1, C5a-C5aR1, C5a$^{-d-Arg}$-C5aR1 respectively.

#### Figure S20

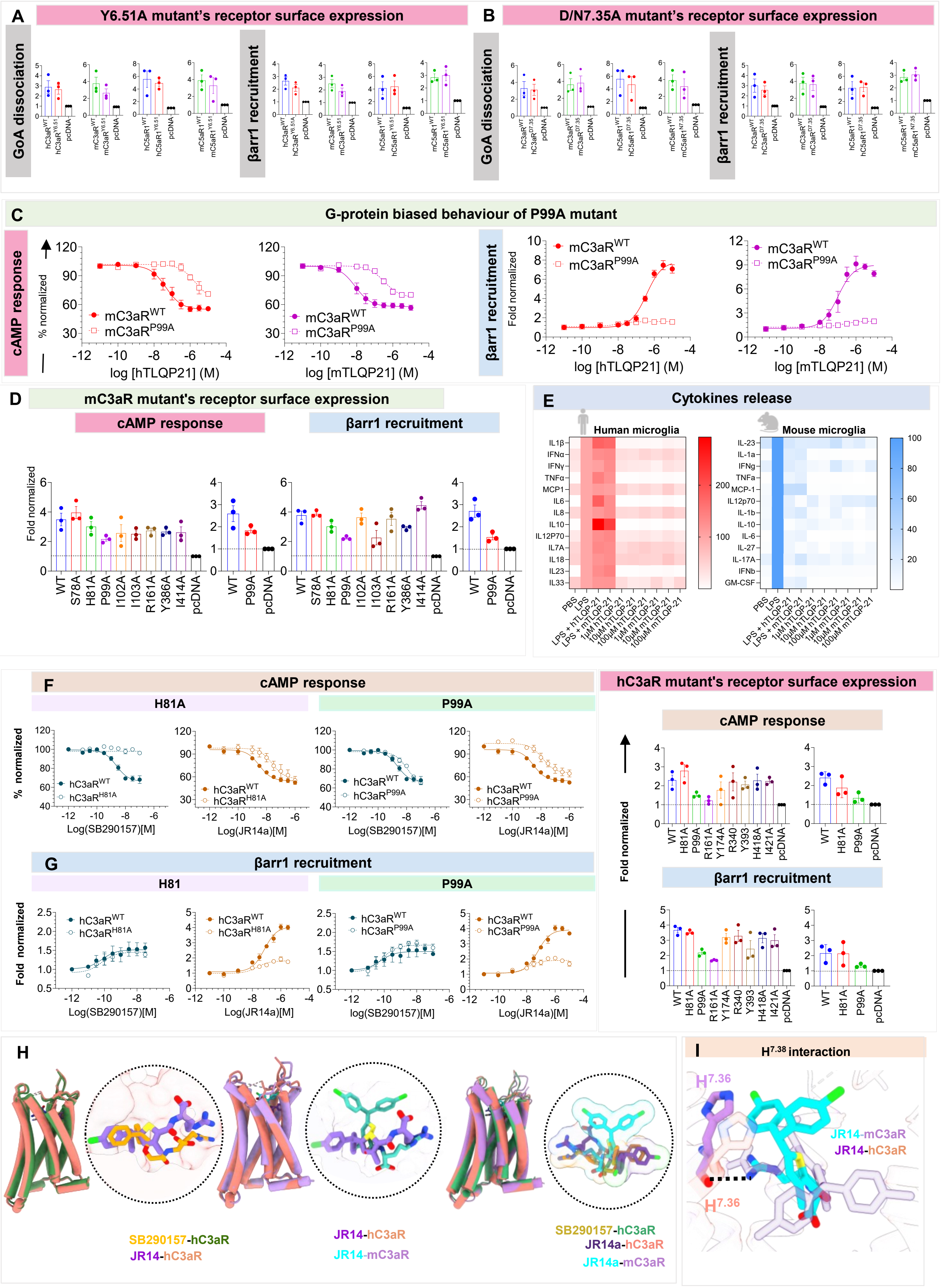
A
Y6.51A mutant's receptor surface expression
GoA dissociation
βarr1 recruitment
B
D/N7.35A mutant's receptor surface expression
C
G-protein biased behaviour of P99A mutant
cAMP response
% normalized
mC3aR^WT
mC3aR^P99A
log [hTLQP21] (M)
log [mTLQP21] (M)
Fold normalized
D
mC3aR mutant's receptor surface expression
WT
S78A
H81A
P99A
I102A
I103A
R161A
Y386A
I414A
pcDNA
E
Cytokines release
Human microglia
Mouse microglia
PBS
LPS
LPS + hTLQP-21
LPS + mTLQP-21
1µM hTLQP-21
10µM hTLQP-21
100µM hTLQP-21
1µM mTLQP-21
10µM mTLQP-21
100µM mTLQP-21
F
H81A
hC3aR^WT
hC3aR^H81A
hC3aR^P99A
Log(SB290157)[M]
Log(JR14a)[M]
hC3aR mutant's receptor surface expression
Y174A
R340
Y393
H418A
I421A
G
H81
H
SB290157-hC3aR
JR14-hC3aR
JR14-mC3aR
JR14a-hC3aR
JR14a-mC3aR
I
H7.38 interaction
H7.36

**Figure S20. Functional characterization of a structure guided engineered G-protein biased mutant in C3aR**

(A) Surface expression of C3aR and C5aR1 WT and mutant $Y^{6.51A}$ (conserved residue) measured using whole-cell ELISA. Data are presented as mean ± SEM (n = 3), normalised to mock-transfected cells (considered 1), corresponding to figure 4 (C and D).

(B) Surface expression of C3aR and C5aR1 WT and mutant $D/N^{7.35A}$ (conserved residue) measured using whole-cell ELISA. Data are presented as mean ± SEM (n = 3), normalised to mock-transfected cells (considered 1), corresponding to figure 4 (D and E).

(C) Dose-dependent TLQP21-induced cAMP and βarr1 recruitment responses measured using the GloSensor assay and NanoBiT-based complementation assay (bystander mode) downstream of mC3aR WT and $P^{99A}$ mutant, respectively. Data are presented as mean ± SEM (n = 3); for GloSensor assay, normalized to the lowest ligand concentration (considered 100%) after forskolin correction, and for βarr1, normalized to the lowest ligand concentration (considered 1).

(D) Surface expression of mC3aR WT and mutants (selected based on critical interacting residues with human and mouse TLQP21) measured using whole-cell ELISA. Data are presented as mean ± SEM (n = 3), normalised to mock-transfected cells (considered 1)

(E) Heatmap indicating cytokine release in human and mouse primary microglial cells treated with PBS, LPS, and different doses of human and mouse TLQP21. Data are presented as mean ± SEM (n = 4), normalised to LPS (considered 100%).

(F and G) The critical residues H81 and P99 in hC3aR show biased transducer selectivity upon stimulation with JR14a but not with SB290157. **F** Ligand-mediated Gαi activation was quantified using the GloSensor assay, where a dose-dependent concentration of SB290157 and JR14a reduces forskolin-induced cAMP levels downstream of WT and mutants H81A and P99A were measured(data presented as mean ± SEM n=3; normalised with the lowest ligand concentration considered 100% after forskolin values correction). **G** βarr1 recruitment was studied via NanoBiT-based complementation assay (bystander mode) in a dose-dependent manner (Receptor_pcDNA + SmBiT_βarr1 + LgBiT_CAAX) (mean ± SEM n=3; normalised with the lowest ligand concentration considered 1). Respective surface expression of hC3aR WT and mutants in GloSensor assay and NanoBiT assay measured using whole cell-based ELISA technique ( mean ± SEM n=3; normalised with mock-transfected cells considered 1).

(H) Structural superimposition of SB290157-hC3aR with JR14a-hC3aR, JR14a-hC3aR with JR14a-mC3aR, and SB290157-hC3aR, JR14a-hC3aR, and JR14a-mC3aR combined. The ligand binding poses are highlighted in the upper right corner, emphasizing key structural alignments.

(I) Interaction of H4.11 in JR14a-mC3aR to stabilize the ligand within the orthosteric pocket of mC3aR, whereas this interaction is absent in JR14a-hC3aR.

#### Figure S21

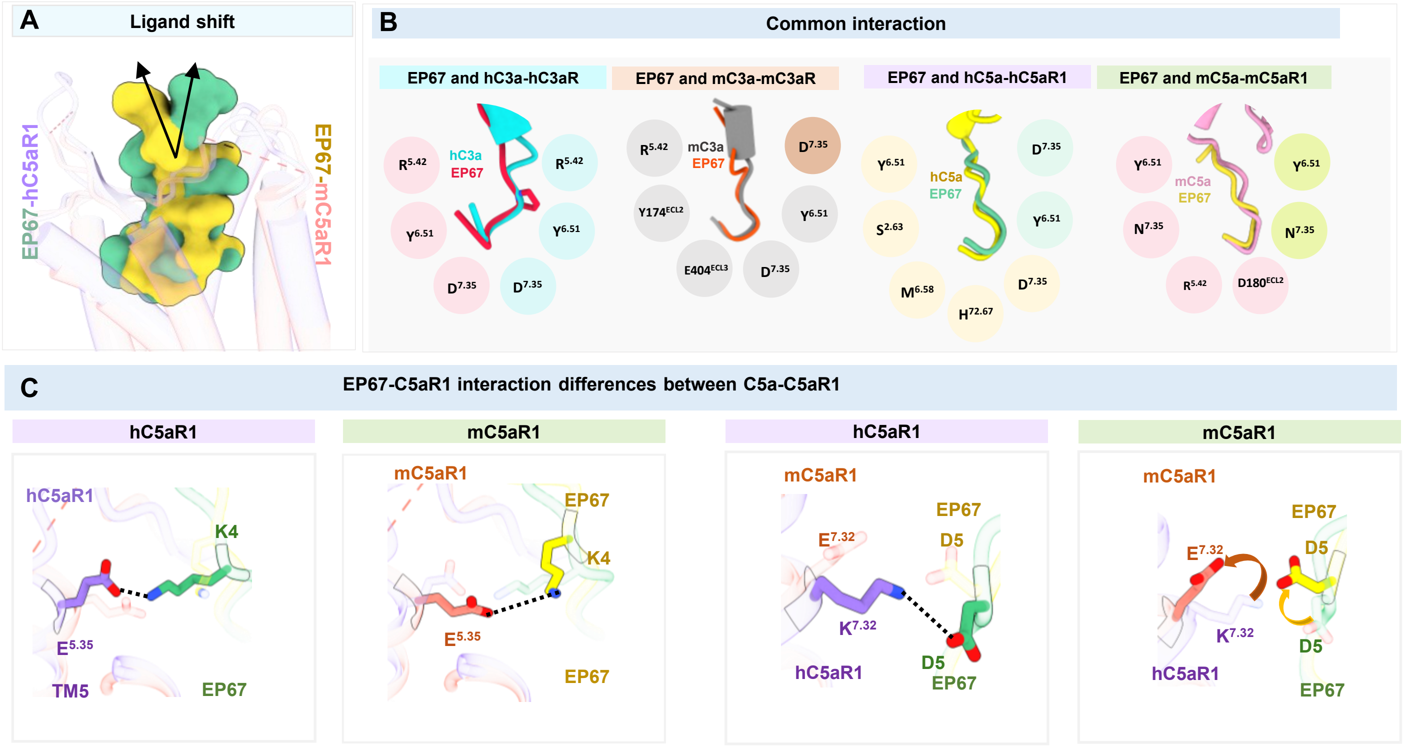

A
Ligand shift
EP67-hC5aR1
EP67-mC5aR1
B
Common interaction
EP67 and hC3a-hC3aR
EP67 and mC3a-mC3aR
EP67 and hC5a-hC5aR1
EP67 and mC5a-mC5aR1
hC3a
EP67
mC3a
EP67
hC5a
EP67
mC5a
EP67
C
EP67-C5aR1 interaction differences between C5a-C5aR1
hC5aR1
mC5aR1
hC5aR1
mC5aR1

**Figure S21. Comparative structural insights into ligand binding dynamics for EP67 and C5adesArg with their respective complement receptors**

(A) Comparison of the EP67 binding to hC5aR1 and mC5aR1 highlights differences in ligand orientation between the two receptor-ligand complexes.

(B) Schematic representation highlighting terminal loop interactions of the ligand EP67 within the orthosteric binding pocket of hC3aR, mC3aR, hC5aR1, and mC5aR1. The interactions are compared with those of endogenous ligands (hC3a, mC3a, hC5a, and mC5a), emphasizing conserved and differential binding patterns among the receptors.

(C) Comparison of key ligand-binding interactions of EP67 with hC5aR1, illustrating critical contacts that facilitate stable binding and receptor activation.
